# An updated genome-scale metabolic network reconstruction of *Pseudomonas aeruginosa* PA14 to characterize mucin-driven shifts in bacterial metabolism

**DOI:** 10.1101/2021.03.10.434463

**Authors:** Dawson D. Payne, Alina Renz, Laura J. Dunphy, Taylor Lewis, Andreas Dräger, Jason A. Papin

## Abstract

Mucins are present in mucosal membranes throughout the body and play a key role in the microbe clearance and infection prevention. Understanding the metabolic responses of pathogens to mucins will further enable the development of protective approaches against infections. We update the genome-scale metabolic network reconstruction (GENRE) of one such pathogen, *Pseudomonas aeruginosa* PA14, through metabolic coverage expansion, format update, extensive annotation addition, and literature-based curation to produce iPau21. We then validate iPau21 through MEMOTE, growth rate, carbon source utilization, and gene essentiality testing to demonstrate its improved quality and predictive capabilities. We then integrate the GENRE with transcriptomic data in order to generate context-specific models of *P. aeruginosa* metabolism. The contextualized models recapitulated known phenotypes of unaltered growth and a differential utilization of fumarate metabolism, while also revealing an increased utilization of propionate metabolism upon MUC5B exposure. This work serves to validate iPau21 and demonstrate its utility for providing biological insights.

## Introduction

The mucosal barrier is a hydrated mucus gel that lines wet epithelial cells throughout the body, including eyes, mouth, lungs, and the gastrointestinal and urogenital tracts^1,2^. It serves as a key mechanism of protection against pathogens. The component responsible for the gel-like properties of the mucosal layer is the glycoprotein mucin^3^. The dysregulation of mucins underlies diseases like cystic fibrosis^4^ and chronic obstructive pulmonary disorders^2^. As mucins are involved in the clearance of microbes^5^, a dysregulation of mucins can result in pathogen overgrowth and severe infections^6^. While some bacterial species, including pathogenic strains of *Pseudomonas*^3^, are capable of residing within the mucosal layer, mucins typically impair the formation of biofilms and surface attachment^7^. Furthermore, mucins are reported to downregulate virulence genes involved in siderophore biosynthesis, quorum sensing, and toxin secretion^1^. By disturbing these key mechanisms of infection, mucins attenuate the virulence and infective potential of *P. aeruginosa*.

Elucidating the metabolic responses of *P. aeruginosa* to mucins can enable the development of protective approaches against infection^8^. Genome-scale metabolic network reconstructions (GENREs) and associated genome-scale metabolic models (GEMs) are well suited for this purpose as they can enable the prediction of cellular behavior under different biological conditions such as the absence or presence of different mucins in an environment^9^. A GENRE can also be used to contextualize high-throughput data, such as transcriptomics or proteomics data^10^. Gene expression data can, for example, be used to constrain specific predicted metabolic fluxes^11^ and thereby increase the predictive value of the model. Metabolically active pathways under different conditions can be identified by integrating high-throughput data with a metabolic network^12^.

*P. aeruginosa* is a critical bacterial species in the ‘Priority Pathogens List’ for research and development of new antibiotics published by the World Health Organization (WHO)^13^. However, the lack of novel antibiotics^14,15^ emphasizes the need for the development of innovative and protective therapeutic approaches. This pressing need for protective strategies coupled with new insights from recent research present an opportunity to further refine the GENRE of the highly virulent strain UCBPP-PA14 by Bartell *et al*.^16^. An updated GENRE can be used to better understand the metabolic underpinnings of *P. aeruginosa* infections and ultimately develop new therapeutic strategies from those insights.

Here, we present iPau21, an updated GENRE of *P. aeruginosa* strain UCBPP-PA14 metabolism. We improve predictions of carbon source utilization and growth yields in order to better recapitulate the behavior of the pathogen. Metabolic network coverage is expanded through the addition of genes, reactions, and metabolites supported by literature evidence. The quality of the reconstruction was improved through an update of standardized formatting, improved annotation, and the addition of binning metabolites representing macromolecular categories to assist with analysis. The metabolic network model was validated by comparing phenotypic predictions to experimental datasets^16–21^ and the quality of the reconstruction was assessed with the MEMOTE benchmarking software^22^. This updated reconstruction was further contextualized with recently published transcriptomic data^1^ in order to demonstrate its utility in elucidating the metabolic shifts of *P. aeruginosa* after exposure to mucins. The validated reconstructions will serve as a key resource for the *Pseudomonas* and microbial metabolic modeling communities and the insights into mucin-driven metabolic shifts in *P. aeruginosa* may serve to inform the future development of therapeutic strategies.

## Results

### *An updated network reconstruction of* Pseudomonas aeruginosa *metabolism*

A metabolic network reconstruction of *P. aeruginosa* PA14 (iPau1129) was previously published^16^ and served as a starting point for an updated reconstruction (iPau21). The metabolic coverage of the reconstruction was expanded, the format and annotations were updated, and an ATP-generating loop was resolved in order to produce a refined model with improved accuracy and extensive annotation.

We expanded iPau1129 by 40 genes, 24 metabolites, and 76 reactions (Fig. 1a) through manual curation based on literature evidence (Supplementary Data 1). Many of these additions served to increase the utility of the reconstruction for simulation (such as the addition of 33 exchange reactions), while others expanded metabolic pathways for amino acid metabolism and glycerophospholipid metabolism. A periplasmic compartment containing hydrogen was added to the reconstruction to better represent the electron transport chain and ATP synthase, which eliminated all ATP-generating loops in the metabolic network. The format was updated from SBML Level 2^23^ to Level 3^24^, which enables additional functionality such as the utilization of several extension packages and the transfer of information content to dedicated new data structures. Annotations from various databases were added to metabolites, reactions, and genes where possible.

**Figure 1:**
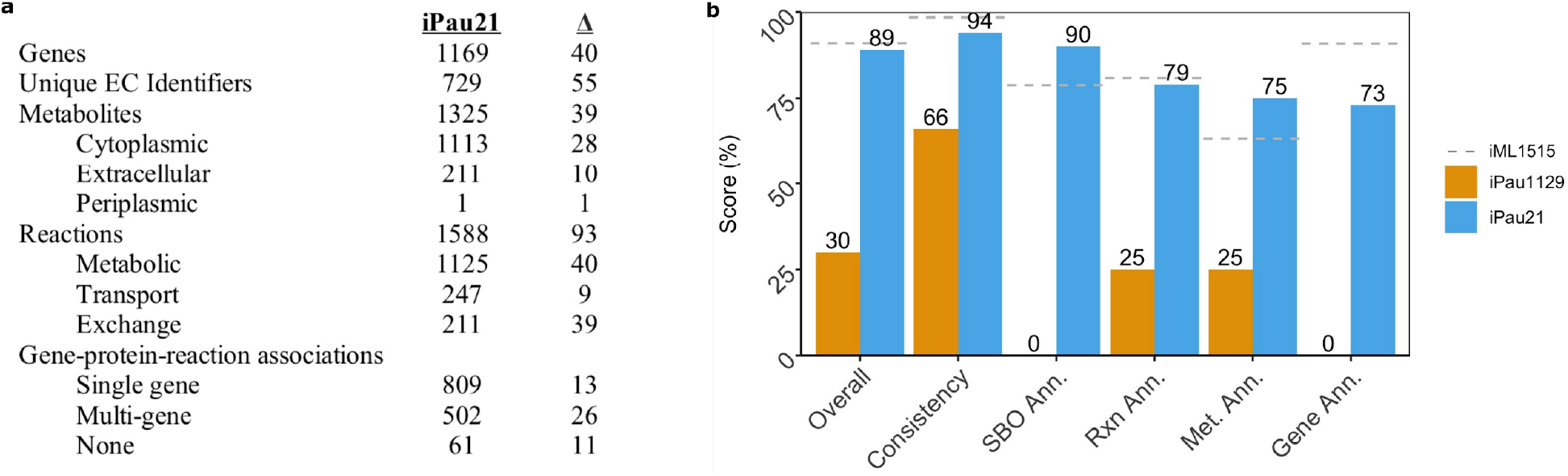
Characteristics and MEMOTE benchmarking of iPau21. **(a)** Properties of iPau21 as compared to iPau1129. **(b)** MEMOTE scores of iPau21, iPau1129, and iML1515, a high-quality reconstruction of *E. coli*.

The overall quality of the updated reconstruction was assessed using MEMOTE^22^, a recently developed GENRE test suite. The MEMOTE score of iPau21 improved in all subcategories when compared to iPau1129 resulting in an increase of the overall score from 30% to 89% (Fig. 1b and Supplementary Data 2). The scores in annotation subcategories were increased by adding annotations and SBO terms to metabolites, reactions, and genes in the updated GENRE. The consistency of the metabolic network was improved through the correction of imbalanced reactions and the resolution of energy generating cycles that were present in iPau1129.

The biomass objective function (BOF) was updated to better reflect the macromolecular components found experimentally in *P. aeruginosa* including the inclusion of lipopolysaccharide^25–27^. BOF substrates were organized into corresponding macromolecular categories (i.e. DNA, RNA, protein, lipid) to better represent the categories of components that are required for growth.

### Model validation

Validation of iPau21 was performed by comparing *in silico* predictions of biomass flux, carbon source utilization, and gene essentiality to experimental data. Biomass flux and subsequent doubling time predictions in simulated lysogeny broth (LB), synthetic cystic fibrosis media (SCFM), and glucose minimal media were compared to experimental values found in literature (Fig. 2a, Supplementary Data 3)^17–19^. Doubling time predictions of iPau21 were 25%, 19%, and 22% more accurate than those of iPau1129 in simulated LB, SCFM, and glucose minimal media, respectively. Compared to the original model, iPau21 doubling times are higher, which reflects the resolution of the ATP-generating loop that previously allowed the model to costlessly convert ADP to ATP. The iPau21 doubling time prediction on glucose minimal media of 40.2 minutes showed agreement with experimental data, falling within the range of experimentally determined values^19^. Model doubling time predictions on LB and SCFM were faster than observed experimentally, which is consistent with metabolic network models that are structured to predict the optimal growth of an organism.

**Figure 2:**
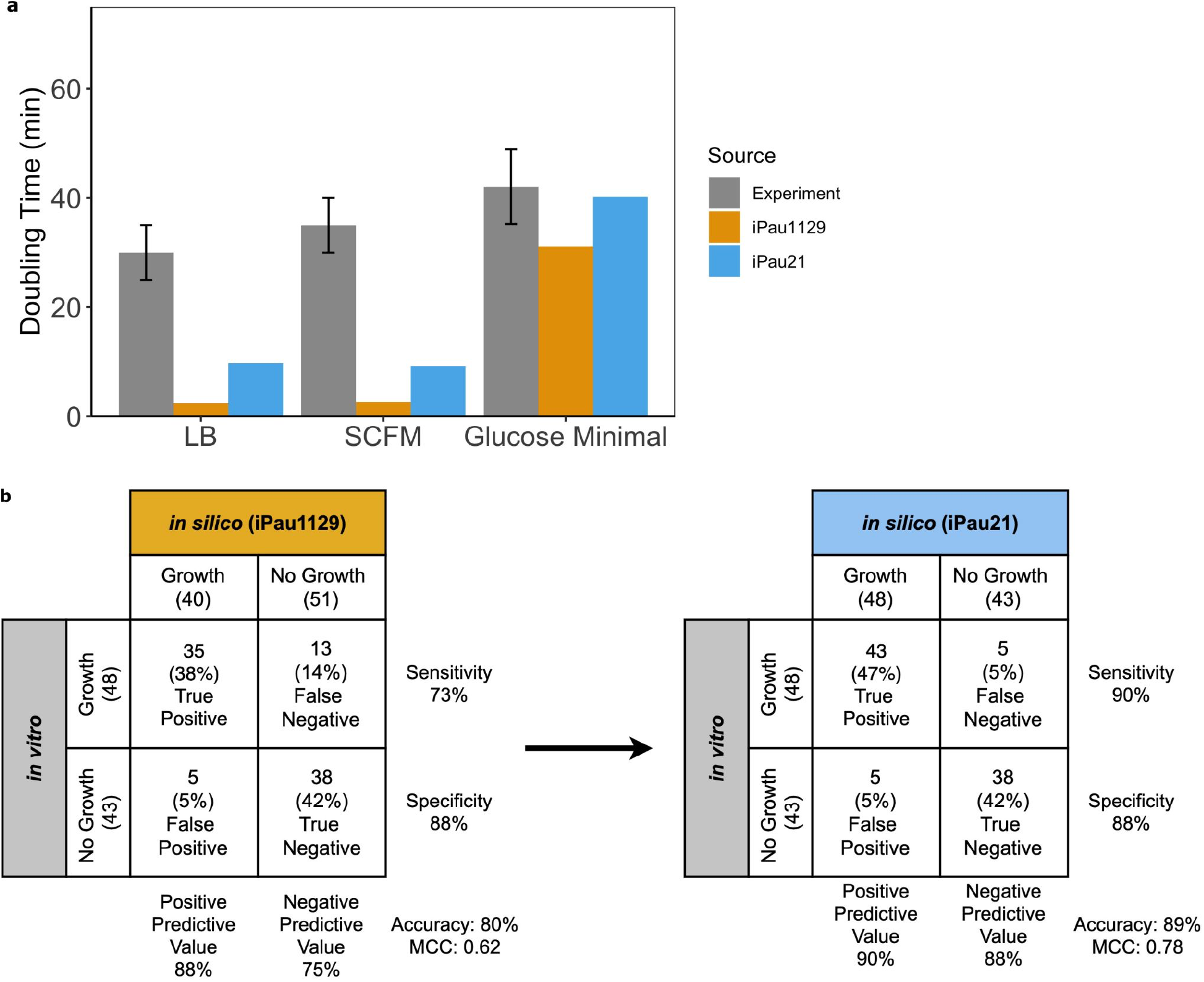
Updated reconstruction of *P. aeruginosa* enables accurate growth rate, gene essentiality, and carbon source utilization predictions. **(a)** Model doubling time predictions compared to results gathered from literature. **(b)** Model carbon source utilization predictions compared to results gathered from literature.

Carbon source utilization predictions were compared to previously collected experimental results across 91 carbon sources^16^. Utilization was predicted by iPau21 with an accuracy of 89% and Matthews correlation coefficient (MCC) of 0.78, while iPau1129 demonstrated an accuracy of 80% and MCC of 0.62 (Fig. 2b and Supplementary Data 4). This increase in accuracy was achieved through the completion of pathways that allow for the utilization of more carbon sources and the removal of an unsupported reaction that previously allowed for the utilization of D-malate. Carbon source predictions of iPau21 remain incorrect for 10 carbon sources. Five of the incorrect predictions are due to the absence of metabolic pathways required for growth on certain carbon sources. When addressing these predictions, our literature survey was unable to provide sufficient evidence for these pathways so the predictions remain incorrect and we opted to not gapfill without that additional evidence. To correct these predictions, summary reactions could be added to the reconstruction, but these reactions would lack the mechanistic granularity of associated genes and could have negative impacts on other aspects of the reconstruction. The other five incorrect predictions were caused by the presence of metabolic pathways that allow for the erroneous growth on the associated carbon sources. In each of these cases, the pathway was investigated and the corresponding genes were verified through the KEGG^28^ and ModelSEED^29^ databases, but there was not strong enough evidence to warrant changes in the reconstruction^28,30^. Some of these discrepancies may be due to considerations that are outside of the scope of the network, such as transcriptional processes. For example, in the case of D-serine, PA14 has the ability to metabolize this carbon source but expression of this gene is not triggered by the presence of D-serine so it is unable to grow on this single carbon source *in vitro*^31^. These inaccurate predictions could be improved by modifying constraints in the metabolic network model. However, since the gene-protein-reaction (GPR) rules were found to be valid and the prediction error could be due to unaccounted for regulatory control, we opted to leave the pathways intact. Overall, we were able to increase carbon source utilization prediction accuracy by 9% in comparison to the previously published model.

Gene essentiality predictions were compared to a published dataset comprised of the overlap of essential genes identified through the growth of strains PAO1 and PA14 transposon insertion mutants in LB media^20,21^. The number of genes accounted for by iPau21 was expanded to 1169 and the gene essentiality prediction accuracy was maintained at 91%, which is equivalent to iPau1129 (Supplementary Data 5). Gene essentiality was predicted by iPau21 with a MCC of 0.50, compared to a value of 0.44 by iPau1129. Three genes labeled as “SPONTANEOUS,” “unassigned,” and “Unassigned” were removed from the reconstruction given that these labels did not correspond to genes belonging to *P. aeruginosa*. Gene essentiality data was not used for curation of the metabolic network given the variability in gene essentiality screens and the resultant challenges with data interpretation^32^. Instead, model predictions were compared to gene essentiality data as one facet of validation. As a reference, iPau21 has a gene essentiality prediction accuracy of 91%, which is near the 93% accuracy of iML1515, a well-curated reconstruction of *Escherichia coli*^33^.

### *Transcriptome-guided modeling of* P. aeruginosa *metabolism in the presence of human mucins*

Mucins are the primary macromolecules in mucosal layers known to modulate microbial phenotypes^2^. In order to investigate how the metabolism of *P. aeruginosa* shifts when it comes into contact with mucins, *in vitro* transcriptomic data was integrated with iPau21 to generate contextualized models that offer more biologically accurate representations of associated metabolic phenotypes. Analysis of the structure and pathway utilization in these transcriptome-guided models offers insights into the metabolic shifts that arise when *P. aeruginosa* is exposed to mucins.

Transcriptomic profiles of *P. aeruginosa* PAO1 grown in agrobacterium minimal medium with thiamine, glucose, and casamino acids (ABTGC) medium supplemented with either MUC5AC, MUC5B, or mucin-glycans were collected from literature^1,34^. MUC5AC and MUC5B are mucin types found both individually and together at different sites of the human body that *P. aeruginosa* is known to infect^8^. The mucin-glycans used in the published experiments were isolated from the backbone of MUC5AC. The experiments were performed with strain PAO1, which has a highly similar genome to strain PA14^35^. The main difference between the strains is the presence of additional gene clusters in PA14 (most linked to virulence) that we would not expect to have a large effect on overall metabolism. PAO1 genes in the transcriptomic dataset were mapped to PA14 orthologs and then the data was integrated with the iPau21 using the RIPTiDe algorithm^36^. RIPTiDe uses transcriptomic evidence to create context-specific metabolic models representative of a parsimonious metabolism consistent with the transcriptional investments of an organism. This analysis resulted in four contextualized models that more accurately represent the metabolism of *P. aeruginosa* when grown without mucin exposure (ABTGC) and when exposed to MUC5AC, MUC5B, and glycans.

Flux samples were generated for each model and BOF flux did not vary significantly among the contextualized models (less than five percent change), recapitulating the phenotype that was observed experimentally^1^. The flux distributions underlying the BOF values were compared across models using non-metric multidimensional scaling (NMDS) in order to compare the metabolic mechanisms of growth utilized by the condition-specific metabolisms (Fig 3a). The fluxes from the 378 consensus reactions (shared across all models) were used for this analysis. NMDS analysis revealed that among the tested conditions, the sampled flux distributions from the MUC5B model clustered the furthest from the ABTGC condition. This result indicates that although there was not a significant difference in the BOF value, exposure to MUC5B caused the largest shift in the metabolic pathways utilized for growth. MUC5AC clustered the second furthest away, while Glycans clustered most closely to the ABTGC model, showing that there was a variable metabolic response to different mucins and glycans by *P. aeruginosa*. Mucin-glycans do not contain the same level of structural and biochemical complexity as MUC5AC and MUC5B, which may account for the slight metabolic shift observed in the Glycans model relative to the MUC5AC and MUC5B models. MUC5AC and MUC5B are known to differ from each other in terms of charge, shape, and glycosylation^37^. These differences could explain the variable metabolic response they elicit in *P. aeruginosa*. Additionally, of the two only MUC5B has been shown to be critical for murine mucociliary transport and antibacterial defense^38^. One mechanism of MUC5B antibacterial effects could be through modulation of pathogen metabolism, which would explain the larger shift in metabolism observed when *P. aeruginosa* was exposed to MUC5B. The conserved BOF flux values and separation observed between clusters of flux samples suggest that while *P. aeruginosa* metabolism is modulated by the presence of mucins, its versatility allows for the utilization of alternative metabolic pathways in order to avoid a growth defect.

**Figure 3:**
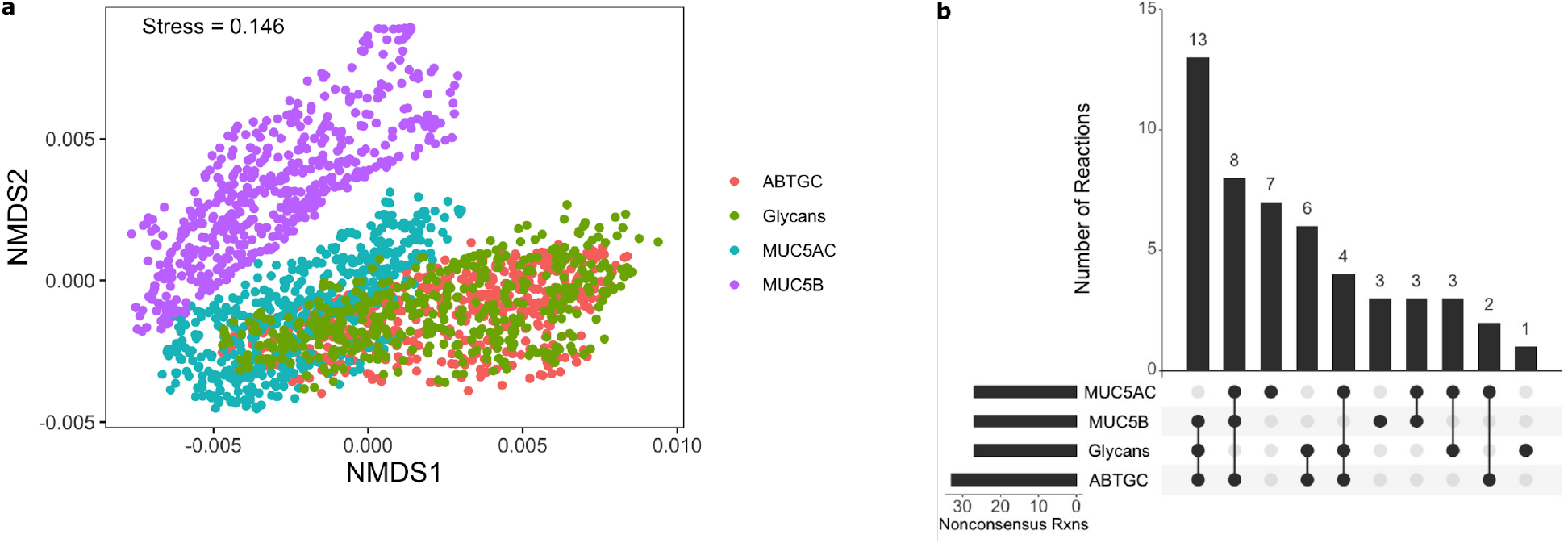
Contextualization of updated reconstruction shows shifts in *P. aeruginosa* metabolism in response to mucins and mucin components. **(a)** NMDS analysis of flux samples (*n* = 500) from each contextualized model. **(b)** Comparison of non-consensus reactions present within models displays subsets of reactions that are shared by groups of contextualized models.

The differences between networks were further investigated through the metabolites that were produced and consumed by models *in silico*. This analysis offers a snapshot of the substrates used and byproducts of particular metabolic states, which can be informative of the metabolism underlying that state. All models were found to consume the same metabolites with some small differences in specific flux values however, there were key differences in the metabolites that models produced (Supplementary Data 6). The graded differences between models seen in NMDS were highlighted by the production of formate by the models. The ABTGC and Glycans models produced substantially higher amounts of formate than the MUC5AC model, while the MUC5B model did not contain the formate exchange reaction. Therefore, with our model, we are able to predict subtle shifts in *P. aeruginosa* metabolism in response to different environmental mucins.

### *Human mucins shift* P. aeruginosa *metabolism*

Further analysis was conducted on the contextualized models to better understand the shifts in metabolism that resulted in the observed dissimilarities in the NMDS analysis. Reactions not shared across all models (non-consensus reactions) were identified and compared to investigate how network structure varies across models (Fig. 3b). This analysis revealed a set of 13 reactions shared by the ABTGC, MUC5B, and Glycans models but absent from the MUC5AC network. This result suggests that while MU5B displayed the largest functional differences in metabolism, MUC5AC is the most structurally unique of our models. Additionally, we found that there was no correlation between network structure and utilization among our contextualized models (Fig. 4, *p*-value = 0.92). Since the NMDS analysis revealed that the ABTGC and MUC5B models had the largest difference in functional metabolism, these two models were further investigated to find key attributes that underlie these large differences. Random forest analysis was conducted on the flux samples from consensus reactions of the ABTGC and MUC5B models to find which reactions were most differentially utilized between the two cases (Fig. 5). Two reactions corresponding to fumarate transport were in the top seven most discriminating reactions between models, suggesting that there was a differential utilization of reactions involved in fumarate metabolism. The MUC5B model utilized the fumarate reactions more highly than the ABTGC model and contained a fumarase reaction that was not present in the ABTGC model, which further suggests that fumarate metabolism is a key point of difference between the models. This observation recapitulates what was noted in the original paper that produced the transcript data used for contextualization^1^. Of the top six most discriminating reactions, five corresponded to propionate metabolism and were more highly utilized by the MUC5B model than the ABTGC model. While there was no propionate in the simulated (or *in vitro*) media, it is a known byproduct of mucin fermenters and has anti-lipogenic and anti-inflammatory properties in humans^39,40^. This analysis revealed that the exposure to MUC5B elicited the largest shift in metabolism compared to MUC5AC and Glycans. Further, an increased utilization of fumarate and propionate metabolism during simulated growth was responsible for this shift.

**Figure 4:**
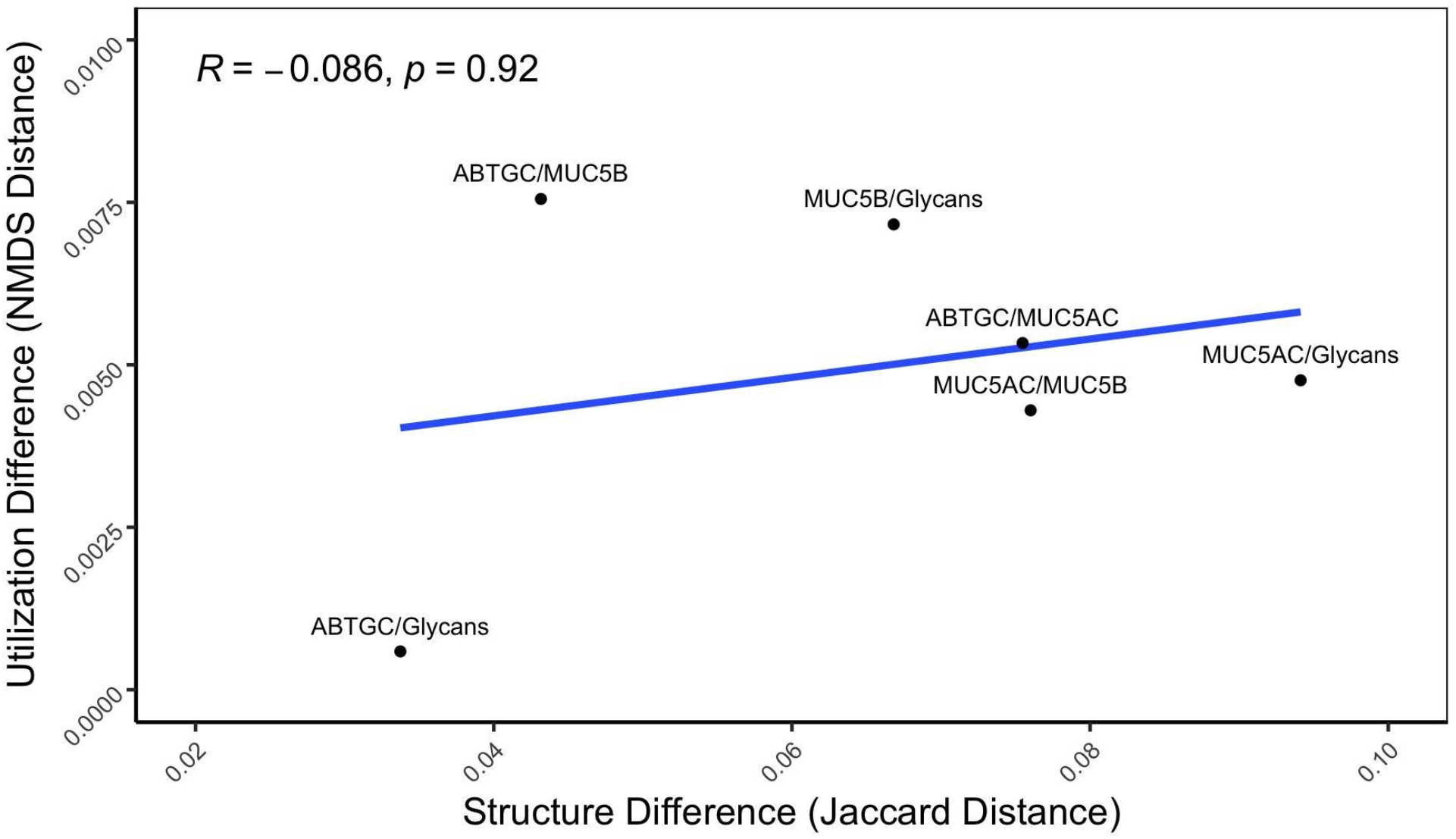
Network utilization does not correlate with network structure. The distance between median NMDS coordinates for each pair of networks was calculated as a metric of difference in network utilization, while the Jaccard distance of network reactions for each pair of networks was calculated as a metric of structural difference. Spearman’s correlation shows an insignificant relation between the two metrics (*p* = 0.92).

**Figure 5:**
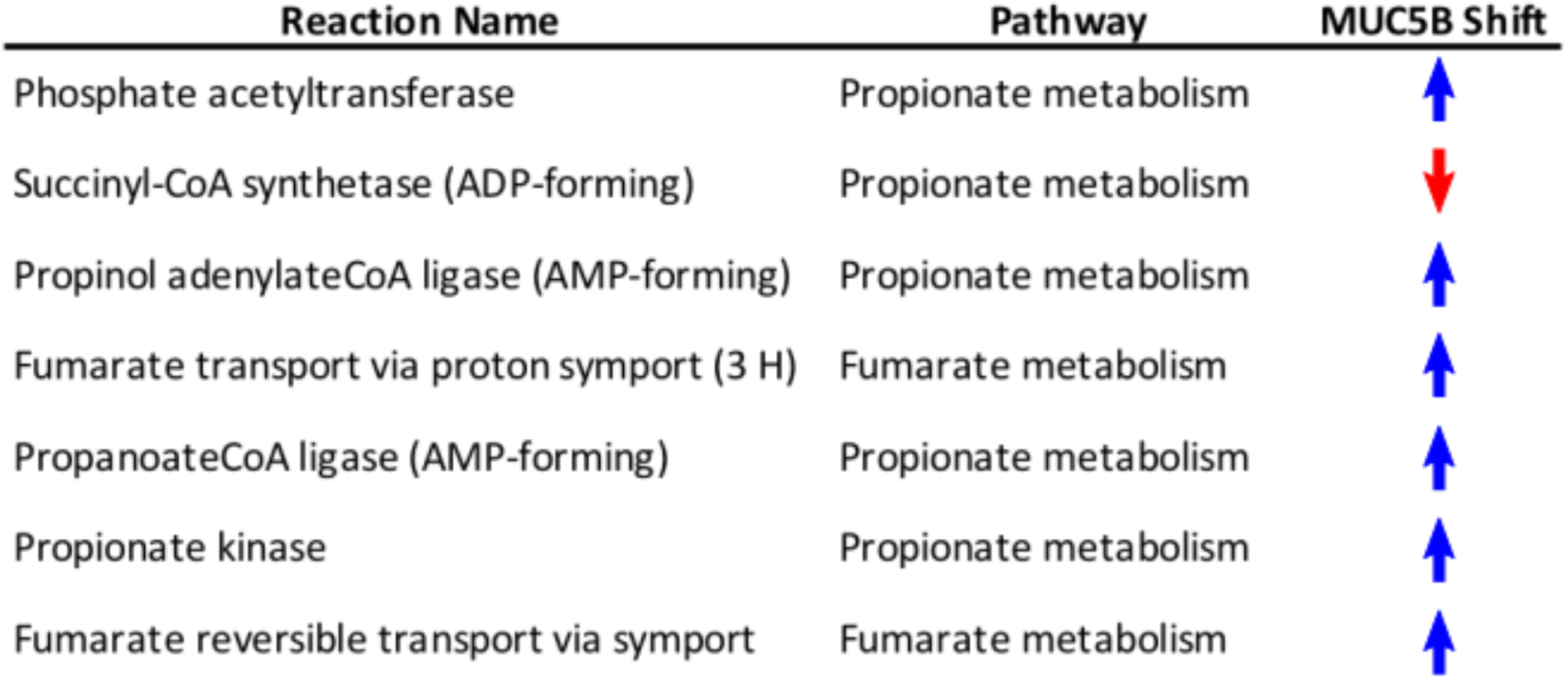
Random forest analysis between ABTGC and MUC5B shows the networks differ most in terms of propionate and fumarate metabolism utilization. The top seven most discriminating reactions between the two models belong to propionate and fumarate metabolism. MUC5B utilizes these two types of metabolism more highly than the ABTGC model.

## Discussion

We generated an updated network reconstruction of *P. aeruginosa* PA14 metabolism with considerable improvements in model annotation and accuracy of growth rate and carbon source utilization predictions. The metabolic reaction coverage of the reconstruction was expanded, the format and annotations were updated to be consistent with current best practices, and an ATP-generating loop was resolved. Model improvements were quantified through various metrics such as accuracy of growth yield and carbon source utilization predictions as well as MEMOTE benchmarking^22^.

The updated network reconstruction was contextualized using transcriptomic data in order to investigate the shifts in metabolism that occur when *P. aeruginosa* is exposed to mucins present in the human body. This analysis recapitulated an unaltered growth rate and differential fumarate metabolism that has been reported in literature and also revealed an increased utilization of propionate metabolism in the presence of mucins. Propionate is a short chain fatty acid with beneficial effects to human health such as anti-lipogenic, anti-inflammatory, and anti-carcinogenic action^39,41^. While propionate is not present in the ABTGC medium, it is known to be produced by bacteria such as *Akkermansia muciniphila* when they come into contact with and catabolize mucins^40^. This shift of *P. aeruginosa* metabolism towards propionate metabolism may indicate a cross-feeding mechanism where MUC5B mucins signal to *Pseudomonas* to prepare to metabolize the propionate produced by other microbes as they break down the mucins. Once validated, this insight could be used to develop therapeutic strategies for *P. aeruginosa* infections of body sites containing MUC5B such as the lung, oral cavity, and middle ear^1^. Antibiotics could be designed to target proteins for propionate metabolism in order to combat drug-resistant strains that cannot be treated with traditional antibiotics.

While the updates made to the model broadly improved the model accuracy, there were incorrect predictions about carbon source utilization, gene essentiality, and growth rate that were not able to be addressed. Some incorrect predictions are due to a lack of literature evidence, such as incorrect carbon source utilization predictions that are due to the absence of metabolic pathways in the model. Other incorrect predictions are due to factors that are outside of the scope of the model, such as the incorrect prediction of growth on D-serine that is caused by the transcriptional regulation of *dsdA*^31^. There are other opportunities for further curation that would result in additional improvements to the MEMOTE score, which can be further interrogated by uploading the iPau21 reconstruction to the MEMOTE website (MEMOTE.io).

The transcriptomic data that was used to contextualize the model was collected through experiments with *P. aeruginosa* strain PAO1. Therefore, the genes in the transcriptomic data set were mapped to their PA14 orthologs before being integrated with the network reconstruction. While the genomes of PAO1 and PA14 are highly similar, the PA14 (6.5Mb) genome is slightly larger than PAO1 (6.3Mb) and contains gene clusters that are not present in PAO1^35^. The genes absent in PAO1 therefore would not be accounted for in the transcriptomic dataset. However, since most of these genes are linked to virulence, they should not have large effects on whole metabolism as simulated here. Therefore, we expect that this application of the model would allow the identification of broad shifts in metabolism due to exposure to mucins irrespective of the specific strain simulated.

The improvements in the *P. aeruginosa* metabolic network reconstruction were made to reconcile key disagreements between *in silico* predictions and *in vitro* results, ultimately producing a higher quality metabolic network reconstruction. Through the update process, we identified key predictions that remain incorrect and offer targets for further curation. The application of the model to investigate metabolic shifts that occur upon exposure to mucins recapitulated phenotypes observed in literature and offered mechanistic insights that would be difficult to delineate experimentally. This application of the reconstruction serves as an example of how the reconstruction and associated models can provide insights into context-specific metabolism. Ultimately, this reconstruction can serve as a resource for investigating the metabolism of *P. aeruginosa* in a variety of settings and conditions.

## Methods

### Genome-scale Metabolic Reconstructions and Models (GENREs and GEMs)

GENREs are network reconstructions that represent the metabolic capabilities of an organism and can be analyzed for various applications. An organism’s genes are connected to the proteins they code for and the reactions that those proteins catalyze. These associations are stored as gene-protein-reaction (GPR) relationships with the reactants and products of each reaction catalogued in a stoichiometric matrix. Metabolites in the reconstruction are assigned to compartments that mirror biologically discrete spaces such as the cytosol and the extracellular space. Exchange and transport reactions allow metabolites to flow between the compartments in the reconstruction. A GENRE is turned into a GEM (Genome-Scale Metabolic Model) by adding reaction bounds that capture the flux constraints and the reversibility of reactions. The flux bounds dictate the amount and direction of flux that a reaction can carry. Objective functions (OFs) that represent metabolic goals are added to the model to simulate biological processes. GEMs can be analyzed using flux balance analysis (FBA)-based methods to investigate and gain insights into the metabolic state of a network^42^. The updated GENRE was named iPau21 according to the community standard naming convention^43^.

### Adding annotations

#### Metabolites and Reactions

Initially, the PA14 reconstruction did not contain extensive annotations for metabolites, reactions, or genes. ModelPolisher^44^ can be used to annotate metabolites and reactions of a metabolic model. To do so, identifiers of the BiGG database^45^ (BiGG-IDs) are required as metabolite or reaction identifiers, respectively. Since the identifiers of the model were obtained from the ModelSEED database^30^, BiGG-IDs needed to be determined. For each metabolite, the BiGG-IDs were assessed manually. Since this is a very time-consuming procedure, the BiGG-IDs for the reactions were resolved in a semi-automated way: The cross-references of the ModelSEED database to other databases, such as BiGG or KEGG^28^, were used to automatically obtain the BiGG-IDs for the respective ModelSEED reaction identifier. If more than one BiGG-ID was returned, the correct identifier was determined by manual inspection of the respective reaction. The BiGG-IDs of the metabolites and reactions were added as biological qualifier (‘BQB_IS’) annotations to the model using libSBML Version 5.17.0^46^. The annotations were added in accordance with the MIRIAM guidelines^47^. After adding the BiGG-IDs to the model, ModelPolisher was used for further annotations of the model’s reactions and metabolites for references to other databases, such as KEGG, MetaNetX^48^, or MetaCyc^44^.

For the reactions, the obtained KEGG annotations were used to further add all pathways that are associated with the respective reaction to the model. The pathways were obtained using the KEGG-ID and KEGG API to request all associated pathways. The pathways were then added to the respective reactions using the biological qualifier ‘BQB_OCCURS_IN’ in libSBML.

#### Genes

The identifiers of the model genes are from the KEGG database. With the help of libSBML, the KEGG gene annotation was added to the model. For further gene annotations, the KEGG API was used to request NCBI^49^ Protein IDs and Uniprot^50^ IDs, which were subsequently added as respective annotations to the model. Additionally, the ID mapper from PATRIC^51^ was used to request RefSeq and NCBI^49^ gene identifiers, as well as identifiers of the ASAP database.

#### SBO-Terms

Systems Biology Ontology (SBO)^52^ terms can give semantic information or be used for annotation purposes. In our network reconstruction, all genes were labelled as genes with the SBO-term ‘SBO:0000243’. All metabolites without a valid SBO-term were labelled as simple chemicals with the SBO-term ‘SBO:0000247’. Transport reactions were divided into (1) active transport if ATP is required for the respective transport reaction (SBO:0000657), (2) passive transport if no external energy is required (SBO:0000658), (3) symporter-mediated transport if two or more molecules are transported into the same relative direction across a membrane (SBO:0000659), or (4) antiporter-mediated transport if two or more molecules are transported in relative opposite directions across a membrane (SBO:0000660).

All metabolic reactions were labelled as biochemical reactions with the SBO-term SBO:0000176.

### Upgrading SBML version

The initial PA14 reconstruction was represented in SBML Level 2 Version 1^53^. The current reconstruction was updated to the latest SBML edition (Level 3)^54^. With the help of libSBML, both the fbc-plugin^55^ and the groups-plugin^56^ were enabled.

#### Fbc-plugin

Initially, the chemical formulas and charges of the metabolites were stored in the notes field. With the fbc-plugin, the charges were added as features of the metabolites to the reconstruction. The fbc-plugin also enables the addition of gene products to the reconstruction.

#### Groups-plugin

In the initial reconstruction, the subsystems of the reactions were saved in the notes field. With libSBML and the groups-plugin, the subsystems were extracted from the notes field and added as groups to the reconstruction. For each subsystem, a list of reactions associated with that pathway according to the notes was created and added to the subsystem as members.

### Correcting charge and mass imbalances

A list of all mass- and charge-imbalanced reactions was extracted from the reconstruction. From this list, all exchange, sink, demand and biomass reactions were excluded. Each remaining reaction was manually checked by looking up the reaction-ID in ModelSEED^29^: (1) If the reaction status in ModelSEED was balanced (‘OK’), but differed from the reaction equation in the reconstruction, the reaction was adapted according to ModelSEED and again checked for imbalances. (2) If the reaction in ModelSEED also had an imbalanced reaction status, other databases like MetaCyc^57^, BiGG^45^, or KEGG^28^ were explored and the reactions were adapted according to the respective reactions in the external databases. Where required, chemical formulas, charges, and coefficients were corrected, or chemical compounds were added or subtracted from the reactions according to the respective database reaction. All changed reactions are listed in Supplementary Data 1.

### Assessing the quality of the reconstruction

MEMOTE is an open-source software that provides a measure for model quality^22^. Every change and improvement of the model was continuously documented and quality-assessed using MEMOTE Version 0.9.11. Full MEMOTE reports are provided for iPau1129, iPau21, and iML1515 (Supplementary Data 2). Gene essentiality predictions were compared to a published dataset that was originally used to validate iPau1129^16^. This dataset is comprised of the overlap of essential genes identified through the growth of PAO1 and PA14 transposon insertion mutants in LB media^20,21^. Carbon source utilization predictions were compared to previously collected experimental results^16^. Prediction accuracy was calculated as the number of correction predictions divided by the number of total predictions. Matthews correlation coefficient (MCC) was calculated in order to assess the quality of predictions^58^. Biomass flux and subsequent doubling time predictions in lysogeny broth (LB), synthetic cystic fibrosis media (SCFM), and glucose minimal media were compared to experimental values found in literature (Fig. 1c)^17–19^.

### Literature-based updates

Previous work identified multiple areas where the original reconstruction (iPau1129) was unable to accurately recapitulate experimental data. This assessment included 18 incorrect carbon source predictions^16^ and several incorrect gene essentiality predictions^59^. Pathways and gene-protein-reaction rules related to each incorrect prediction were manually curated to reflect the most recent evidence from literature, KEGG, and MetaCyc. In the absence of sufficient evidence, no changes were made, even if this absence of a change meant a prediction would remain uncorrected.

### Evaluating and updating the BOF

Macromolecular categories represented in the dry weight of *P. aeruginosa* were identified through a literature survey. Metabolites in the biomass objective function (BOF) were organized into these macromolecular categories in order to better represent the components required for growth. During organization, no additional metabolites were added and the ratios of metabolites in the BOF were kept the same.

The BOF was also updated to include lipopolysaccharide (cpd17065) to reflect its presence in Gram-negative bacteria^60^. A metabolite representing biomass was also added to the products of the BOF to represent the accumulation of biomass.

### Addition of exchange reactions

A list of all extracellular metabolites in the reconstruction was compiled and compared to a list of all exchange reactions in the reconstruction. Exchange reactions were added for 33 extracellular metabolites that previously did not have one.

### Removal of energy generating cycles

Exchange reactions were closed and the objective function was set to energy dissipation reactions for electron carriers (ATP, NADH, NADPH, FADH_2_, and H^+^). The model was able to generate flux for only the ATP energy dissipation objective function, which indicated that an energy generating cycle existed. The cycle was resolved through the addition of a periplasm compartment to contain hydrogen involved in the electron transport chain and correcting the reversibility of four participating reactions.

### RIPTiDe Contextualization & Analysis

Published transcriptomic data was integrated with the model using RIPTiDe^36^. The transcriptomic data was normalized then translated from PAO1 genes to the orthologous PA14 genes prior to integration^61^. ABTGC medium was simulated *in silico* and applied to the model (Supplementary Data 3). Then, RIPTiDe was used to produce the contextualized models for *in vitro* media conditions.

NMDS analysis was conducted on flux samples from each contextualized model (*n* = 500 samples per model) using the Vegan package in R^62^. Only consensus reactions across all four contextualized models were included in the flux sample data set and a constant was added to each flux value in the data set to make all data points positive to facilitate comparison. Median fluxes for every reaction in each model are provided in Supplementary Data 7.

Random forest analysis was conducted on flux sampling data (*n* = 500 samples per model) from the consensus reactions of the ABTGC and MUC5B models using the randomForest package in R^63^. Reactions that were differentially present in contextualized models were identified and connected to their corresponding metabolic pathways manually.

The Jaccard distance of network structures was calculated by comparing the reactions contained in pairs of networks^64^. The NMDS distance was calculated as the distance between the median NMDS coordinates of network pairs. Spearman’s correlation was used to calculate a *p*-value for the relationship between network structure and network utilization across all pairs of networks.

## Data and code availability

All code and data for this project are available on GitHub (github.com/dawsonpayne/iPau21). The genome-scale metabolic model iPau21 is available in the BioModels Database^65^ as an SBML Level 3 Version 154 file within a COMBINE Archive OMEX file^66^ including the contextualized models and metadata^67^ at identifiers.org/biomodels.db/MODEL2106110001.

## Acknowledgements

The authors would like to thank our colleagues Jennifer Bartell and Anna Blazier for their previous work on iPau1129 and their helpful suggestions for our curation of iPau21.

This work is supported with U.S. federal funds from the National Institutes of Health (R01 AI154242) and funded by the *Deutsche Forschungsgemeinschaft* (DFG, German Research Foundation) under Germany’s Excellence Strategy – EXC 2124 – 390838134 within the Cluster of Excellence CMFI (Controlling Microbes to Fight Infections). AD is funded by the Germany Center for Infection Research (DZIF) within the *Deutsche Zentren der Gesundheitsforschung* (BMBF-DZG, Germany Centers for Health Research of the Federal Ministry of Education and Research), grant № 8020708703.

## Competing interests

The authors declare no competing interests.

## Author contributions

L.D. and J.P. conceived the project. D.P., A.R., L.D., and T.L. researched for GENRE updates. D.P., A.R., and L.D. prepared the GENRE. D.P. ran simulations with the GENRE and produced/analyzed contextualized models. D.P., L.D. and J.P. interpreted the results. D.P., A.R., and L.D. wrote the manuscript. All authors critically revised the manuscript and approved the final version.

## List of abbreviations

(ABTGC): Agrobacterium minimal medium with thiamine, glucose, and casamino acids
(ADP): Adenosine diphosphate
(ATP): Adenosine triphosphate
(BiGG): Biochemical Genetic and Genomic
(BOF): biomass objective function
(DNA): deoxyribonucleic acid
(GPR): gene-protein-reaction
(FBA): flux balance analysis
(FADH2): flavin adenine dinucleotide
(GEMs): genome-scale metabolic models
(GENREs): genome-scale metabolic network reconstructions
(KEGG): Kyoto Encyclopedia of Genes and Genomes
(LB),: lysogeny broth
(MCC): Matthews correlation coefficient
(MEMOTE): metabolic model testing
(NADH): reduced nicotinamide adenine dinucleotide
(NADPH): Nicotinamide adenine dinucleotide phosphate
(NMDS): non-metric multidimensional scaling
(PA14): UCBPP-PA14
(RIPTiDe): Reaction Inclusion by Parsimony and Transcript Distribution
(SBO): Systems Biology Ontology
(SBML): Systems Biology Markup Language
(SCFM): synthetic cystic fibrosis media
(RNA): ribonucleic acid
(WHO): World Health Organization

## Notes

https://github.com/dawsonpayne/iPau21

https://identifiers.org/biomodels.db/MODEL2106110001

### Competing Interest Statement

The authors have declared no competing interest.

