## Supplemental Data 1 for "An updated genome-scale metabolic network reconstruction of *Pseudomonas aeruginosa* PA14 to characterize mucin-driven shifts in bacterial metabolism"

Note: "Code Line" corresponds to "iPau21_notebook.ipynb"

| Code Line | Model Object | Change | Justification | Lit Reference |
|---|---|---|---|---|
| 11 | rJB00275 | Remove | Leftover demand reaction from previous analysis | - |
| 12 | rJB00280 | Remove | Leftover demand reaction from previous analysis | - |
| 13 | rJB00281 | Remove | Leftover demand reaction from previous analysis | - |
| 14 | EX_cpd00147_c | Remove | Leftover demand reaction from previous analysis | - |
| 16-24 | cJB00034_e | Change to ModelSeed ID (cpd00212_e) and add annotations | Metabolite match in ModelSeed | - |
| 26-29 | cPY00161_c | Change to ModelSeed ID (cpd23005_c) and add annotations | Metabolite match in ModelSeed | - |
| 32-39 | rJB00259 | Change to ModelSeed ID (rxn36698) and add annotations | Reactions match in ModelSeed | - |
| 41-47 | rJB00260 | Change to ModelSeed ID (rxn37440) and add annotations | Reactions match in ModelSeed | - |
| 49-54 | rJB00262 | Change to ModelSeed ID (rxn36761) and add annotations | Reactions match in ModelSeed | - |
| 56-61 | rJB00265 | Change to ModelSeed ID (rxn36840), add annotations, and update GPR based on ModelSeed/Kegg Databases | Reactions match in ModelSeed | - |
| 63-68 | rJB00266 | Change to ModelSeed ID (rxn37076) and add annotations | Reactions match in ModelSeed | - |
| 70-77 | rJB00276 | Change to ModelSeed ID (rxn10902) and add annotations | Reactions match in ModelSeed | - |
| 79-84 | rPY00216 | Change to ModelSeed ID (rxn42682) and add annotations | Reactions match in ModelSeed | - |
| 86-90 | rPY00220 | Change to ModelSeed ID (rxn21407) and add annotations | Reactions match in ModelSeed | - |
| 93-101 | EX_cJB00127_e (cis-2-Decenoic acid -->) | Add exchange reaction for extracellular metabolite | Added for ease of media simulation | - |
| 104-112 | EX_cpd00047_e (Formate -->) | Add exchange reaction for extracellular metabolite | Added for ease of media simulation | - |
| 114-122 | EX_cpd00059_e (L-Ascorbate -->) | Add exchange reaction for extracellular metabolite | Added for ease of media simulation | - |
| 124-132 | EX_cpd00085_e (beta-Alanine -->) | Add exchange reaction for extracellular metabolite | Added for ease of media simulation | - |
| 134-142 | EX_cpd00136_e (4-Hydroxybenzoate -->) | Add exchange reaction for extracellular metabolite | Added for ease of media simulation | - |
| 144-152 | EX_cpd00147_e (5-Methylthioadenosine -->) | Add exchange reaction for extracellular metabolite | Added for ease of media simulation | - |
| 154-162 | EX_cpd00152_e (Agmatine -->) | Add exchange reaction for extracellular metabolite | Added for ease of media simulation | - |
| 164-172 | EX_cpd00170_e (Gluconolactone -->) | Add exchange reaction for extracellular metabolite | Added for ease of media simulation | - |
| 174-182 | EX_cpd00186_e (D-Glutamate -->) | Add exchange reaction for extracellular metabolite | Added for ease of media simulation | - |
| 184-192 | EX_cpd00214_e (Palmitate -->) | Add exchange reaction for extracellular metabolite | Added for ease of media simulation | - |
| 194-202 | EX_cpd00222_e (GLCN -->) | Add exchange reaction for extracellular metabolite | Added for ease of media simulation | - |
| 204-212 | EX_cpd00229_e (Glycolaldehyde -->) | Add exchange reaction for extracellular metabolite | Added for ease of media simulation | - |
| 214-222 | EX_cpd00247_e (Orotate -->) | Add exchange reaction for extracellular metabolite | Added for ease of media simulation | - |
| 224-232 | EX_cpd00266_e (Carnitine -->) | Add exchange reaction for extracellular metabolite | Added for ease of media simulation | - |
| 234-242 | EX_cpd00363_e (Ethanol -->) | Add exchange reaction for extracellular metabolite | Added for ease of media simulation | - |
| 244-252 | EX_cpd00379_e (Glutarate -->) | Add exchange reaction for extracellular metabolite | Added for ease of media simulation | - |
| 254-262 | EX_cpd00480_e (2-Dehydro-D-gluconate -->) | Add exchange reaction for extracellular metabolite | Added for ease of media simulation | - |
| 264-272 | EX_cpd00666_e (Tartrate -->) | Add exchange reaction for extracellular metabolite | Added for ease of media simulation | - |
| 274-282 | EX_cpd00794_e (TRHL -->) | Add exchange reaction for extracellular metabolite | Added for ease of media simulation | - |
| 284-292 | EX_cpd00870_e (gamma-butyrobetaine -->) | Add exchange reaction for extracellular metabolite | Added for ease of media simulation | - |
| 294-302 | EX_cpd01293_e (5-Oxoproline -->) | Add exchange reaction for extracellular metabolite | Added for ease of media simulation | - |
| 304-312 | EX_cpd01502_e (Citraconate -->) | Add exchange reaction for extracellular metabolite | Added for ease of media simulation | - |
| 314-322 | EX_cpd01630_e (cis,cis-Muconate -->) | Add exchange reaction for extracellular metabolite | Added for ease of media simulation | - |
| 324-332 | EX_cpd01947_e (BDOH -->) | Add exchange reaction for extracellular metabolite | Added for ease of media simulation | - |
| 334-342 | EX_cpd02012_e (N-Formyl-L-methionine -->) | Add exchange reaction for extracellular metabolite | Added for ease of media simulation | - |
| 344-352 | EX_cpd03294_e (Aerobactin -->) | Add exchange reaction for extracellular metabolite | Added for ease of media simulation | - |

| | | | | |
|---|---|---|---|---|
| 354-362 | EX_cpd03726_e (Fe-enterochlin -->) | Add exchange reaction for extracellular metabolite | Added for ease of media simulation | - |
| 364-372 | EX_cpd03847_e (Myristic acid -->) | Add exchange reaction for extracellular metabolite | Added for ease of media simulation | - |
| 374-382 | EX_cpd04099_e (Phosphonate -->) | Add exchange reaction for extracellular metabolite | Added for ease of media simulation | - |
| 384-392 | EX_cpd07061_e (Stachydrine -->) | Add exchange reaction for extracellular metabolite | Added for ease of media simulation | - |
| 394-402 | EX_cpd11608_e (Peptide -->) | Add exchange reaction for extracellular metabolite | Added for ease of media simulation | - |
| 404-412 | EX_cpd15901_e (2-oxohexanedioic acid -->) | Add exchange reaction for extracellular metabolite | Added for ease of media simulation | - |
| 414-422 | EX_cpd17074_e (Acetylated Alginate -->) | Add exchange reaction for extracellular metabolite | Added for ease of media simulation | - |
| 428 | cPY00016_e | Remove metabolite | Doesn't participate in any reactions, likely leftover from analysis | - |
| 429 | cPY00116_e | Remove metabolite | Replaced by cpd03587_e | - |
| 432-438 | cpd03587_e | Add metabolite | Replaces cPY00116_e | - |
| 441-442 | cpd16660_c | Correct change & formula | ModelSeed Database | - |
| 444 | cpd11595_c | Correct formula | ModelSeed Database | - |
| 446 | cpd11595_e | Correct formula | ModelSeed Database | - |
| 448-450 | cpd17016_c | Correct charge & add annotations | ModelSeed Database | - |
| 452-454 | cpd17017_c | Correct charge & add annotations | ModelSeed Database | - |
| 457-459 | rJB00226 | Change to ModelSeed ID (rxn09171) | ModelSeed Database | Jolkver, Elena, et al. "Identification and characterization of a bacterial transport system for the uptake of pyruvate, propionate, and acetate in Corynebacterium glutamicum." Journal of Bacteriology 191.3 (2009): 940. |
| 461-467 | rJB00234 | Change to ModelSeed ID (rxn10559) & add annotations | ModelSeed Database | - |
| 469-470 | rJB00235 | Change to ModelSeed ID (rxn08262) | ModelSeed Database | - |
| 472-476 | rJB00279 | Change to ModelSeed ID (rxn07967) and add annotations | ModelSeed Database | - |
| 478-482 | rPY00214 | Change to ModelSeed ID (rxn06971), update information and add metabolite | ModelSeed Database | - |
| 484-485 | rAB00001 | Change to ModelSeed ID (rxn00979) | ModelSeed Database | - |
| 487-496 | rJB00258 | Change to ModelSeed ID, change direction of reaction, add annotations | ModelSeed Database | Cooper, R. A., & Kornberg, H. L. (1964). The utilization of itaconate by Pseudomonas sp. Biochemical Journal, 91(1), 82. |
| 499 | EX_cJB00034_e | Remove reaction | ID of metabolite changed | - |
| 500 | EX_cPY00016_e | Remove reaction | ID of metabolite changed | - |
| 501 | EX_cPY00116_e | Remove reaction | ID of metabolite changed | - |
| 504-512 | EX_cpd00212_e (L-Sorbose -->) | Add exchange reaction for extracellular metabolite | Added for ease of media simulation | - |
| 514-522 | EX_cpd00138_e (D-Mannose -->) | Add exchange reaction for extracellular metabolite | Added for ease of media simulation | - |
| 524-532 | EX_cpd03587_e (kdo2-lipid a -->) | Add exchange reaction for extracellular metabolite | Added for ease of media simulation | - |
| 540-546 | cpd00477_e | Add metabolite | Added for participation in other reaction(s) | - |
| 549-556 | cpd02175_c | Add metabolite | Added for participation in other reaction(s) | - |
| 559-566 | cpd02626_c | Add metabolite | Added for participation in other reaction(s) | - |
| 569-576 | cpd00340_c | Add metabolite | Added for participation in other reaction(s) | - |
| 579-582 | cpd01715_c | Add metabolite | Added for participation in other reaction(s) | - |
| 585-593 | cpd02507_c | Add metabolite | Added for participation in other reaction(s) | - |
| 596-604 | cpd03326_c | Add metabolite | Added for participation in other reaction(s) | - |
| 607-614 | cpd03708_c | Add metabolite | Added for participation in other reaction(s) | - |
| 617-624 | cpd14700_c | Add metabolite | Added for participation in other reaction(s) | - |
| 627-635 | cpd02700_c | Add metabolite | Added for participation in other reaction(s) | - |

| | | | | |
|---|---|---|---|---|
| 638-646 | cpd02187_c | Add metabolite | Added for participation in other reaction(s) | - |
| 649-657 | cpd03572_c | Add metabolite | Added for participation in other reaction(s) | - |
| 660-668 | cpd02096_c | Add metabolite | Added for participation in other reaction(s) | - |
| 672-680 | EX_cpd00477_e (N-Acetyl-L-glutamate -->) | Add exchange reaction for extracellular metabolite | Added for ease of media simulation | - |
| 682-683 | cpd00477_e | Add annotation and fix charge | ModelSeed Database | - |
| 686-703 | rLD05146 (H2O + ATP + N-Acetyl-L-glutamate --> ADP + Phosphate + H+ + N-Acetyl-L-glutamate) | Add reaction | Added based on literature | 'Johnson DA, Tetu SG, Phillippy K, Chen J, Ren Q, Paulsen IT (2008) High-Throughput Phenotypic Characterization of Pseudomonas aeruginosa Membrane Transport Genes. Plos Genetics. 4(10): e1000211.'} |
| 706-720 | rLD05298 (N-Acetyl-L-glutamate + Na+ <=> N-Acetyl-L-glutamate + Na+) | Add reaction | Added based on literature | Johnson DA, Tetu SG, Phillippy K, Chen J, Ren Q, Paulsen IT (2008) High-Throughput Phenotypic Characterization of Pseudomonas aeruginosa Membrane Transport Genes. Plos Genetics. 4(10): e1000211 |
| 723-736 | rxn12634 (H2O + gly-glu-L --> L-Glutamate + Glycine) | Add reaction | Added based on literature | Ouidir T, Jarnier F, Cosette P, Jouenne T, Hardouin J (2015) Characterization of N-terminal protein modifications in Pseudomonas aeruginosa PA14. Journal of Proteomics 114:214-225'} |
| 739-752 | rxn12638 9H2O + gly-pro-L --> Glycine + L-Proline) | Add reaction | Added based on literature | Ouidir T, Jarnier F, Cosette P, Jouenne T, Hardouin J (2015) Characterization of N-terminal protein modifications in Pseudomonas aeruginosa PA14. Journal of Proteomics 114:214-225 |
| 755-767 | rxn02360 (trans-4-Hydroxy-L-proline <=> cis-4-Hydroxy-D-proline) | Add reaction | Added based on literature | Li, Guoqing, and Chung-Dar Lu. "Molecular characterization of LhpR in control of hydroxyproline catabolism and transport in Pseudomonas aeruginosa PAO1." Microbiology 162.7 (2016): 1232-1242. |
| 770-785 | rLD02946 (FAD + cis-4-Hydroxy-D-proline --> H+ + FADH2 + 1-Pyrroline-4-hydroxy-2-carboxylate) | Add reaction | Added based on literature | Li, Guoqing, and Chung-Dar Lu. "Molecular characterization of LhpR in control of hydroxyproline catabolism and transport in Pseudomonas aeruginosa PAO1." Microbiology 162.7 (2016): 1232-1242. |
| 788-803 | rxn01635 (H2O + H+ + 1-Pyrroline-4-hydroxy-2-carboxylate --> NH3 + 2,5-Dioxopentanoate) | Add reaction | Added based on literature | Li, Guoqing, and Chung-Dar Lu. "Molecular characterization of LhpR in control of hydroxyproline catabolism and transport in Pseudomonas aeruginosa PAO1." Microbiology 162.7 (2016): 1232-1242. |

| | | | | |
|---|---|---|---|---|
| 807-823 | rxn00196 (H2O + NADP + 2,5-Dioxopentanoate <=> NADPH + 2-Oxoglutarate + 2.0 H+) | Add reaction | Added based on literature | Li, Guoqing, and Chung-Dar Lu. "Molecular characterization of LhpR in control of hydroxyproline catabolism and transport in Pseudomonas aeruginosa PAO1." Microbiology 162.7 (2016): 1232-1242. |
| 826-837 | rxn00178 (2.0 Acetyl-CoA --> CoA + Acetoacetyl-CoA) | Add reaction | Added based on literature | Arevalo-Ferro Catalina, Hentzer M, Reil G, Gorg A, Kjelleberg S, Givskov M, Riedel K, Eberl L (2003) Identification of quorum-sensing regulated proteins in the opportunistic pathogen Pseudomonas aeruginosa by proteomics. Environmental Microbiology. 5(12):1350-1369 |
| 840-855 | rxn01725 (NAD + Carnitine <=> NADH + H+ + 3-Dehydrocarnitine) | Add reaction | Added based on literature | Meadows, Jamie A., and Matthew J. Wargo. "Transcriptional regulation of Carnitine catabolism in Pseudomonas aeruginosa by CdhR." MSphere 3.1 (2018). |
| 857-866 | cLD00540_c | Add metabolite | Added based on literature | Meadows, Jamie A., and Matthew J. Wargo. "Transcriptional regulation of Carnitine catabolism in Pseudomonas aeruginosa by CdhR." MSphere 3.1 (2018). |
| 869-884 | rLD01726 (Acetyl-CoA + 3-Dehydrocarnitine <=> BET-CoA + H+ + Acetoacetate) | Add reaction | Added based on literature | Meadows, Jamie A., and Matthew J. Wargo. "Transcriptional regulation of Carnitine catabolism in Pseudomonas aeruginosa by CdhR." MSphere 3.1 (2018)., Wargo, Matthew J., and Deborah A. Hogan. "Identification of genes required for Pseudomonas aeruginosa carnitine catabolism." Microbiology 155.Pt 7 (2009): 2411., Bastard, Karine, et al. "Revealing the hidden functional diversity of an enzyme family." Nature chemical biology 10.1 (2014): 42-49. |
| 887-900 | rLD01727 (BET-CoA + H2O <=> CoA + BET) | Add reaction | Added based on literature | Meadows, Jamie A., and Matthew J. Wargo. "Transcriptional regulation of Carnitine catabolism in Pseudomonas aeruginosa by CdhR." MSphere 3.1 (2018)., Wargo, Matthew J., and Deborah A. Hogan. "Identification of genes required for Pseudomonas aeruginosa carnitine catabolism." Microbiology 155.Pt 7 (2009): 2411., Bastard, Karine, et al. "Revealing the hidden functional diversity of an enzyme family." Nature chemical biology 10.1 (2014): 42-49. |

| | | | | |
|---|---|---|---|---|
| 903-908 | rxn02028 | Update reaction GPR | Added based on literature | Meadows, Jamie A., and Matthew J. Wargo. "Transcriptional regulation of Carnitine catabolism in Pseudomonas aeruginosa by CdhR." MSphere 3.1 (2018). |
| 910-926 | rxn12739 (H2O + FAD + Dimethylglycine <=> Formaldehyde + Sarcosine + FADH2) | Add reaction | Added based on literature | Meadows, Jamie A., and Matthew J. Wargo. "Transcriptional regulation of Carnitine catabolism in Pseudomonas aeruginosa by CdhR." MSphere 3.1 (2018). Wargo, Matthew J., Benjamin S. Szwergold, and Deborah A. Hogan. "Identification of two gene clusters and a transcriptional regulator required for Pseudomonas aeruginosa glycine betaine catabolism." Journal of bacteriology 190.8 (2008): 2690. |
| 929-940 | rxn03040 (5-Carboxymethyl-2-hydroxymuconate <=> 5-Carboxy-2-oxohept-3-enedioate) | Add reaction | Added based on genetic evidence | - |
| 943-956 | rxn03041 (H+ + 5-Carboxy-2-oxohept-3-enedioate --> CO2 + 2-Hydroxyhepta-2,4-dienedioate) | Add reaction | Added based on genetic evidence | - |
| 960-972 | rxn04706 (H2O + 2-Hydroxyhepta-2,4-dienedioate <=> 2,4-Dihydroxyhept-2-enedioate) | Add reaction | Added based on genetic evidence | - |
| 975-987 | rxn01203 (Pyruvate + 4-Oxobutanoate <-- 2,4-Dihydroxyhept-2-enedioate) | Add reaction | Added based on genetic evidence | - |
| 990-992 | rxn00670 | Open reaction bounds | Bounds were set to 0 for unknown reason | - |
| 995-997 | rxn00985 | Open reaction bounds | Bounds were set to 0 for unknown reason | - |
| 1000-1001 | rxn00166 | Add note to reaction to explain why this reaction makes D-serine carbon source prediction incorrect | - | Li, Guoqing, and Chung-Dar Lu. "The cryptic dsdA gene encodes a functional D-Serine dehydratase in Pseudomonas aeruginosa PAO1." Current microbiology 72.6 (2016): 788-794. |
| 1004-1018 | rxn07430 (TPP + H+ + 3-Methyl-2-oxobutanoate --> CO2 + 2-Methyl-1-hydroxypropyl-TPP) | Add reaction | Added based on genetic evidence | - |
| 1021-1034 | rxn07431 (Lipoamide + 2-Methyl-1-hydroxypropyl-TPP <=> TPP + S-(2-Methylpropionyl)-dihydrolipoamide) | Add reaction | Added based on genetic evidence | - |
| 1037-1050 | rxn01925 (Dihydrolipoamide + Isobutyryl-CoA <=> CoA + S-(2-Methylpropionyl)-dihydrolipoamide) | Add reaction | Added based on genetic evidence | - |
| 1053-1066 | rxn01924 (FAD + Isobutyryl-CoA <=> FADH2 + Methacrylyl-CoA) | Add reaction | Added based on genetic evidence | - |
| 1069-1081 | rxn02949 (H2O + Methacrylyl-CoA <=> (S)-3-Hydroxyisobutyryl-CoA) | Add reaction | Added based on genetic evidence | - |
| 1084-1097 | rxn02925 (L-Alanine + 3-Oxo-2-methylpropanoate <=> Pyruvate + L-3-Amino-isobutyrate) | Add reaction | Added based on genetic evidence | - |
| 1100-1106 | rxn00868 | Flip directionality of reaction | Now matches ModelSeed database | - |

| | | | | |
|---|---|---|---|---|
| 1110 | rxn00160 | Remove reaction | Was added from P. Putida model, but goes against biolog predictions | - |
| 1113 | rxn01355 | Remove PA14_39190 from reaction | Was found to be non essential in vitro | - |
| 1116 | rxn05155 | Add EC code to annotation | Based on KEGG | - |
| 1120-1153 | PA14_17050, PA14_28590, PA14_47840, PA14_47860, PA14_48000, PA14_36050, PA14_48010, PA14_31530, PA14_13090, PA14_63250, PA14_71140, PA14_71420, PA14_71410, PA14_71280, PA14_71260, PA14_10610, PA14_39100, PA14_10650, PA14_10640, PA14_10590, PA14_10570, PA14_35520, PA14_35500, PA14_35970, PA14_44590, PA14_51120, PA14_66040, PA14_49080, PA14_19740, PA14_40980, PA14_70160, PA14_48440, PA14_51990, PA14_37340 | Add SBO annotations for genes | Based on Memote report | - |
| 1156-1189 | PA14_48440, PA14_51990, PA14_37340, PA14_28590, PA14_17050, PA14_47840, PA14_47860, PA14_48000, PA14_48010, PA14_36050, PA14_13090, PA14_63250, PA14_31530, PA14_71140, PA14_71410, PA14_71420, PA14_71260, PA14_71280, PA14_10610, PA14_39100, PA14_10650, PA14_10640, PA14_10590, PA14_10570, PA14_35520, PA14_35500, PA14_66040, PA14_35970, PA14_51120, PA14_49080, PA14_44590, PA14_40980, PA14_19740, PA14_70160 | Add Kegg annotations for genes | Based on Memote report | - |
| 1192 | ATPM | Correct SBO number on reaction | Based on Memote report | - |
| 1195 | cJB00125_c | Remove incorrect annotation | - | - |
| 1198-1202 | Various reactions | Remove mislabeled annotations | - | - |
| 1208 | cpd01700_c | Correct formula to fix rxn02749 imbalance | Based on ModelSeed | - |
| 1211-1215 | rxn13839 | Fix reaction imbalance by correcting metabolite formulas/charges and adding missing metabolite | Based on ModelSeed | - |
| 1218-1219 | rxn11544 | Fix reaction imbalance | Based on ModelSeed | - |
| 1222 | rxn03512 | Fix reaction imbalance | Based on ModelSeed | - |
| 1225-1226 | rxn00105 | Fix reaction imbalance | Based on ModelSeed | - |
| 1229-1262 | Various reactions/metabolites | Fix dates in notes | - | - |
| 1267-1274 | cpd15666_c | Add Peptidoglycan polymer (n-1 subunits) metabolite | Based on ModelSeed | - |
| 1277-1289 | rxn10194, rxn10195, rxn10196, rxn10199 | Correct reaction imbalances by adding cpd15666_c from reactions (and removing cpd15665_c from some) | - | - |
| 1292-1303 | rxn12751 (Undecaprenyl-diphospho-N-acetylmuramoyl--N-acetylglucosamine-L-ala-D-glu-meso-2-6-diaminopimeloyl-D-ala-D-ala <=> Bactoprenyl diphosphate + Peptidoglycan polymer (n-1 subunits)) | Add synthesis rxns for cpd15666_c | From P. putida KT2440 model | - |
| 1308-1326 | cpd00067_p | Add periplasmic hydrogen to model | Allows reconstruction to be more biologically accurate | - |

| | | | | |
|---|---|---|---|---|
| 1328-1357 | rxn10042, rJB00264, rxn13726, rxn13689, rJB00271 | Change extracellular hydrogen to periplasmic hydrogen in reactions associated with the electron transport chain (ETC) in order to assist in fixing ATP generating loop | Allows reconstruction to be more biologically accurate | - |
| 1362-1399 | rxn00251, rxn13919, rxn05145, rxn05158, rxn01452, rxn00097, rxn00161, rxn08066, rxn13842 | Make minimal changes to reactions involved in ATP-generating loop in order to resolve the loop | Loop was allowing model to generate ATP without cost, which is not feasible biologically | - |
| 1404-1405 | rxn00178 | Make irreversible to correct L-valine and acetoacetate growth predictions | Reaction belongs in reconstruction but the addition of it made carbon source predictions incorrect | - |
| 1409-1411 | PA14_68390 | Add gene to model and add it to associated reaction (rxn00708) | Based on KEGG | - |
| 1416-1518 | Various metabolites and reactions | Add metabolites and exchange reactions that are required for carbon source prediction testing | - | - |
| 1588-1601 | rxn00994 | Add reaction | Based on Blastn evidence | - |
| 1605-1618 | rxn00875 | Add reaction | Based on Blastn evidence | - |
| 1621-1639 | SPONTANEOUS, unassigned, Unassigned (GPRs) | Remove GPRs from model | Do not represent actual genes | - |
| 1644-1845 | PA14_Biomass | Create macromolecular bins for metabolites (protein_c, rna_c, dna_c, lipid_c) and organize the biomass OF into these categories. Add biomass metabolite (cpd11416_c), create biomass sink (SK_cpd11416_c), and add LPS (cpd17065_c) and biomass metabolite to biomass objective function | Organized BOF for ease of analysis, added LPS to BOF based on literature | Darveau, R.P., and R.E. Hancock. 1983. Procedure for isolation of bacterial lipopolysaccharides from both smooth and rough Pseudomonas aeruginosa and Salmonella typhimurium strains. Journal of Bacteriology. 155(2):831. |
| 1847-1851 | Model Object | Update model metadata | added id, name, taxonomy, and creators of model | - |
| 1853-1856 | Model media | Set lower bound of all exchanges to 0 and upper bound to 1000 | - | - |

Note: "Code Line" corresponds to "Correcting_mass_charge_imbalances.ipynb"

| Code Line | Model Object | Change | Justification |
|---|---|---|---|
| 7 | rxn10122 | stoichiometric coefficients | Based on ModelSeed |
| 9 | cpd17051_c | Chemical formula | Based on ModelSeed residue from cpd14938 |
| 11 | cpd17097_c | Chemical formula and charge | Balance based on participating metabolite cpd11669 |
| 13 | rAB00001 | Add proton | Based on MetaCyc |
| 15 | rxn13896 | Add proton | Based on MetaCyc |
| 17 | rxn00190 | Add proton | Based on MetaCyc |
| 19 | rxn00138 | Add proton | Based on MetaCyc |
| 21 | rxn13788 | Add proton | Based on MetaCyc |
| 23 | rxn10215 | Add proton | Based on ModelSeed |
| 25 | rxn10218 | Add proton | Based on ModelSeed |
| 27 | rxn10214 | Add proton | Based on ModelSeed |
| 29 | rxn10219 | Add proton | Based on ModelSeed |
| 31 | rxn10217 | Add proton | Based on ModelSeed |
| 33 | rxn10216 | Add proton | Based on ModelSeed |
| 35 | rxn10212 | Add proton | Based on previous reactions (e.g., rxn10216) |
| 37 | rxn10213 | Add proton | Based on previous reactions (e.g., rxn10216) |
| 39 | rxn10221 | Add proton | Based on ModelSeed |

| | | | |
|---|---|---|---|
| 41 | rxn10224 | Add proton | Based on ModelSeed |
| 43 | rxn08306 | Add proton | Based on ModelSeed |
| 45 | rxn08307 | Add proton | Based on ModelSeed |
| 47 | rxn08308 | Add proton | Based on ModelSeed |
| 49 | rxn08310 | Add proton | Based on ModelSeed |
| 51 | rxn08311 | Add proton | Based on ModelSeed |
| 53 | rxn08312 | Add proton | Based on ModelSeed |
| 55 | rxn10220 | Add proton | Based on ModelSeed |
| 57 | rxn10225 | Add proton | Based on ModelSeed |
| 59 | rxn10223 | Add proton | Based on ModelSeed |
| 61 | rxn10222 | Add proton | Based on ModelSeed |
| 63 | rxn00986 | Set reaction direction, remove proton | Based on ModelSeed |
| 65 | rxn00416 | Add proton | Based on MetaCyc |
| 67 | rxn11080 | Add water and proton | Based on ModelSeed |
| 69 | rxn00065 | Add proton | Based on ModelSeed |
| 71 | rxn13877 | Add proton | Based on ModelSeed |
| 73 | rxn08309 | Add proton | Based on ModelSeed |
| 75 | rPY00163 | Remove proton | Inferred from imbalance |
| 77 | rPY00168 | Remove proton | Inferred from imbalance |
| 79 | rxn00300 | Add proton | Based on MetaCyc |
| 81 | rxn06280 | Add proton | Based on ModelSeed |
| 83 | rxn09240 | Add proton | Based on ModelSeed |
| 85 | rxn12848 | Add proton | Based on ModelSeed |
| 87 | rxn12849 | Add proton | Based on ModelSeed |
| 89 | rxn12850 | Add proton | Based on ModelSeed |
| 91 | rxn12851 | Add proton | Based on ModelSeed |
| 93 | rxn00711 | Add proton | Based on ModelSeed |
| 95 | rxn06439 | Add proton | Based on ModelSeed |
| 97 | rxn13804 | Change ubiquinone to ubiquinone-8 | Based on MetaCyc |
| 99 | cpd01015_e | Change charge | Based on ModelSeed |
| 103 | rxn00126 | Add proton | Based on ModelSeed |
| 105 | rxn00791 | Add proton | Based on ModelSeed |
| 107 | rxn00670 | Remove proton | Based on ModelSeed |
| 109 | rxn13904 | Add proton | Based on ModelSeed |
| 111 | rxn10090 | Add proton | Based on ModelSeed |
| 111 | cpd03422_c | Change charge | Inferred from imbalance |
| 113 | rxn11544 | Add proton | Based on ModelSeed |
| 115 | rxn08707 | Add proton | Based on ModelSeed |
| 117 | rxn00139 | Add proton | Based on ModelSeed |
| 119 | rxn06831 | Add proton | Based on ModelSeed |
| 119 | cpd11451_c | Change formula | Based on MetaCyc |
| 121 | cpd00155_c | Change formula and charge | Based on BiGG |
| 124 | rxn13817 | Change stoichiometric coefficients | Based on MetaCyc |
| 126 | rxn13815 | Change stoichiometric coefficients | Based on MetaCyc |
| 126 | rxn13815 | Remove proton | Based on KEGG |
| 128 | rxn06447 | Add proton | Based on ModelSeed and e.g., reaction rxn06439 |
| 130 | rxn05451 | Add proton | Based on ModelSeed and imbalance |
| 132 | rxn05455 | Add proton | Based on ModelSeed and imbalance |

| | | | |
|---|---|---|---|
| 134 | rxn05453 | Add proton | Based on ModelSeed and imbalance |
| 136 | rxn05452 | Add proton | Based on ModelSeed and imbalance |
| 138 | rxn05456 | Add proton | Based on ModelSeed and imbalance |
| 140 | rxn05454 | Add proton | Based on ModelSeed and imbalance |
| 142 | rxn05358 | Add proton | Based on ModelSeed |
| 144 | rxn05408 | Add proton | Based on ModelSeed |
| 146 | rxn05383 | Add proton | Based on ModelSeed |
| 148 | rxn05459 | Add proton | Based on ModelSeed |
| 150 | rxn05457 | Add proton | Based on other reactions, e.g., rxn05459 |
| 152 | rxn02405 | Add proton | Based on ModelSeed |
| 154 | rxn00790 | Add proton | Based on ModelSeed |
| 156 | rxn06937 | Remove proton | Based on BiGG |
| 158 | cpd17048_c | Change charge | Inferred from participating metabolite cpd00869 |
| 160 | cpd17079_c | Change charge | Inferred from participating metabolite cpd00869 |
| 162 | cpd17052_c | Change charge | Inferred from participating metabolite cpd00870 |
| 164 | cJB00124_c | Change formula | Inferred from imbalance |
| 166 | rxn13864 | Change ubiquinone to ubiquinone-8 | Based on MetaCyc |
| 167 | cpd00659_c | Change formula and charge | Based on ModelSeed |
| 169 | rxn13903 | Add proton | Based on ModelSeed |
| 173 | rxn06443 | Remove proton | Based on MetaCyc |
| 175 | rxn01434 | Add proton | Based on MetaCyc |
| 177 | rxn13874 | Remove proton | Based on ModelSeed |
| 177 | cpd17067_c | Change charge | Inferred from imbalance |
| 179 | rxn13889 | Change stoichiometric coefficients | Based on MetaCyc |
| 182 | rxn05030 | Add proton | Based on ModelSeed |
| 184 | cpd00134_c | Change formula and charge | Based on ModelSeed |
| 187 | rxn00947 | Remove proton | Based on MetaCyc |
| 189 | cpd03126_c | Change formula and charge | Based on ModelSeed |
| 191 | cpd03113_c | Change formula and charge | Based on ModelSeed |
| 193 | cpd03114_c | Change formula and charge | Based on ModelSeed |
| 195 | rxn00917 | Add proton | Based on MetaCyc |
| 197 | rxn06441 | Remove proton | Based on MetaCyc |
| 201 | rxn01466 | Add proton | Based on ModelSeed |
| 203 | rxn13826 | Add proton | Based on previous reactions (e.g., rxn01466) |
| 205 | rxn13887 | Change stoichiometric coefficients | Based on MetaCyc |
| 208 | rxn00792 | Change reaction directionality, remove proton | Based on ModelSeed |
| 210 | rxn06448 | Remove proton | Based on MetaCyc |
| 212 | rxn00789 | Add proton | Based on ModelSeed |
| 214 | rxn02402 | Remove proton | Based on ModelSeed |
| 216 | rxn00100 | Add proton | Based on ModelSeed |
| 218 | rxn09037 | Add protons | Based on ModelSeed |
| 220 | rxn05028 | Add proton | Based on ModelSeed |
| 222 | rxn00915 | Add proton | Based on ModelSeed |
| 224 | rxn00836 | Add proton | Based on ModelSeed |
| 226 | rxn13816 | Change stoichiometric coefficients | Based on MetaCyc |
| 229 | rxn01791 | Add proton | Based on MetaCyc |

| | | | |
|---|---|---|---|
| 231 | rxn13894 | Change stoichiometric coefficients | Based on MetaCyc |
| 234 | rxn06432 | Remove proton | Based on MetaCyc |
| 236 | rxn12879 | Add proton | Based on ModelSeed |
| 238 | rxn02834 | Add proton | Based on MetaCyc |
| 240 | rxn13108 | Remove oxygen | Inferred from imbalance |
| 242 | rxn09177 | Add proton | Based on MetaCyc |
| 244 | rxn01519 | Add proton | Based on MetaCyc |
| 246 | rxn01362 | Add proton | Based on ModelSeed |
| 248 | rxn03893 | Add proton | Based on ModelSeed |
| 250 | rxn05034 | Add proton | Based on ModelSeed |
| 252 | rxn13818 | Change stoichiometric coefficients | Based on MetaCyc |
| 255 | rxn13899 | Add protons | Based on ModelSeed and inferred from imbalance |
| 257 | rxn13900 | Add protons | Based on ModelSeed and inferred from imbalance |
| 259 | rxn13901 | Add protons | Based on ModelSeed and inferred from imbalance |
| 261 | rxn13868 | Add protons | Based on ModelSeed |
| 263 | rxn13876 | Add proton | Based on ModelSeed |
| 265 | rxn01093 | Remove proton | Based on ModelSeed |
| 267 | rxn05118 | Remove proton | Based on ModelSeed |
| 269 | rxn13921 | Add proton | Based on ModelSeed |
| 271 | rxn13825 | Remove proton | Based on ModelSeed |
| 273 | rxn00695 | Remove proton | Based on ModelSeed |
| 275 | rxn05964 | Add proton | Based on ModelSeed |
| 277 | cpd11611_c | Change formula and charge | Based on ModelSeed |
| 280 | rxn05824 | Add proton | Based on ModelSeed |
| 282 | rxn00735 | Remove proton | Based on ModelSeed |
| 284 | rxn01208 | Add proton | Based on ModelSeed |
