## Supplemental Data 2 for "An updated genome-scale metabolic network reconstruction of *Pseudomonas aeruginosa* PA14 to characterize mucin-driven shifts in bacterial metabolism"

iPau21 MEMOTE Report Follows

#### Independent Section

Contains tests that are independent of the class of modeled organism, a model's complexity or types of identifiers that are

##### Consistency

| | | |
|---|---|---|
| Stoichiometric Consistency | 99.2% | X3 |
| Mass Balance | 91.2% | |
| Charge Balance | 89.0% | |
| Metabolite Connectivity | 100.0% | |
| Unbounded Flux In Default Medium | 75.2% | |
| Sub Total | 93% | X3 |

##### Annotation - Metabolites

| | |
|---|---|
| Presence of Metabolite Annotation | 100.0% |
| Metabolite Annotations Per Database | Info |
| pubchem.compound | 0.0% |
| kegg.compound | 52.7% |
| seed.compound | 80.7% |
| inchikey | 0.0% |
| inchi | 0.0% |
| chebi | 53.3% |
| hmdb | 43.0% |

#### Specific Section

Covers general statistics and specific aspects of a metabolic network that are not universally applicable. See readme for

##### SBML

| | |
|---|---|
| SBML Level and Version | Errored |
| FBC enabled | Errored |

##### Basic Information

| | |
|---|---|
| Model Identifier | iPAU |
| Total Metabolites | 1,310 |
| Total Reactions | 1,571 |
| Total Genes | 1,171 |
| Total Compartments | 3 |
| Metabolic Coverage | 1.34 |

##### Metabolite Information

| | |
|---|---|
| Unique Metabolites | 1,310 |
| Duplicate Metabolites in Identical Compartments | 0 |
| Metabolites without Charge | 0 |
| Metabolites without Formula | 0 |
| Medium Components | 36 |

| | |
|---|---|
| bigg.metabolite | 81.5% |
| biocyc | 51.0% |
| **Metabolite Annotation Conformity Per Database** | Info |
| pubchem.compound | 0.0% |
| kegg.compound | 100.0% |
| seed.compound | 100.0% |
| inchikey | 0.0% |
| inchi | 0.0% |
| chebi | 100.0% |
| hmdb | 100.0% |
| reactome | 0.0% |
| metanetx.chemical | 100.0% |
| bigg.metabolite | 99.9% |
| biocyc | 100.0% |
| **Uniform Metabolite Identifier Namespace** | 100.0% |
| **Sub Total** | 75% |

#### Annotation - Reactions

| | |
|---|---|
| **Presence of Reaction Annotation** | 100.0% |
| **Reaction Annotations Per Database** | Info |
| rhea | 29.9% |

[illegible]

| | |
|---|---|
| Purely Metabolic Reactions with Constraints | 12 |
| Transport Reactions | 246 |
| Transport Reactions with Constraints | 1 |
| Thermodynamic Reversibility of Purely Metabolic Reactions | 0.22 |
| Reactions With Partially Identical Annotations | 0.00 |
| Duplicate Reactions | 0.00 |
| Reactions With Identical Genes | 0.47 |

#### Gene-Protein-Reaction (GPR) Associations

| | |
|---|---|
| Reactions without GPR | 52 |
| Fraction of Transport Reactions without GPR | 0.04 |
| Enzyme Complexes | 194 |

#### Biomass

| | |
|---|---|
| Biomass Reactions Identified | 1 |
| Biomass Consistency | Errored |
| Biomass Production In Default Medium | 3.64 |
| Unrealistic Growth Rate In Default Medium | true |
| Biomass Production In Complete Medium | 159.68 |

[illegible]

| | |
|---|---|
| metanetx.reaction | 45.5% |
| bigg.reaction | 50.4% |
| reactome | 0.0% |
| ec-code | 64.2% |
| brenda | 0.0% |
| biocyc | 30.6% |
| **Reaction Annotation Conformity Per Database** | **Info** |
| rhea | 100.0% |
| kegg.reaction | 100.0% |
| seed.reaction | 89.1% |
| metanetx.reaction | 100.0% |
| bigg.reaction | 100.0% |
| reactome | 0.0% |
| ec-code | 98.9% |
| brenda | 0.0% |
| biocyc | 100.0% |
| **Uniform Reaction Identifier Namespace** | 100.0% |
| **Sub Total** | 79% |

#### Annotation - Genes

| | |
|---|---|
| **Presence of Gene Annotation** | 100.0% |

| | |
|---|---|
| **Ratio of Direct Metabolites in Biomass Reaction** | 0.00 |
| **Number of Missing Essential Biomass Precursors** | 15 |

#### Energy Metabolism

| | |
|---|---|
| **Non-Growth Associated Maintenance Reaction** | 1 |
| **Growth-associated Maintenance in Biomass Reaction** | true |
| **Number of Reversible Oxygen-Containing Reactions** | 12 |
| **Erroneous Energy-generating Cycles** | **Info** |
| MNXM3 | Skipped |
| MNXM63 | Skipped |
| MNXM51 | Skipped |
| MNXM121 | Skipped |
| MNXM423 | Skipped |
| MNXM6 | Skipped |
| MNXM10 | Skipped |
| MNXM38 | Skipped |
| MNXM208 | Skipped |
| MNXM191 | Skipped |
| MNXM223 | Skipped |
| MNXM7517 | Skipped |
| MNXM12233 | Skipped |

| | |
|---|---|
| uniprot | 96.8% |
| ecogene | 0.0% |
| kegg.genes | 100.0% |
| ncbigi | 0.0% |
| ncbigene | 96.8% |
| ncbiprotein | 96.8% |
| ccds | 0.0% |
| hprd | 0.0% |
| asap | 96.8% |

**Gene Annotation Conformity Per Database** Info

| | |
|---|---|
| refseq | 100.0% |
| uniprot | 100.0% |
| ecogene | 0.0% |
| kegg.genes | 100.0% |
| ncbigi | 0.0% |
| ncbigene | 100.0% |
| ncbiprotein | 100.0% |
| ccds | 0.0% |
| hprd | 0.0% |
| asap | 100.0% |

**Sub Total** 73%

| | |
|---|---|
| MNXM89557 | Skipped |

#### Network Topology

| | |
|---|---|
| Universally Blocked Reactions | 606 |
| Orphan Metabolites | 45 |
| Dead-end Metabolites | 79 |
| Stoichiometrically Balanced Cycles | 279 |
| Metabolite Production In Complete Medium | 295 |
| Metabolite Consumption In Complete Medium | 549 |

#### Matrix Conditioning

| | |
|---|---|
| Ratio Min/Max Non-Zero Coefficients | 0.00 |
| Independent Conservation Relations | 55 |
| Rank | 1255 |
| Degrees Of Freedom | 316 |

#### Experimental Data Comparison

| | |
|---|---|
| Growth Prediction | Skipped |
| Gene Essentiality Prediction | Skipped |

#### Misc. Tests

| | |
|---|---|
| Platform | Linux |
| Memote Version | 0.10.2 |

| | |
|---|---|
| Metabolite SBO:0000247 Presence | 100.0% |
| Reaction General SBO Presence | 100.0% |
| Metabolic Reaction SBO:0000176 Presence | 99.9% |
| Transport Reaction SBO:0000185 Presence | 100.0% |
| Exchange Reaction SBO:0000627 Presence | 100.0% |
| Demand Reaction SBO:0000628 Presence | Skipped |
| Sink Reactions SBO:0000632 Presence | Skipped |
| Gene General SBO Presence | 100.0% |
| Gene SBO:0000243 Presence | 100.0% |
| Biomass Reactions SBO:0000629 Presence | 100.0% |
| Sub Total | 82% X2 |
| Total Score | 85% |

Total Score

# 85%

Score per Category

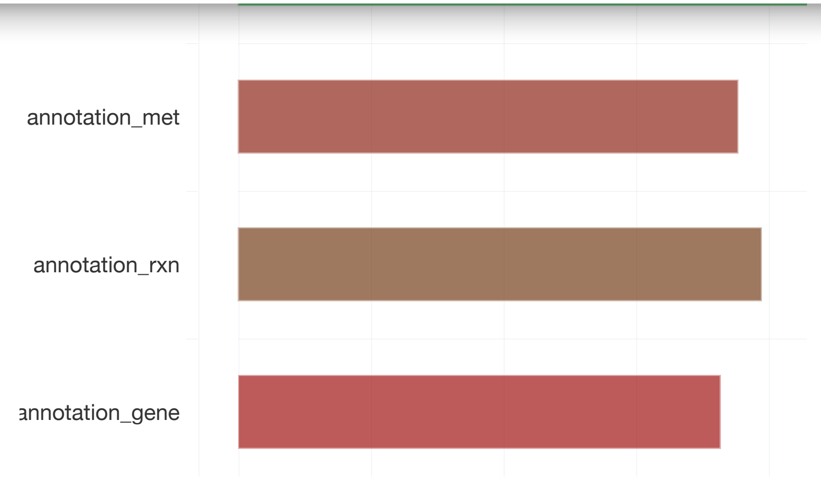
annotation_met
annotation_rxn
annotation_gene

iPau1129 MEMOTE Report Follows

Expand All Readme 2020-07-22 14:30

### Independent Section

Contains tests that are independent of the class of modeled organism, a model's complexity or types of identifiers that are used

#### Consistency

| | | |
|---|---|---|
| Stoichiometric Consistency | 36.2% | X3 |
| Mass Balance | 92.4% | |
| Charge Balance | 91.1% | |
| Metabolite Connectivity | 100.0% | |
| Unbounded Flux In Default Medium | 72.6% | |
| Sub Total | 66% | X3 |

#### Annotation - Metabolites

| | |
|---|---|
| Presence of Metabolite Annotation | 0.0% |
| Metabolite Annotations Per Database | Info |
| pubchem.compound | 0.0% |
| kegg.compound | 0.0% |
| seed.compound | 0.0% |
| inchikey | 0.0% |
| inchi | 0.0% |
| chebi | 0.0% |
| hmdb | 0.0% |
| reactome | 0.0% |

### Specific Section

Covers general statistics and specific aspects of a metabolic network that are not universally applicable. See readme for more

#### SBML

| | |
|---|---|
| SBML Level and Version | Errored |
| FBC enabled | Errored |

#### Basic Information

| | |
|---|---|
| Model Identifier | |
| Total Metabolites | 1,286 |
| Total Reactions | 1,495 |
| Total Genes | 1,132 |
| Total Compartments | 2 |
| Metabolic Coverage | 1.32 |

#### Metabolite Information

| | |
|---|---|
| Unique Metabolites | 1,099 |
| Duplicate Metabolites in Identical Compartments | 0 |
| Metabolites without Charge | 3 |
| Metabolites without Formula | 5 |
| Medium Components | 33 |

#### Reaction Information

| | |
|---|---|
| biocyc | 0.0% |
| **Metabolite Annotation Conformity Per Database** | Info |
| pubchem.compound | 0.0% |
| kegg.compound | 0.0% |
| seed.compound | 0.0% |
| inchikey | 0.0% |
| inchi | 0.0% |
| chebi | 0.0% |
| hmdb | 0.0% |
| reactome | 0.0% |
| metanetx.chemical | 0.0% |
| bigg.metabolite | 0.0% |
| biocyc | 0.0% |
| **Uniform Metabolite Identifier Namespace** | 100.0% |
| **Sub Total** | 25% |

#### Annotation - Reactions

| | |
|---|---|
| **Presence of Reaction Annotation** | 0.0% |
| **Reaction Annotations Per Database** | Info |
| rhea | 0.0% |
| kegg.reaction | 0.0% |
| seed.reaction | 0.0% |
| metanetx.reaction | 0.0% |

| | |
|---|---|
| **Transport Reactions** | 243 |
| **Transport Reactions with Constraints** | 1 |
| **Thermodynamic Reversibility of Purely Metabolic Reactions** | 1.00 |
| **Reactions With Partially Identical Annotations** | 0.00 |
| **Duplicate Reactions** | 0.00 |
| **Reactions With Identical Genes** | 0.48 |

#### Gene-Protein-Reaction (GPR) Associations

| | |
|---|---|
| **Reactions without GPR** | 44 |
| **Fraction of Transport Reactions without GPR** | 0.04 |
| **Enzyme Complexes** | 186 |

#### Biomass

| | |
|---|---|
| **Biomass Reactions Identified** | 0 |
| **Biomass Consistency** | Skipped |
| **Biomass Production In Default Medium** | Skipped |
| **Unrealistic Growth Rate In Default Medium** | Skipped |
| **Biomass Production In Complete Medium** | Skipped |
| **Blocked Biomass Precursors In Default Medium** | Skipped |
| **Blocked Biomass Precursors In Complete Medium** | Skipped |
| **Ratio of Direct Metabolites in Biomass Reaction** | Skipped |

| | |
|---|---|
| ec-code | 0.0% |
| brenda | 0.0% |
| biocyc | 0.0% |
| **Reaction Annotation Conformity Per Database** | Info |
| rhea | 0.0% |
| kegg.reaction | 0.0% |
| seed.reaction | 0.0% |
| metanetx.reaction | 0.0% |
| bigg.reaction | 0.0% |
| reactome | 0.0% |
| ec-code | 0.0% |
| brenda | 0.0% |
| biocyc | 0.0% |
| **Uniform Reaction Identifier Namespace** | 100.0% |
| **Sub Total** | 25% |

#### Annotation - Genes

| | |
|---|---|
| **Presence of Gene Annotation** | 0.0% |
| **Gene Annotations Per Database** | Info |
| refseq | 0.0% |
| uniprot | 0.0% |
| ecogene | 0.0% |
| kegg.genes | 0.0% |

#### Energy Metabolism

| | |
|---|---|
| **Non-Growth Associated Maintenance Reaction** | 1 |
| **Growth-associated Maintenance in Biomass Reaction** | Skipped |
| **Number of Reversible Oxygen-Containing Reactions** | 12 |
| **Erroneous Energy-generating Cycles** | Info |
| MNXM3 | Skipped |
| MNXM63 | Skipped |
| MNXM51 | Skipped |
| MNXM121 | Skipped |
| MNXM423 | Skipped |
| MNXM6 | Skipped |
| MNXM10 | Skipped |
| MNXM38 | Skipped |
| MNXM208 | Skipped |
| MNXM191 | Skipped |
| MNXM223 | Skipped |
| MNXM7517 | Skipped |
| MNXM12233 | Skipped |
| MNXM558 | Skipped |
| MNXM21 | Skipped |
| MNXM89557 | Skipped |

| | |
|---|---|
| ncbiprotein | 0.0% |
| ccds | 0.0% |
| hprd | 0.0% |
| asap | 0.0% |
| **Gene Annotation Conformity Per Database** | Info |
| refseq | 0.0% |
| uniprot | 0.0% |
| ecogene | 0.0% |
| kegg.genes | 0.0% |
| ncbigi | 0.0% |
| ncbigene | 0.0% |
| ncbiprotein | 0.0% |
| ccds | 0.0% |
| hprd | 0.0% |
| asap | 0.0% |
| **Sub Total** | 0% |

#### Annotation - SBO Terms

| | |
|---|---|
| Metabolite General SBO Presence | 0.0% |
| Metabolite SBO:0000247 Presence | 0.0% |
| Reaction General SBO Presence | 0.0% |
| Metabolic Reaction SBO:0000176 Presence | 0.0% |
| Transport Reaction SBO:0000185 Presence | 0.0% |

| | |
|---|---|
| Orphan Metabolites | 56 |
| Dead-end Metabolites | 86 |
| Stoichiometrically Balanced Cycles | 253 |
| Metabolite Production In Complete Medium | 353 |
| Metabolite Consumption In Complete Medium | 580 |

#### Matrix Conditioning

| | |
|---|---|
| Ratio Min/Max Non-Zero Coefficients | 0.00 |
| Independent Conservation Relations | 69 |
| Rank | 1217 |
| Degrees Of Freedom | 278 |

#### Experimental Data Comparison

| | |
|---|---|
| Growth Prediction | Skipped |
| Gene Essentiality Prediction | Skipped |

#### Misc. Tests

#### Environment

| | |
|---|---|
| Python Version | 3.6.10 |
| Platform | Linux |
| Memote Version | 0.10.2 |

| | | |
|---|---|---|
| Sink Reactions SBO:0000632 Presence | Skipped | |
| Gene General SBO Presence | 0.0% | |
| Gene SBO:0000243 Presence | 0.0% | |
| Biomass Reactions SBO:0000629 Presence | Skipped | |
| **Sub Total** | 0% | X2 |
| **Total Score** | 30% | |

Total Score

# 30%

Score per Category

iML1515 MEMOTE Report Follows

### Independent Section

Contains tests that are independent of the class of modeled organism, a model's complexity or types of identifiers that are

#### Consistency

| | | |
|---|---|---|
| Stoichiometric Consistency | 100.0% | x3 |
| Mass Balance | 100.0% | |
| Charge Balance | 100.0% | |
| Metabolite Connectivity | 100.0% | |
| Unbounded Flux In Default Medium | 88.9% | |
| Sub Total | 98% | x3 |

#### Annotation - Metabolites

| | |
|---|---|
| Presence of Metabolite Annotation | 100.0% |
| Metabolite Annotations Per Database | Info |
| pubchem.compound | 0.0% |
| kegg.compound | 71.9% |
| seed.compound | 85.3% |
| inchikey | 0.0% |
| inchi | 0.0% |
| chebi | 76.7% |
| hmdb | 56.5% |

### Specific Section

Covers general statistics and specific aspects of a metabolic network that are not universally applicable. See readme for

#### SBML

| | |
|---|---|
| SBML Level and Version | Errored |
| FBC enabled | Errored |

#### Basic Information

| | |
|---|---|
| Model Identifier | iML1515 |
| Total Metabolites | 1,877 |
| Total Reactions | 2,712 |
| Total Genes | 1,516 |
| Total Compartments | 3 |
| Metabolic Coverage | 1.79 |

#### Metabolite Information

| | |
|---|---|
| Unique Metabolites | 1,169 |
| Duplicate Metabolites in Identical Compartments | 0 |
| Metabolites without Charge | 0 |
| Metabolites without Formula | 0 |
| Medium Components | 24 |

| | |
|---|---|
| bigg.metabolite | 100.0% |
| biocyc | 79.9% |
| **Metabolite Annotation Conformity Per Database** | Info |
| pubchem.compound | 0.0% |
| kegg.compound | 100.0% |
| seed.compound | 100.0% |
| inchikey | 0.0% |
| inchi | 0.0% |
| chebi | 100.0% |
| hmdb | 100.0% |
| reactome | 0.0% |
| metanetx.chemical | 100.0% |
| bigg.metabolite | 100.0% |
| biocyc | 100.0% |
| **Uniform Metabolite Identifier Namespace** | 100.0% |

**Sub Total** 79%

#### Annotation - Reactions

| | |
|---|---|
| **Presence of Reaction Annotation** | 100.0% |
| **Reaction Annotations Per Database** | Info |
| rhea | 43.4% |

| | |
|---|---|
| **Purely Metabolic Reactions with Constraints** | 2 |
| **Transport Reactions** | 831 |
| **Transport Reactions with Constraints** | 0 |
| **Thermodynamic Reversibility of Purely Metabolic Reactions** | 0.41 |
| **Reactions With Partially Identical Annotations** | 0.28 |
| **Duplicate Reactions** | 0.00 |
| **Reactions With Identical Genes** | 0.54 |

#### Gene-Protein-Reaction (GPR) Associations

| | |
|---|---|
| **Reactions without GPR** | 109 |
| **Fraction of Transport Reactions without GPR** | 0.06 |
| **Enzyme Complexes** | 309 |

#### Biomass

| | |
|---|---|
| **Biomass Reactions Identified** | 2 |
| **Biomass Consistency** | Info |
| BIOMASS_Ec_iML1515_WT_75p37M | 1.00 |
| BIOMASS_Ec_iML1515_core_75p37M | 1.00 |
| **Biomass Production In Default Medium** | Info |

| | | |
|---|---|---|
| metanetx.reaction | 94.4% | |
| bigg.reaction | 100.0% | |
| reactome | 0.0% | |
| ec-code | 40.4% | |
| brenda | 0.0% | |
| biocyc | 40.2% | |
| **Reaction Annotation Conformity Per Database** | | Info |
| rhea | 98.1% | |
| kegg.reaction | 100.0% | |
| seed.reaction | 100.0% | |
| metanetx.reaction | 100.0% | |
| bigg.reaction | 100.0% | |
| reactome | 0.0% | |
| ec-code | 98.0% | |
| brenda | 0.0% | |
| biocyc | 100.0% | |
| **Uniform Reaction Identifier Namespace** | 100.0% | |

**Sub Total** 81%

#### Annotation - Genes

| | |
|---|---|
| **Presence of Gene Annotation** | 100.0% |

| | | |
|---|---|---|
| **Unrealistic Growth Rate In Default Medium** | | Info |
| BIOMASS_Ec_iML1515_WT_75p37M | false | |
| BIOMASS_Ec_iML1515_core_75p37M | false | |
| **Biomass Production In Complete Medium** | | Info |
| BIOMASS_Ec_iML1515_WT_75p37M | 74.96 | |
| BIOMASS_Ec_iML1515_core_75p37M | 74.90 | |
| **Blocked Biomass Precursors In Default Medium** | | Info |
| BIOMASS_Ec_iML1515_WT_75p37M | 1 | |
| BIOMASS_Ec_iML1515_core_75p37M | 0 | |
| **Blocked Biomass Precursors In Complete Medium** | | Info |
| BIOMASS_Ec_iML1515_WT_75p37M | 0 | |
| BIOMASS_Ec_iML1515_core_75p37M | 0 | |
| **Ratio of Direct Metabolites in Biomass Reaction** | | Info |
| BIOMASS_Ec_iML1515_WT_75p37M | 0.09 | |
| BIOMASS_Ec_iML1515_core_75p37M | 0.13 | |
| **Number of Missing Essential Biomass Precursors** | | Info |
| BIOMASS_Ec_iML1515_WT_75p37M | 1 | |
| BIOMASS_Ec_iML1515_core_75p37M | 1 | |

#### Energy Metabolism

| | |
|---|---|
| **Non-Growth Associated Maintenance Reaction** | 1 |

| | |
|---|---|
| uniprot | 99.9% |
| ecogene | 99.9% |
| kegg.genes | 0.0% |
| ncbigi | 99.8% |
| ncbigene | 99.9% |
| ncbiprotein | 0.0% |
| ccds | 0.0% |
| hprd | 0.0% |
| asap | 99.9% |

**Gene Annotation Conformity Per Database** Info

| | |
|---|---|
| refseq | 0.0% |
| uniprot | 100.0% |
| ecogene | 100.0% |
| kegg.genes | 0.0% |
| ncbigi | 0.2% |
| ncbigene | 100.0% |
| ncbiprotein | 0.0% |
| ccds | 0.0% |
| hprd | 0.0% |
| asap | 100.0% |

**Sub Total** 63%

| | |
|---|---|
| BIOMASS_Ec_iML1515_core_75p37M | true |

**Number of Reversible Oxygen-Containing Reactions** 5

**Erroneous Energy-generating Cycles** Info

| | |
|---|---|
| MNXM3 | Skipped |
| MNXM63 | Skipped |
| MNXM51 | Skipped |
| MNXM121 | Skipped |
| MNXM423 | Skipped |
| MNXM6 | Skipped |
| MNXM10 | Skipped |
| MNXM38 | Skipped |
| MNXM208 | Skipped |
| MNXM191 | Skipped |
| MNXM223 | Skipped |
| MNXM7517 | Skipped |
| MNXM12233 | Skipped |
| MNXM558 | Skipped |
| MNXM21 | Skipped |
| MNXM89557 | Skipped |

### Network Topology

**Universally Blocked Reactions** 260

| | | |
|---|---|---|
| Metabolite SBO:0000247 Presence | 100.0% | |
| Reaction General SBO Presence | 100.0% | |
| Metabolic Reaction SBO:0000176 Presence | 100.0% | |
| Transport Reaction SBO:0000185 Presence | 98.8% | |
| Exchange Reaction SBO:0000627 Presence | 100.0% | |
| Demand Reaction SBO:0000628 Presence | 100.0% | |
| Sink Reactions SBO:0000632 Presence | Skipped | |
| Gene General SBO Presence | 100.0% | |
| Gene SBO:0000243 Presence | 100.0% | |
| Biomass Reactions SBO:0000629 Presence | 100.0% | |
| Sub Total | 91% | X2 |
| Total Score | 91% | |

Total Score

# 91%

Score per Category

| | |
|---|---|
| Stoichiometrically Balanced Cycles | 61 |
| Metabolite Production In Complete Medium | 181 |
| Metabolite Consumption In Complete Medium | 233 |

#### Matrix Conditioning

| | |
|---|---|
| Ratio Min/Max Non-Zero Coefficients | 0.00 |
| Independent Conservation Relations | 31 |
| Rank | 1845 |
| Degrees Of Freedom | 867 |

#### Experimental Data Comparison

| | |
|---|---|
| Growth Prediction | Skipped |
| Gene Essentiality Prediction | Skipped |

#### Misc. Tests

#### Environment

| | |
|---|---|
| Python Version | 3.6.10 |
| Platform | Linux |
| Memote Version | 0.10.2 |

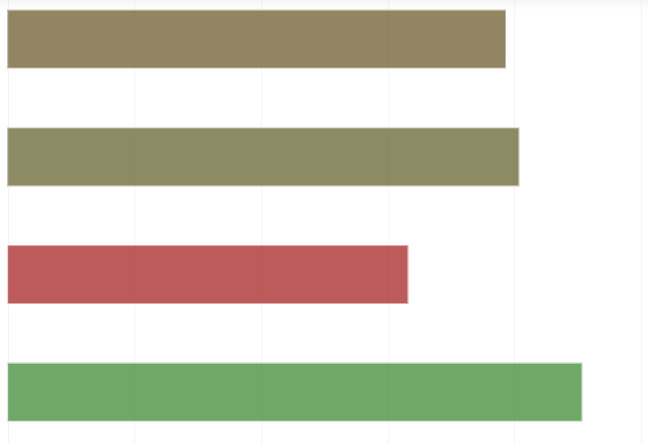
