## Supplemental Data 3 for "An updated genome-scale metabolic network reconstruction of *Pseudomonas aeruginosa* PA14 to characterize mucin-driven shifts in bacterial metabolism"

| ABTGC in silico formulation | | |
|---|---|---|
| Metabolite Name | Met ID | Lower Bound |
| H2O | cpd00001_e | -1000 |
| H+ | cpd00067_e | -1000 |
| Phosphate | cpd00009_e | -1000 |
| CO2 | cpd00011_e | -1000 |
| Fe2+ | cpd00021_e | -1000 |
| Mg | cpd00254_e | -1000 |
| Ca2+ | cpd00063_e | -1000 |
| Cl- | cpd00099_e | -1000 |
| NH3 | cpd00013_e | -1000 |
| Na+ | cpd00971_e | -1000 |
| K+ | cpd00205_e | -1000 |
| Sulfate | cpd00048_e | -1000 |
| D-glucose | cpd00027_e | -10 |
| L-Alanine | cpd00035_e | -10 |
| L-Arginine | cpd00051_e | -10 |
| L-Asparagine | cpd00132_e | -10 |
| L-Aspartate | cpd00041_e | -10 |
| L-Cysteine | cpd00084_e | -10 |
| L-Glutamine | cpd00053_e | -10 |
| L-Glutamate | cpd00023_e | -10 |
| Glycine | cpd00033_e | -10 |
| L-Histidine | cpd00119_e | -10 |
| L-Isoleucine | cpd00322_e | -10 |
| L-Leucine | cpd00107_e | -10 |
| L-Lysine | cpd00039_e | -10 |
| L-Phenylalanine | cpd00066_e | -10 |
| L-Proline | cpd00129_e | -10 |
| L-Serine | cpd00054_e | -10 |
| L-Threonine | cpd00161_e | -10 |
| L-Tyrosine | cpd00069_e | -10 |
| L-Valine | cpd00156_e | -10 |
| Thiamine | cpd00305_e | -10 |

| LB in silico formulation | | |
|---|---|---|
| Metabolite Name | Met ID | Lower Bound |
| H2O | cpd00001_e | -1000 |
| Phosphate | cpd00009_e | -1000 |
| CO2 | cpd00011_e | -1000 |
| Fe2+ | cpd00021_e | -1000 |
| Zn2+ | cpd00034_e | -1000 |
| Sulfate | cpd00048_e | -1000 |
| Cu2+ | cpd00058_e | -1000 |
| K+ | cpd00205_e | -1000 |
| Mg | cpd00254_e | -1000 |
| Na+ | cpd00971_e | -1000 |
| Cd2+ | cpd01012_e | -1000 |
| H+ | cpd00067_e' | -1000 |
| L-Glutamate | cpd00023_e | -10 |
| D-Glucose | cpd00027_e | -10 |
| Glycine | cpd00033_e | -10 |
| L-Alanine | cpd00035_e | -10 |
| L-Lysine | cpd00039_e | -10 |
| L-Aspartate | cpd00041_e | -10 |
| L-Arginine | cpd00051_e | -10 |
| L-Serine | cpd00054_e | -10 |
| L-Methionine | cpd00060_e | -10 |
| L-Tryptophan | cpd00065_e | -10 |
| L-Phenylalanine | cpd00066_e | -10 |
| L-Tyrosine | cpd00069_e | -10 |
| L-Cysteine | cpd00084_e | -10 |
| L-Leucine | cpd00107_e | -10 |
| L-Histidine | cpd00119_e | -10 |
| L-Proline | cpd00129_e | -10 |
| L-Valine | cpd00156_e | -10 |
| L-Threonine | cpd00161_e | -10 |
| Thiamin | cpd00305_e | -10 |
| L-Isoleucine | cpd00322_e | -10 |
| Uracil | cpd00092_e | -10 |
| Cytosine | cpd00307_e | -10 |
| L-5'-Deoxyadenosine | cpd03091_e | -10 |

| SCFM in silico formulation | | |
|---|---|---|
| Metabolite Name | Met ID | Lower Bound |
| H2O | cpd00001_e | -10 |
| Phosphate | cpd00009_e | -10 |
| CO2 | cpd00011_e | -10 |
| Fe2+ | cpd00021_e | -10 |
| L-Glutamate | cpd00023_e | -10 |
| D-Glucose | cpd00027_e | -10 |
| Glycine | cpd00033_e | -10 |
| L-Alanine | cpd00035_e | -10 |
| L-Lysine | cpd00039_e | -10 |
| L-Aspartate | cpd00041_e | -10 |
| Sulfate | cpd00048_e | -10 |
| L-Arginine | cpd00051_e | -10 |
| L-Serine | cpd00054_e | -10 |
| L-Methionine | cpd00060_e | -10 |
| Ornithine | cpd00064_e | -10 |
| L-Tryptophan | cpd00065_e | -10 |
| L-Phenylalanine | cpd00066_e | -10 |
| H+ | cpd00067_e | -10 |
| L-Tyrosine | cpd00069_e | -10 |
| L-Cysteine | cpd00084_e | -10 |
| L-Leucine | cpd00107_e | -10 |
| L-Histidine | cpd00119_e | -10 |
| L-Proline | cpd00129_e | -10 |
| L-Valine | cpd00156_e | -10 |
| D-Lactate | cpd00221_e | -10 |
| L-Threonine | cpd00161_e | -10 |
| K+ | cpd00205_e | -10 |
| Nitrate | cpd00209_e | -10 |
| Mg | cpd00254_e | -10 |
| L-Isoleucine | cpd00322_e | -10 |
| Na+ | cpd00971_e | -10 |
| NH3 | cpd00013_e | -10 |

| Glucose Minimal in silico formulation | | |
|---|---|---|
| Metabolite | Met ID | Lower Bound |
| H2O | cpd00001_e | -1000 |
| Phosphate | cpd00009_e | -1000 |
| CO2 | cpd00011_e | -1000 |
| Fe2+ | cpd00021_e | -1000 |
| Mn2+ | cpd00030_e | -1000 |
| Zn2+ | cpd00034_e | -1000 |
| Sulfate | cpd00048_e | -1000 |
| Cu2+ | cpd00058_e | -1000 |
| H+ | cpd00067_e | -1000 |
| Co2+ | cpd00149_e | -1000 |
| K+ | cpd00205_e | -1000 |
| Mg | cpd00254_e | -1000 |
| nitrogen | cpd00528_e | -1000 |
| Na+ | cpd00971_e | -1000 |
| NH3 | cpd00013_e | -1000 |
| Cd2+ | cpd01012_e | -1000 |
| fe3 | cpd10516_e | -1000 |
| Ni2+ | cpd00244_e' | -1000 |
| Glucose | cpd00027_e | -10 |
