## Supplemental Data 4 for "An updated genome-scale metabolic network reconstruction of *Pseudomonas aeruginosa* PA14 to characterize mucin-driven shifts in bacterial metabolism"

| Metabolite ID | Carbon Source Name | In Vitro | iPau1129 | iPau21 |
|---|---|---|---|---|
| cpd00489 | 4-Hydroxyphenylacetate | Growth | No growth | Growth |
| cpd00266 | Carnitine | Growth | No growth | Growth |
| cpd00222 | GLCN | Growth | No growth | Growth |
| cpd11592 | gly-glu-L | Growth | No growth | Growth |
| cpd11588 | gly-pro-L | Growth | No growth | Growth |
| cpd00477 | N-Acetyl-L-glutamate | Growth | No growth | Growth |
| cpd00141 | Propionate | Growth | No growth | Growth |
| cpd00851 | trans-4-Hydroxy-L-proline | Growth | No growth | Growth |
| cpd00024 | 2-Oxoglutarate | Growth | Growth | Growth |
| cpd00136 | 4-Hydroxybenzoate | Growth | Growth | Growth |
| cpd00029 | Acetate | Growth | Growth | Growth |
| cpd00182 | Adenosine | Growth | Growth | Growth |
| cpd00162 | Aminoethanol | Growth | Growth | Growth |
| cpd00137 | Citrate | Growth | Growth | Growth |
| cpd00117 | D-Alanine | Growth | Growth | Growth |
| cpd00082 | D-Fructose | Growth | Growth | Growth |
| cpd00027 | D-Glucose | Growth | Growth | Growth |
| cpd00314 | D-Mannitol | Growth | Growth | Growth |
| cpd00106 | Fumarate | Growth | Growth | Growth |
| cpd00281 | GABA | Growth | Growth | Growth |
| cpd00100 | Glycerol | Growth | Growth | Growth |
| cpd00080 | Glycerol-3-phosphate | Growth | Growth | Growth |
| cpd00033 | Glycine | Growth | Growth | Growth |
| cpd00380 | Itaconate | Growth | Growth | Growth |
| cpd00035 | L-Alanine | Growth | Growth | Growth |
| cpd00051 | L-Arginine | Growth | Growth | Growth |
| cpd00132 | L-Asparagine | Growth | Growth | Growth |
| cpd00041 | L-Aspartate | Growth | Growth | Growth |
| cpd00023 | L-Glutamate | Growth | Growth | Growth |
| cpd00053 | L-Glutamine | Growth | Growth | Growth |
| cpd00119 | L-Histidine | Growth | Growth | Growth |
| cpd00322 | L-Isoleucine | Growth | Growth | Growth |
| cpd00159 | L-Lactate | Growth | Growth | Growth |
| cpd00107 | L-Leucine | Growth | Growth | Growth |
| cpd00130 | L-Malate | Growth | Growth | Growth |
| cpd00066 | L-Phenylalanine | Growth | Growth | Growth |
| cpd00129 | L-Proline | Growth | Growth | Growth |
| cpd00054 | L-Serine | Growth | Growth | Growth |
| cpd00308 | Malonate | Growth | Growth | Growth |
| cpd00064 | Ornithine | Growth | Growth | Growth |
| cpd00118 | Putrescine | Growth | Growth | Growth |
| cpd00020 | Pyruvate | Growth | Growth | Growth |
| cpd00036 | Succinate | Growth | Growth | Growth |
| cpd00386 | D-Malate | No growth | Growth | No growth |
| cpd01949 | (SS)-23-Butanediol | No growth | No growth | No growth |
| cpd00094 | 2-Oxobutyrate | No growth | No growth | No growth |
| cpd03320 | 3-Hydroxyphenylacetate | No growth | No growth | No growth |
| cpd00142 | Acetoacetate | No growth | No growth | No growth |
| cpd00361 | ACTN | No growth | No growth | No growth |

| | | | | |
|---|---|---|---|---|
| cpd00361 | ACTN | No growth | No growth | No growth |
| cpd01262 | Amylotriose | No growth | No growth | No growth |
| cpd00158 | CELB | No growth | No growth | No growth |
| cpd00072 | D-fructose-6-phosphate | No growth | No growth | No growth |
| cpd00280 | D-Galacturonate | No growth | No growth | No growth |
| cpd00609 | D-Glucarate | No growth | No growth | No growth |
| cpd00079 | D-glucose-6-phosphate | No growth | No growth | No growth |
| cpd00138 | D-Mannose | No growth | No growth | No growth |
| cpd00652 | D-Mucic - acid | No growth | No growth | No growth |
| cpd00294 | dAMP | No growth | No growth | No growth |
| cpd00047 | Formate | No growth | No growth | No growth |
| cpd00089 | Glucose-1-phosphate | No growth | No growth | No growth |
| cpd00164 | Glucuronate | No growth | No growth | No growth |
| cpd11589 | gly-asp-L | No growth | No growth | No growth |
| cpd00155 | Glycogen | No growth | No growth | No growth |
| cpd00139 | Glycolate | No growth | No growth | No growth |
| cpd00040 | Glyoxalate | No growth | No growth | No growth |
| cpd00224 | L-Arabinose | No growth | No growth | No growth |
| cpd00227 | L-Homoserine | No growth | No growth | No growth |
| cpd00121 | L-Inositol | No growth | No growth | No growth |
| cpd00060 | L-Methionine | No growth | No growth | No growth |
| cpd00156 | L-Valine | No growth | No growth | No growth |
| cpd00179 | Maltose | No growth | No growth | No growth |
| cpd00599 | salicylate | No growth | No growth | No growth |
| cpd00588 | Sorbitol | No growth | No growth | No growth |
| cpd00666 | Tartrate | No growth | No growth | No growth |
| cpd00666 | Tartrate | No growth | No growth | No growth |
| cpd00666 | Tartrate | No growth | No growth | No growth |
| cpd00184 | Thymidine | No growth | No growth | No growth |
| cpd01242 | Thyminose | No growth | No growth | No growth |
| cpd00794 | TRHL | No growth | No growth | No growth |
| cpd00154 | Xylose | No growth | No growth | No growth |
| cpd00477 | N-Acetyl-L-glutamate | Growth | No growth | Growth |
| cpd11592 | gly-glu-L | Growth | No growth | Growth |
| cpd11588 | gly-pro-L | Growth | No growth | Growth |
| cpd00851 | trans-4-Hydroxy-L-proline | Growth | No growth | Growth |
| cpd00266 | Carnitine | Growth | No growth | Growth |
| cpd00222 | GLCN | Growth | No growth | Growth |
| cpd00489 | 4-Hydroxyphenylacetate | Growth | No growth | Growth |
| cpd00141 | Propionate | Growth | No growth | Growth |
| cpd01502 | Citraconate | No growth | No growth | Growth |
| cpd00105 | D-Ribose | No growth | Growth | Growth |
| cpd00550 | D-Serine | No growth | Growth | Growth |
| cpd00246 | Inosine | No growth | Growth | Growth |
| cpd00161 | L-Threonine | No growth | Growth | Growth |
| cpd00797 | (R)-3-Hydroxybutanoate | Growth | No growth | No growth |
| cpd00211 | Butyrate | Growth | No growth | No growth |
| cpd11585 | L-alanylglycine | Growth | No growth | No growth |
| cpd00039 | L-Lysine | Growth | No growth | No growth |
| cpd00249 | Uridine | Growth | No growth | No growth |
