## Supplemental Data 5 for "An updated genome-scale metabolic network reconstruction of *Pseudomonas aeruginosa* PA14 to characterize mucin-driven shifts in bacterial metabolism"

| Gene Name | In Vitro | iPau21 |
|---|---|---|
| PA14_23860 | Essential | Essential |
| PA14_25660 | Essential | Essential |
| PA14_17290 | Essential | Essential |
| PA14_69440 | Essential | Essential |
| PA14_15310 | Essential | Essential |
| PA14_17220 | Essential | Essential |
| PA14_61710 | Essential | Essential |
| PA14_62850 | Essential | Essential |
| PA14_64100 | Essential | Essential |
| PA14_57410 | Essential | Essential |
| PA14_66550 | Essential | Essential |
| PA14_41350 | Essential | Essential |
| PA14_57500 | Essential | Essential |
| PA14_25550 | Essential | Essential |
| PA14_73170 | Essential | Essential |
| PA14_17270 | Essential | Essential |
| PA14_17120 | Essential | Essential |
| PA14_41400 | Essential | Essential |
| PA14_04580 | Essential | Essential |
| PA14_61580 | Essential | Essential |
| PA14_30670 | Essential | Essential |
| PA14_43680 | Essential | Essential |
| PA14_17080 | Essential | Essential |
| PA14_66900 | Essential | Essential |
| PA14_04310 | Essential | Essential |
| PA14_60330 | Essential | Essential |
| PA14_17190 | Essential | Essential |
| PA14_65500 | Essential | Essential |
| PA14_64110 | Essential | Essential |
| PA14_57390 | Essential | Essential |
| PA14_66610 | Essential | Essential |
| PA14_14880 | Essential | Essential |
| PA14_23560 | Essential | Essential |
| PA14_23800 | Essential | Essential |
| PA14_57340 | Essential | Essential |
| PA14_30110 | Essential | Essential |
| PA14_65960 | Essential | Essential |
| PA14_62940 | Essential | Essential |
| PA14_23320 | Essential | Essential |
| PA14_17210 | Essential | Essential |
| PA14_61660 | Essential | Essential |
| PA14_25530 | Essential | Essential |
| PA14_57330 | Essential | Essential |
| PA14_07910 | Essential | Essential |
| PA14_07590 | Essential | Essential |
| PA14_66060 | Essential | Essential |
| PA14_25510 | Essential | Essential |
| PA14_04480 | Essential | Essential |
| PA14_66230 | Essential | Essential |
| PA14_16950 | Essential | Essential |
| PA14_51270 | Essential | Essential |
| PA14_08400 | Essential | Essential |
| PA14_61750 | Essential | Essential |
| PA14_23220 | Essential | Essential |
| PA14_14820 | Essential | Essential |
| PA14_69450 | Essential | Essential |
| PA14_68200 | Essential | Essential |
| PA14_11550 | Essential | Essential |
| PA14_17130 | Essential | Essential |
| PA14_57380 | Essential | Essential |
| PA14_44710 | Nonessential | Nonessential |
| PA14_09710 | Nonessential | Nonessential |
| PA14_33860 | Nonessential | Nonessential |
| PA14_71030 | Nonessential | Nonessential |
| PA14_71500 | Nonessential | Nonessential |
| PA14_60730 | Nonessential | Nonessential |
| PA14_58220 | Nonessential | Nonessential |
| PA14_60270 | Nonessential | Nonessential |
| PA14_38460 | Nonessential | Nonessential |
| PA14_05080 | Nonessential | Nonessential |
| PA14_71410 | Nonessential | Nonessential |
| PA14_10900 | Nonessential | Nonessential |
| PA14_52150 | Nonessential | Nonessential |

| | | |
|---|---|---|
| PA14_02330 | Nonessential | Nonessential |
| PA14_05200 | Nonessential | Nonessential |
| PA14_32110 | Nonessential | Nonessential |
| PA14_06830 | Nonessential | Nonessential |
| PA14_11760 | Nonessential | Nonessential |
| PA14_71740 | Nonessential | Nonessential |
| PA14_70920 | Nonessential | Nonessential |
| PA14_66820 | Nonessential | Nonessential |
| PA14_54150 | Nonessential | Nonessential |
| PA14_07900 | Nonessential | Nonessential |
| PA14_52660 | Nonessential | Nonessential |
| PA14_67770 | Nonessential | Nonessential |
| PA14_23370 | Nonessential | Nonessential |
| PA14_13800 | Nonessential | Nonessential |
| PA14_24780 | Nonessential | Nonessential |
| PA14_68890 | Nonessential | Nonessential |
| PA14_54940 | Nonessential | Nonessential |
| PA14_04110 | Nonessential | Nonessential |
| PA14_06600 | Nonessential | Nonessential |
| PA14_43970 | Nonessential | Nonessential |
| PA14_20650 | Nonessential | Nonessential |
| PA14_03950 | Nonessential | Nonessential |
| PA14_05690 | Nonessential | Nonessential |
| PA14_05310 | Nonessential | Nonessential |
| PA14_34790 | Nonessential | Nonessential |
| PA14_36120 | Nonessential | Nonessential |
| PA14_51990 | Nonessential | Nonessential |
| PA14_72780 | Nonessential | Nonessential |
| PA14_68370 | Nonessential | Nonessential |
| PA14_09600 | Nonessential | Nonessential |
| PA14_67310 | Nonessential | Nonessential |
| PA14_52840 | Nonessential | Nonessential |
| PA14_44210 | Nonessential | Nonessential |
| PA14_71260 | Nonessential | Nonessential |
| PA14_10650 | Nonessential | Nonessential |
| PA14_68920 | Nonessential | Nonessential |
| PA14_46240 | Nonessential | Nonessential |
| PA14_38470 | Nonessential | Nonessential |
| PA14_69850 | Nonessential | Nonessential |
| PA14_02990 | Nonessential | Nonessential |
| PA14_14060 | Nonessential | Nonessential |
| PA14_26500 | Nonessential | Nonessential |
| PA14_22050 | Nonessential | Nonessential |
| PA14_29890 | Nonessential | Nonessential |
| PA14_24960 | Nonessential | Nonessential |
| PA14_61680 | Nonessential | Nonessential |
| PA14_35460 | Nonessential | Nonessential |
| PA14_16070 | Nonessential | Nonessential |
| PA14_41640 | Nonessential | Nonessential |
| PA14_23450 | Nonessential | Nonessential |
| PA14_09450 | Nonessential | Nonessential |
| PA14_43460 | Nonessential | Nonessential |
| PA14_55740 | Nonessential | Nonessential |
| PA14_34280 | Nonessential | Nonessential |
| PA14_34340 | Nonessential | Nonessential |
| PA14_03880 | Nonessential | Nonessential |
| PA14_71140 | Nonessential | Nonessential |
| PA14_68870 | Nonessential | Nonessential |
| PA14_72510 | Nonessential | Nonessential |
| PA14_32270 | Nonessential | Nonessential |
| PA14_47840 | Nonessential | Nonessential |
| PA14_43110 | Nonessential | Nonessential |
| PA14_36840 | Nonessential | Nonessential |
| PA14_24290 | Nonessential | Nonessential |
| PA14_49840 | Nonessential | Nonessential |
| PA14_52870 | Nonessential | Nonessential |
| PA14_62570 | Nonessential | Nonessential |
| PA14_18120 | Nonessential | Nonessential |
| PA14_41130 | Nonessential | Nonessential |
| PA14_47280 | Nonessential | Nonessential |
| PA14_18740 | Nonessential | Nonessential |
| PA14_09290 | Nonessential | Nonessential |
| PA14_32230 | Nonessential | Nonessential |
| PA14_51050 | Nonessential | Nonessential |

| | | |
|---|---|---|
| PA14_23080 | Nonessential | Nonessential |
| PA14_23210 | Nonessential | Nonessential |
| PA14_68070 | Nonessential | Nonessential |
| PA14_31970 | Nonessential | Nonessential |
| PA14_48000 | Nonessential | Nonessential |
| PA14_35270 | Nonessential | Nonessential |
| PA14_47860 | Nonessential | Nonessential |
| PA14_66380 | Nonessential | Nonessential |
| PA14_69170 | Nonessential | Nonessential |
| PA14_10590 | Nonessential | Nonessential |
| PA14_23390 | Nonessential | Nonessential |
| PA14_54040 | Nonessential | Nonessential |
| PA14_06720 | Nonessential | Nonessential |
| PA14_44740 | Nonessential | Nonessential |
| PA14_02550 | Nonessential | Nonessential |
| PA14_63990 | Nonessential | Nonessential |
| PA14_37965 | Nonessential | Nonessential |
| PA14_22690 | Nonessential | Nonessential |
| PA14_46970 | Nonessential | Nonessential |
| PA14_22980 | Nonessential | Nonessential |
| PA14_39280 | Nonessential | Nonessential |
| PA14_73240 | Nonessential | Nonessential |
| PA14_26400 | Nonessential | Nonessential |
| PA14_51420 | Nonessential | Nonessential |
| PA14_52210 | Nonessential | Nonessential |
| PA14_09220 | Nonessential | Nonessential |
| PA14_23440 | Nonessential | Nonessential |
| PA14_62910 | Nonessential | Nonessential |
| PA14_03810 | Nonessential | Nonessential |
| PA14_71000 | Nonessential | Nonessential |
| PA14_36050 | Nonessential | Nonessential |
| PA14_38510 | Nonessential | Nonessential |
| PA14_23250 | Nonessential | Nonessential |
| PA14_23930 | Nonessential | Nonessential |
| PA14_36220 | Nonessential | Nonessential |
| PA14_69140 | Nonessential | Nonessential |
| PA14_23380 | Nonessential | Nonessential |
| PA14_34390 | Nonessential | Nonessential |
| PA14_66840 | Nonessential | Nonessential |
| PA14_47850 | Nonessential | Nonessential |
| PA14_36270 | Nonessential | Nonessential |
| PA14_66680 | Nonessential | Nonessential |
| PA14_02900 | Nonessential | Nonessential |
| PA14_18480 | Nonessential | Nonessential |
| PA14_19900 | Nonessential | Nonessential |
| PA14_53290 | Nonessential | Nonessential |
| PA14_47720 | Nonessential | Nonessential |
| PA14_46070 | Nonessential | Nonessential |
| PA14_18610 | Nonessential | Nonessential |
| PA14_10260 | Nonessential | Nonessential |
| PA14_32650 | Nonessential | Nonessential |
| PA14_65370 | Nonessential | Nonessential |
| PA14_03000 | Nonessential | Nonessential |
| PA14_62620 | Nonessential | Nonessential |
| PA14_02850 | Nonessential | Nonessential |
| PA14_07930 | Nonessential | Nonessential |
| PA14_68900 | Nonessential | Nonessential |
| PA14_19350 | Nonessential | Nonessential |
| PA14_38860 | Nonessential | Nonessential |
| PA14_40890 | Nonessential | Nonessential |
| PA14_47670 | Nonessential | Nonessential |
| PA14_39590 | Nonessential | Nonessential |
| PA14_06500 | Nonessential | Nonessential |
| PA14_67260 | Nonessential | Nonessential |
| PA14_10550 | Nonessential | Nonessential |
| PA14_35890 | Nonessential | Nonessential |
| PA14_48440 | Nonessential | Nonessential |
| PA14_36230 | Nonessential | Nonessential |
| PA14_56340 | Nonessential | Nonessential |
| PA14_52900 | Nonessential | Nonessential |
| PA14_53470 | Nonessential | Nonessential |
| PA14_62130 | Nonessential | Nonessential |
| PA14_20560 | Nonessential | Nonessential |
| PA14_64900 | Nonessential | Nonessential |

| | | |
|---|---|---|
| PA14_71460 | Nonessential | Nonessential |
| PA14_03030 | Nonessential | Nonessential |
| PA14_40470 | Nonessential | Nonessential |
| PA14_60120 | Nonessential | Nonessential |
| PA14_26220 | Nonessential | Nonessential |
| PA14_07700 | Nonessential | Nonessential |
| PA14_71990 | Nonessential | Nonessential |
| PA14_54910 | Nonessential | Nonessential |
| PA14_08360 | Nonessential | Nonessential |
| PA14_03830 | Nonessential | Nonessential |
| PA14_14100 | Nonessential | Nonessential |
| PA14_35490 | Nonessential | Nonessential |
| PA14_64290 | Nonessential | Nonessential |
| PA14_52690 | Nonessential | Nonessential |
| PA14_12410 | Nonessential | Nonessential |
| PA14_02640 | Nonessential | Nonessential |
| PA14_64390 | Nonessential | Nonessential |
| PA14_70140 | Nonessential | Nonessential |
| PA14_00440 | Nonessential | Nonessential |
| PA14_51040 | Nonessential | Nonessential |
| PA14_19100 | Nonessential | Nonessential |
| PA14_36810 | Nonessential | Nonessential |
| PA14_18520 | Nonessential | Nonessential |
| PA14_49130 | Nonessential | Nonessential |
| PA14_71530 | Nonessential | Nonessential |
| PA14_41510 | Nonessential | Nonessential |
| PA14_17860 | Nonessential | Nonessential |
| PA14_73310 | Nonessential | Nonessential |
| PA14_52850 | Nonessential | Nonessential |
| PA14_07870 | Nonessential | Nonessential |
| PA14_39350 | Nonessential | Nonessential |
| PA14_72640 | Nonessential | Nonessential |
| PA14_02450 | Nonessential | Nonessential |
| PA14_71220 | Nonessential | Nonessential |
| PA14_04240 | Nonessential | Nonessential |
| PA14_64870 | Nonessential | Nonessential |
| PA14_18510 | Nonessential | Nonessential |
| PA14_44800 | Nonessential | Nonessential |
| PA14_67880 | Nonessential | Nonessential |
| PA14_32240 | Nonessential | Nonessential |
| PA14_54520 | Nonessential | Nonessential |
| PA14_68090 | Nonessential | Nonessential |
| PA14_10250 | Nonessential | Nonessential |
| PA14_56680 | Nonessential | Nonessential |
| PA14_61460 | Nonessential | Nonessential |
| PA14_04230 | Nonessential | Nonessential |
| PA14_69430 | Nonessential | Nonessential |
| PA14_71910 | Nonessential | Nonessential |
| PA14_72690 | Nonessential | Nonessential |
| PA14_44970 | Nonessential | Nonessential |
| PA14_13590 | Nonessential | Nonessential |
| PA14_29410 | Nonessential | Nonessential |
| PA14_72630 | Nonessential | Nonessential |
| PA14_72850 | Nonessential | Nonessential |
| PA14_50530 | Nonessential | Nonessential |
| PA14_42230 | Nonessential | Nonessential |
| PA14_13430 | Nonessential | Nonessential |
| PA14_10610 | Nonessential | Nonessential |
| PA14_21160 | Nonessential | Nonessential |
| PA14_01660 | Nonessential | Nonessential |
| PA14_66310 | Nonessential | Nonessential |
| PA14_34410 | Nonessential | Nonessential |
| PA14_01580 | Nonessential | Nonessential |
| PA14_71970 | Nonessential | Nonessential |
| PA14_15820 | Nonessential | Nonessential |
| PA14_16360 | Nonessential | Nonessential |
| PA14_11190 | Nonessential | Nonessential |
| PA14_10630 | Nonessential | Nonessential |
| PA14_40670 | Nonessential | Nonessential |
| PA14_66330 | Nonessential | Nonessential |
| PA14_66040 | Nonessential | Nonessential |
| PA14_57800 | Nonessential | Nonessential |
| PA14_46470 | Nonessential | Nonessential |
| PA14_38850 | Nonessential | Nonessential |

| | | |
|---|---|---|
| PA14_43400 | Nonessential | Nonessential |
| PA14_48570 | Nonessential | Nonessential |
| PA14_29970 | Nonessential | Nonessential |
| PA14_05840 | Nonessential | Nonessential |
| PA14_58420 | Nonessential | Nonessential |
| PA14_18275 | Nonessential | Nonessential |
| PA14_05270 | Nonessential | Nonessential |
| PA14_67920 | Nonessential | Nonessential |
| PA14_52700 | Nonessential | Nonessential |
| PA14_01250 | Nonessential | Nonessential |
| PA14_17610 | Nonessential | Nonessential |
| PA14_69670 | Nonessential | Nonessential |
| PA14_70950 | Nonessential | Nonessential |
| PA14_52050 | Nonessential | Nonessential |
| PA14_23410 | Nonessential | Nonessential |
| PA14_10640 | Nonessential | Nonessential |
| PA14_47100 | Nonessential | Nonessential |
| PA14_46930 | Nonessential | Nonessential |
| PA14_13810 | Nonessential | Nonessential |
| PA14_36960 | Nonessential | Nonessential |
| PA14_39880 | Nonessential | Nonessential |
| PA14_57540 | Nonessential | Nonessential |
| PA14_35920 | Nonessential | Nonessential |
| PA14_41670 | Nonessential | Nonessential |
| PA14_50560 | Nonessential | Nonessential |
| PA14_66940 | Nonessential | Nonessential |
| PA14_33010 | Nonessential | Nonessential |
| PA14_34360 | Nonessential | Nonessential |
| PA14_57570 | Nonessential | Nonessential |
| PA14_66350 | Nonessential | Nonessential |
| PA14_67350 | Nonessential | Nonessential |
| PA14_67440 | Nonessential | Nonessential |
| PA14_57190 | Nonessential | Nonessential |
| PA14_60710 | Nonessential | Nonessential |
| PA14_02580 | Nonessential | Nonessential |
| PA14_02070 | Nonessential | Nonessential |
| PA14_29990 | Nonessential | Nonessential |
| PA14_17960 | Nonessential | Nonessential |
| PA14_67970 | Nonessential | Nonessential |
| PA14_67320 | Nonessential | Nonessential |
| PA14_37340 | Nonessential | Nonessential |
| PA14_11030 | Nonessential | Nonessential |
| PA14_09270 | Nonessential | Nonessential |
| PA14_19580 | Nonessential | Nonessential |
| PA14_52460 | Nonessential | Nonessential |
| PA14_43380 | Nonessential | Nonessential |
| PA14_48470 | Nonessential | Nonessential |
| PA14_56240 | Nonessential | Nonessential |
| PA14_11130 | Nonessential | Nonessential |
| PA14_45170 | Nonessential | Nonessential |
| PA14_18970 | Nonessential | Nonessential |
| PA14_36690 | Nonessential | Nonessential |
| PA14_41530 | Nonessential | Nonessential |
| PA14_17320 | Nonessential | Nonessential |
| PA14_04220 | Nonessential | Nonessential |
| PA14_14010 | Nonessential | Nonessential |
| PA14_34050 | Nonessential | Nonessential |
| PA14_35500 | Nonessential | Nonessential |
| PA14_22000 | Nonessential | Nonessential |
| PA14_29930 | Nonessential | Nonessential |
| PA14_49870 | Nonessential | Nonessential |
| PA14_51410 | Nonessential | Nonessential |
| PA14_08340 | Nonessential | Nonessential |
| PA14_43440 | Nonessential | Nonessential |
| PA14_02340 | Nonessential | Nonessential |
| PA14_45430 | Nonessential | Nonessential |
| PA14_17620 | Nonessential | Nonessential |
| PA14_61040 | Nonessential | Nonessential |
| PA14_11810 | Nonessential | Nonessential |
| PA14_73290 | Nonessential | Nonessential |
| PA14_38840 | Nonessential | Nonessential |
| PA14_45190 | Nonessential | Nonessential |
| PA14_18580 | Nonessential | Nonessential |
| PA14_05460 | Nonessential | Nonessential |

| | | |
|---|---|---|
| PA14_52990 | Nonessential | Nonessential |
| PA14_19130 | Nonessential | Nonessential |
| PA14_41010 | Nonessential | Nonessential |
| PA14_39640 | Nonessential | Nonessential |
| PA14_20140 | Nonessential | Nonessential |
| PA14_71800 | Nonessential | Nonessential |
| PA14_22990 | Nonessential | Nonessential |
| PA14_35940 | Nonessential | Nonessential |
| PA14_61480 | Nonessential | Nonessential |
| PA14_18140 | Nonessential | Nonessential |
| PA14_39970 | Nonessential | Nonessential |
| PA14_63580 | Nonessential | Nonessential |
| PA14_52750 | Nonessential | Nonessential |
| PA14_64890 | Nonessential | Nonessential |
| PA14_68780 | Nonessential | Nonessential |
| PA14_18430 | Nonessential | Nonessential |
| PA14_64960 | Nonessential | Nonessential |
| PA14_20330 | Nonessential | Nonessential |
| PA14_23460 | Nonessential | Nonessential |
| PA14_19050 | Nonessential | Nonessential |
| PA14_06750 | Nonessential | Nonessential |
| PA14_18300 | Nonessential | Nonessential |
| PA14_23170 | Nonessential | Nonessential |
| PA14_52790 | Nonessential | Nonessential |
| PA14_69925 | Nonessential | Nonessential |
| PA14_64860 | Nonessential | Nonessential |
| PA14_21370 | Nonessential | Nonessential |
| PA14_18565 | Nonessential | Nonessential |
| PA14_25580 | Nonessential | Nonessential |
| PA14_23840 | Nonessential | Nonessential |
| PA14_49760 | Nonessential | Nonessential |
| PA14_13780 | Nonessential | Nonessential |
| PA14_38690 | Nonessential | Nonessential |
| PA14_28650 | Nonessential | Nonessential |
| PA14_53800 | Nonessential | Nonessential |
| PA14_51390 | Nonessential | Nonessential |
| PA14_19370 | Nonessential | Nonessential |
| PA14_03550 | Nonessential | Nonessential |
| PA14_07940 | Nonessential | Nonessential |
| PA14_72590 | Nonessential | Nonessential |
| PA14_54620 | Nonessential | Nonessential |
| PA14_02570 | Nonessential | Nonessential |
| PA14_51150 | Nonessential | Nonessential |
| PA14_62600 | Nonessential | Nonessential |
| PA14_15700 | Nonessential | Nonessential |
| PA14_06660 | Nonessential | Nonessential |
| PA14_33630 | Nonessential | Nonessential |
| PA14_08480 | Nonessential | Nonessential |
| PA14_45940 | Nonessential | Nonessential |
| PA14_72870 | Nonessential | Nonessential |
| PA14_53970 | Nonessential | Nonessential |
| PA14_07600 | Nonessential | Nonessential |
| PA14_25970 | Nonessential | Nonessential |
| PA14_68850 | Nonessential | Nonessential |
| PA14_31760 | Nonessential | Nonessential |
| PA14_64280 | Nonessential | Nonessential |
| PA14_11690 | Nonessential | Nonessential |
| PA14_53220 | Nonessential | Nonessential |
| PA14_38110 | Nonessential | Nonessential |
| PA14_10600 | Nonessential | Nonessential |
| PA14_33500 | Nonessential | Nonessential |
| PA14_30630 | Nonessential | Nonessential |
| PA14_21910 | Nonessential | Nonessential |
| PA14_71710 | Nonessential | Nonessential |
| PA14_26460 | Nonessential | Nonessential |
| PA14_34270 | Nonessential | Nonessential |
| PA14_23790 | Nonessential | Nonessential |
| PA14_61470 | Nonessential | Nonessential |
| PA14_02830 | Nonessential | Nonessential |
| PA14_57720 | Nonessential | Nonessential |
| PA14_19510 | Nonessential | Nonessential |
| PA14_39960 | Nonessential | Nonessential |
| PA14_64310 | Nonessential | Nonessential |
| PA14_67630 | Nonessential | Nonessential |

| | | |
|---|---|---|
| PA14_06960 | Nonessential | Nonessential |
| PA14_26650 | Nonessential | Nonessential |
| PA14_19090 | Nonessential | Nonessential |
| PA14_69870 | Nonessential | Nonessential |
| PA14_64370 | Nonessential | Nonessential |
| PA14_47790 | Nonessential | Nonessential |
| PA14_44590 | Nonessential | Nonessential |
| PA14_42850 | Nonessential | Nonessential |
| PA14_47690 | Nonessential | Nonessential |
| PA14_35520 | Nonessential | Nonessential |
| PA14_68060 | Nonessential | Nonessential |
| PA14_41150 | Nonessential | Nonessential |
| PA14_62580 | Nonessential | Nonessential |
| PA14_58470 | Nonessential | Nonessential |
| PA14_37100 | Nonessential | Nonessential |
| PA14_43920 | Nonessential | Nonessential |
| PA14_25690 | Nonessential | Nonessential |
| PA14_41160 | Nonessential | Nonessential |
| PA14_08620 | Nonessential | Nonessential |
| PA14_57320 | Nonessential | Nonessential |
| PA14_30020 | Nonessential | Nonessential |
| PA14_03450 | Nonessential | Nonessential |
| PA14_68350 | Nonessential | Nonessential |
| PA14_34350 | Nonessential | Nonessential |
| PA14_71420 | Nonessential | Nonessential |
| PA14_42730 | Nonessential | Nonessential |
| PA14_31990 | Nonessential | Nonessential |
| PA14_05590 | Nonessential | Nonessential |
| PA14_09420 | Nonessential | Nonessential |
| PA14_70470 | Nonessential | Nonessential |
| PA14_51120 | Nonessential | Nonessential |
| PA14_18150 | Nonessential | Nonessential |
| PA14_47430 | Nonessential | Nonessential |
| PA14_52610 | Nonessential | Nonessential |
| PA14_10230 | Nonessential | Nonessential |
| PA14_38480 | Nonessential | Nonessential |
| PA14_10620 | Nonessential | Nonessential |
| PA14_05700 | Nonessential | Nonessential |
| PA14_24220 | Nonessential | Nonessential |
| PA14_11000 | Nonessential | Nonessential |
| PA14_25250 | Nonessential | Nonessential |
| PA14_28590 | Nonessential | Nonessential |
| PA14_54880 | Nonessential | Nonessential |
| PA14_13610 | Nonessential | Nonessential |
| PA14_36320 | Nonessential | Nonessential |
| PA14_32100 | Nonessential | Nonessential |
| PA14_33000 | Nonessential | Nonessential |
| PA14_31800 | Nonessential | Nonessential |
| PA14_03940 | Nonessential | Nonessential |
| PA14_16690 | Nonessential | Nonessential |
| PA14_30180 | Nonessential | Nonessential |
| PA14_31580 | Nonessential | Nonessential |
| PA14_68260 | Nonessential | Nonessential |
| PA14_19770 | Nonessential | Nonessential |
| PA14_10160 | Nonessential | Nonessential |
| PA14_33610 | Nonessential | Nonessential |
| PA14_05150 | Nonessential | Nonessential |
| PA14_20300 | Nonessential | Nonessential |
| PA14_31500 | Nonessential | Nonessential |
| PA14_64090 | Nonessential | Nonessential |
| PA14_43370 | Nonessential | Nonessential |
| PA14_32660 | Nonessential | Nonessential |
| PA14_25390 | Nonessential | Nonessential |
| PA14_15780 | Nonessential | Nonessential |
| PA14_06510 | Nonessential | Nonessential |
| PA14_60420 | Nonessential | Nonessential |
| PA14_43950 | Nonessential | Nonessential |
| PA14_26390 | Nonessential | Nonessential |
| PA14_68390 | Nonessential | Nonessential |
| PA14_56570 | Nonessential | Nonessential |
| PA14_37590 | Nonessential | Nonessential |
| PA14_10890 | Nonessential | Nonessential |
| PA14_12010 | Nonessential | Nonessential |
| PA14_48600 | Nonessential | Nonessential |

| | | |
|---|---|---|
| PA14_47150 | Nonessential | Nonessential |
| PA14_51980 | Nonessential | Nonessential |
| PA14_21140 | Nonessential | Nonessential |
| PA14_22910 | Nonessential | Nonessential |
| PA14_54640 | Nonessential | Nonessential |
| PA14_73320 | Nonessential | Nonessential |
| PA14_69150 | Nonessential | Nonessential |
| PA14_32220 | Nonessential | Nonessential |
| PA14_09400 | Nonessential | Nonessential |
| PA14_09460 | Nonessential | Nonessential |
| PA14_72580 | Nonessential | Nonessential |
| PA14_07850 | Nonessential | Nonessential |
| PA14_05220 | Nonessential | Nonessential |
| PA14_63850 | Nonessential | Nonessential |
| PA14_71650 | Nonessential | Nonessential |
| PA14_44760 | Nonessential | Nonessential |
| PA14_01600 | Nonessential | Nonessential |
| PA14_52670 | Nonessential | Nonessential |
| PA14_62630 | Nonessential | Nonessential |
| PA14_44500 | Nonessential | Nonessential |
| PA14_68280 | Nonessential | Nonessential |
| PA14_36345 | Nonessential | Nonessential |
| PA14_26050 | Nonessential | Nonessential |
| PA14_69130 | Nonessential | Nonessential |
| PA14_24445 | Nonessential | Nonessential |
| PA14_14860 | Nonessential | Nonessential |
| PA14_71490 | Nonessential | Nonessential |
| PA14_19110 | Nonessential | Nonessential |
| PA14_00120 | Nonessential | Nonessential |
| PA14_62160 | Nonessential | Nonessential |
| PA14_05870 | Nonessential | Nonessential |
| PA14_46490 | Nonessential | Nonessential |
| PA14_47210 | Nonessential | Nonessential |
| PA14_03430 | Nonessential | Nonessential |
| PA14_33040 | Nonessential | Nonessential |
| PA14_36660 | Nonessential | Nonessential |
| PA14_45970 | Nonessential | Nonessential |
| PA14_20420 | Nonessential | Nonessential |
| PA14_23070 | Nonessential | Nonessential |
| PA14_52270 | Nonessential | Nonessential |
| PA14_16860 | Nonessential | Nonessential |
| PA14_52910 | Nonessential | Nonessential |
| PA14_28170 | Nonessential | Nonessential |
| PA14_29110 | Nonessential | Nonessential |
| PA14_12400 | Nonessential | Nonessential |
| PA14_23470 | Nonessential | Nonessential |
| PA14_47180 | Nonessential | Nonessential |
| PA14_19910 | Nonessential | Nonessential |
| PA14_21175 | Nonessential | Nonessential |
| PA14_08350 | Nonessential | Nonessential |
| PA14_34250 | Nonessential | Nonessential |
| PA14_67500 | Nonessential | Nonessential |
| PA14_38530 | Nonessential | Nonessential |
| PA14_40980 | Nonessential | Nonessential |
| PA14_26240 | Nonessential | Nonessential |
| PA14_49250 | Nonessential | Nonessential |
| PA14_21930 | Nonessential | Nonessential |
| PA14_02790 | Nonessential | Nonessential |
| PA14_46920 | Nonessential | Nonessential |
| PA14_70370 | Nonessential | Nonessential |
| PA14_47760 | Nonessential | Nonessential |
| PA14_23620 | Nonessential | Nonessential |
| PA14_04550 | Nonessential | Nonessential |
| PA14_65040 | Nonessential | Nonessential |
| PA14_34600 | Nonessential | Nonessential |
| PA14_52800 | Nonessential | Nonessential |
| PA14_00250 | Nonessential | Nonessential |
| PA14_07740 | Nonessential | Nonessential |
| PA14_35290 | Nonessential | Nonessential |
| PA14_18600 | Nonessential | Nonessential |
| PA14_33560 | Nonessential | Nonessential |
| PA14_69610 | Nonessential | Nonessential |
| PA14_70280 | Nonessential | Nonessential |
| PA14_02840 | Nonessential | Nonessential |

| | | |
|---|---|---|
| PA14_13110 | Nonessential | Nonessential |
| PA14_43940 | Nonessential | Nonessential |
| PA14_34260 | Nonessential | Nonessential |
| PA14_34640 | Nonessential | Nonessential |
| PA14_21150 | Nonessential | Nonessential |
| PA14_19470 | Nonessential | Nonessential |
| PA14_01910 | Nonessential | Nonessential |
| PA14_18750 | Nonessential | Nonessential |
| PA14_55130 | Nonessential | Nonessential |
| PA14_13770 | Nonessential | Nonessential |
| PA14_41020 | Nonessential | Nonessential |
| PA14_41950 | Nonessential | Nonessential |
| PA14_22890 | Nonessential | Nonessential |
| PA14_56900 | Nonessential | Nonessential |
| PA14_66750 | Nonessential | Nonessential |
| PA14_69500 | Nonessential | Nonessential |
| PA14_11420 | Nonessential | Nonessential |
| PA14_34630 | Nonessential | Nonessential |
| PA14_23360 | Nonessential | Nonessential |
| PA14_53000 | Nonessential | Nonessential |
| PA14_70810 | Nonessential | Nonessential |
| PA14_48010 | Nonessential | Nonessential |
| PA14_18450 | Nonessential | Nonessential |
| PA14_44950 | Nonessential | Nonessential |
| PA14_57780 | Nonessential | Nonessential |
| PA14_23000 | Nonessential | Nonessential |
| PA14_40420 | Nonessential | Nonessential |
| PA14_32080 | Nonessential | Nonessential |
| PA14_50550 | Nonessential | Nonessential |
| PA14_39750 | Nonessential | Nonessential |
| PA14_47300 | Nonessential | Nonessential |
| PA14_60380 | Nonessential | Nonessential |
| PA14_06810 | Nonessential | Nonessential |
| PA14_66950 | Nonessential | Nonessential |
| PA14_30340 | Nonessential | Nonessential |
| PA14_19140 | Nonessential | Nonessential |
| PA14_68580 | Nonessential | Nonessential |
| PA14_71470 | Nonessential | Nonessential |
| PA14_26010 | Nonessential | Nonessential |
| PA14_52780 | Nonessential | Nonessential |
| PA14_23160 | Nonessential | Nonessential |
| PA14_09210 | Nonessential | Nonessential |
| PA14_06230 | Nonessential | Nonessential |
| PA14_12960 | Nonessential | Nonessential |
| PA14_20200 | Nonessential | Nonessential |
| PA14_54170 | Nonessential | Nonessential |
| PA14_00060 | Nonessential | Nonessential |
| PA14_07860 | Nonessential | Nonessential |
| PA14_13750 | Nonessential | Nonessential |
| PA14_51430 | Nonessential | Nonessential |
| PA14_63090 | Nonessential | Nonessential |
| PA14_20320 | Nonessential | Nonessential |
| PA14_67280 | Nonessential | Nonessential |
| PA14_52810 | Nonessential | Nonessential |
| PA14_18260 | Nonessential | Nonessential |
| PA14_68360 | Nonessential | Nonessential |
| PA14_05230 | Nonessential | Nonessential |
| PA14_16390 | Nonessential | Nonessential |
| PA14_57160 | Nonessential | Nonessential |
| PA14_19190 | Nonessential | Nonessential |
| PA14_46320 | Nonessential | Nonessential |
| PA14_44290 | Nonessential | Nonessential |
| PA14_00640 | Nonessential | Nonessential |
| PA14_70830 | Nonessential | Nonessential |
| PA14_05260 | Nonessential | Nonessential |
| PA14_43320 | Nonessential | Nonessential |
| PA14_60700 | Nonessential | Nonessential |
| PA14_01460 | Nonessential | Nonessential |
| PA14_39720 | Nonessential | Nonessential |
| PA14_44770 | Nonessential | Nonessential |
| PA14_31030 | Nonessential | Nonessential |
| PA14_52720 | Nonessential | Nonessential |
| PA14_52770 | Nonessential | Nonessential |
| PA14_67250 | Nonessential | Nonessential |

| | | |
|---|---|---|
| PA14_35970 | Nonessential | Nonessential |
| PA14_17630 | Nonessential | Nonessential |
| PA14_67270 | Nonessential | Nonessential |
| PA14_04610 | Nonessential | Nonessential |
| PA14_03700 | Nonessential | Nonessential |
| PA14_10140 | Nonessential | Nonessential |
| PA14_71940 | Nonessential | Nonessential |
| PA14_04210 | Nonessential | Nonessential |
| PA14_62150 | Nonessential | Nonessential |
| PA14_62730 | Nonessential | Nonessential |
| PA14_23850 | Nonessential | Nonessential |
| PA14_35320 | Nonessential | Nonessential |
| PA14_01900 | Nonessential | Nonessential |
| PA14_11770 | Nonessential | Nonessential |
| PA14_70940 | Nonessential | Nonessential |
| PA14_58870 | Nonessential | Nonessential |
| PA14_42080 | Nonessential | Nonessential |
| PA14_37950 | Nonessential | Nonessential |
| PA14_20400 | Nonessential | Nonessential |
| PA14_55770 | Nonessential | Nonessential |
| PA14_63640 | Nonessential | Nonessential |
| PA14_42720 | Nonessential | Nonessential |
| PA14_06290 | Nonessential | Nonessential |
| PA14_09660 | Nonessential | Nonessential |
| PA14_66570 | Nonessential | Nonessential |
| PA14_17050 | Nonessential | Nonessential |
| PA14_19500 | Nonessential | Nonessential |
| PA14_58410 | Nonessential | Nonessential |
| PA14_25980 | Nonessential | Nonessential |
| PA14_62440 | Nonessential | Nonessential |
| PA14_38490 | Nonessential | Nonessential |
| PA14_58490 | Nonessential | Nonessential |
| PA14_14940 | Nonessential | Nonessential |
| PA14_67340 | Nonessential | Nonessential |
| PA14_04090 | Nonessential | Nonessential |
| PA14_37090 | Nonessential | Nonessential |
| PA14_03130 | Nonessential | Nonessential |
| PA14_01830 | Nonessential | Nonessential |
| PA14_58450 | Nonessential | Nonessential |
| PA14_37610 | Nonessential | Nonessential |
| PA14_11790 | Nonessential | Nonessential |
| PA14_38360 | Nonessential | Nonessential |
| PA14_03930 | Nonessential | Nonessential |
| PA14_64350 | Nonessential | Nonessential |
| PA14_67050 | Nonessential | Nonessential |
| PA14_57210 | Nonessential | Nonessential |
| PA14_26485 | Nonessential | Nonessential |
| PA14_53360 | Nonessential | Nonessential |
| PA14_34420 | Nonessential | Nonessential |
| PA14_63250 | Nonessential | Nonessential |
| PA14_31700 | Nonessential | Nonessential |
| PA14_67030 | Nonessential | Nonessential |
| PA14_17010 | Nonessential | Nonessential |
| PA14_17740 | Nonessential | Nonessential |
| PA14_70860 | Nonessential | Nonessential |
| PA14_64300 | Nonessential | Nonessential |
| PA14_18380 | Nonessential | Nonessential |
| PA14_50520 | Nonessential | Nonessential |
| PA14_25270 | Nonessential | Nonessential |
| PA14_47730 | Nonessential | Nonessential |
| PA14_09240 | Nonessential | Nonessential |
| PA14_06700 | Nonessential | Nonessential |
| PA14_12990 | Nonessential | Nonessential |
| PA14_54450 | Nonessential | Nonessential |
| PA14_10850 | Nonessential | Nonessential |
| PA14_33810 | Nonessential | Nonessential |
| PA14_62000 | Nonessential | Nonessential |
| PA14_62930 | Nonessential | Nonessential |
| PA14_52820 | Nonessential | Nonessential |
| PA14_03650 | Nonessential | Nonessential |
| PA14_08560 | Nonessential | Nonessential |
| PA14_44860 | Nonessential | Nonessential |
| PA14_25210 | Nonessential | Nonessential |
| PA14_17450 | Nonessential | Nonessential |

| | | |
|---|---|---|
| PA14_53940 | Nonessential | Nonessential |
| PA14_39320 | Nonessential | Nonessential |
| PA14_12390 | Nonessential | Nonessential |
| PA14_26590 | Nonessential | Nonessential |
| PA14_64770 | Nonessential | Nonessential |
| PA14_71930 | Nonessential | Nonessential |
| PA14_10420 | Nonessential | Nonessential |
| PA14_09490 | Nonessential | Nonessential |
| PA14_35300 | Nonessential | Nonessential |
| PA14_36080 | Nonessential | Nonessential |
| PA14_58550 | Nonessential | Nonessential |
| PA14_03050 | Nonessential | Nonessential |
| PA14_63605 | Nonessential | Nonessential |
| PA14_03860 | Nonessential | Nonessential |
| PA14_15890 | Nonessential | Nonessential |
| PA14_16090 | Nonessential | Nonessential |
| PA14_06570 | Nonessential | Nonessential |
| PA14_45060 | Nonessential | Nonessential |
| PA14_26530 | Nonessential | Nonessential |
| PA14_09470 | Nonessential | Nonessential |
| PA14_24830 | Nonessential | Nonessential |
| PA14_68740 | Nonessential | Nonessential |
| PA14_38330 | Nonessential | Nonessential |
| PA14_09410 | Nonessential | Nonessential |
| PA14_72550 | Nonessential | Nonessential |
| PA14_71960 | Nonessential | Nonessential |
| PA14_68080 | Nonessential | Nonessential |
| PA14_40410 | Nonessential | Nonessential |
| PA14_13830 | Nonessential | Nonessential |
| PA14_68530 | Nonessential | Nonessential |
| PA14_15030 | Nonessential | Nonessential |
| PA14_51080 | Nonessential | Nonessential |
| PA14_45050 | Nonessential | Nonessential |
| PA14_13600 | Nonessential | Nonessential |
| PA14_38640 | Nonessential | Nonessential |
| PA14_19400 | Nonessential | Nonessential |
| PA14_42740 | Nonessential | Nonessential |
| PA14_17640 | Nonessential | Nonessential |
| PA14_18010 | Nonessential | Nonessential |
| PA14_57560 | Nonessential | Nonessential |
| PA14_62590 | Nonessential | Nonessential |
| PA14_26360 | Nonessential | Nonessential |
| PA14_30750 | Nonessential | Nonessential |
| PA14_68730 | Nonessential | Nonessential |
| PA14_73230 | Nonessential | Nonessential |
| PA14_36730 | Nonessential | Nonessential |
| PA14_62480 | Nonessential | Nonessential |
| PA14_35330 | Nonessential | Nonessential |
| PA14_10200 | Nonessential | Nonessential |
| PA14_29980 | Nonessential | Nonessential |
| PA14_33450 | Nonessential | Nonessential |
| PA14_36330 | Nonessential | Nonessential |
| PA14_10990 | Nonessential | Nonessential |
| PA14_72280 | Nonessential | Nonessential |
| PA14_44260 | Nonessential | Nonessential |
| PA14_35340 | Nonessential | Nonessential |
| PA14_21110 | Nonessential | Nonessential |
| PA14_33690 | Nonessential | Nonessential |
| PA14_65760 | Nonessential | Nonessential |
| PA14_13090 | Nonessential | Nonessential |
| PA14_17880 | Nonessential | Nonessential |
| PA14_43280 | Nonessential | Nonessential |
| PA14_68955 | Nonessential | Nonessential |
| PA14_13580 | Nonessential | Nonessential |
| PA14_39690 | Nonessential | Nonessential |
| PA14_53480 | Nonessential | Nonessential |
| PA14_23750 | Nonessential | Nonessential |
| PA14_65740 | Nonessential | Nonessential |
| PA14_70850 | Nonessential | Nonessential |
| PA14_12890 | Nonessential | Nonessential |
| PA14_52630 | Nonessential | Nonessential |
| PA14_47490 | Nonessential | Nonessential |
| PA14_21960 | Nonessential | Nonessential |
| PA14_57670 | Nonessential | Nonessential |

| | | |
|---|---|---|
| PA14_31540 | Nonessential | Nonessential |
| PA14_71560 | Nonessential | Nonessential |
| PA14_49080 | Nonessential | Nonessential |
| PA14_15790 | Nonessential | Nonessential |
| PA14_40390 | Nonessential | Nonessential |
| PA14_58630 | Nonessential | Nonessential |
| PA14_33650 | Nonessential | Nonessential |
| PA14_24730 | Nonessential | Nonessential |
| PA14_30050 | Nonessential | Nonessential |
| PA14_54930 | Nonessential | Nonessential |
| PA14_02590 | Nonessential | Nonessential |
| PA14_50540 | Nonessential | Nonessential |
| PA14_13170 | Nonessential | Nonessential |
| PA14_23010 | Nonessential | Nonessential |
| PA14_12920 | Nonessential | Nonessential |
| PA14_65770 | Nonessential | Nonessential |
| PA14_38590 | Nonessential | Nonessential |
| PA14_65480 | Nonessential | Nonessential |
| PA14_65250 | Nonessential | Nonessential |
| PA14_03960 | Nonessential | Nonessential |
| PA14_33550 | Nonessential | Nonessential |
| PA14_54670 | Nonessential | Nonessential |
| PA14_03670 | Nonessential | Nonessential |
| PA14_38320 | Nonessential | Nonessential |
| PA14_04630 | Nonessential | Nonessential |
| PA14_72620 | Nonessential | Nonessential |
| PA14_68290 | Nonessential | Nonessential |
| PA14_02680 | Nonessential | Nonessential |
| PA14_16660 | Nonessential | Nonessential |
| PA14_65110 | Nonessential | Nonessential |
| PA14_46100 | Nonessential | Nonessential |
| PA14_18410 | Nonessential | Nonessential |
| PA14_31470 | Nonessential | Nonessential |
| PA14_47190 | Nonessential | Nonessential |
| PA14_39945 | Nonessential | Nonessential |
| PA14_17930 | Nonessential | Nonessential |
| PA14_13500 | Nonessential | Nonessential |
| PA14_10530 | Nonessential | Nonessential |
| PA14_03900 | Nonessential | Nonessential |
| PA14_68770 | Nonessential | Nonessential |
| PA14_38550 | Nonessential | Nonessential |
| PA14_12020 | Nonessential | Nonessential |
| PA14_10240 | Nonessential | Nonessential |
| PA14_26890 | Nonessential | Nonessential |
| PA14_14110 | Nonessential | Nonessential |
| PA14_67890 | Nonessential | Nonessential |
| PA14_38660 | Nonessential | Nonessential |
| PA14_29920 | Nonessential | Nonessential |
| PA14_10180 | Nonessential | Nonessential |
| PA14_61250 | Nonessential | Nonessential |
| PA14_31820 | Nonessential | Nonessential |
| PA14_54660 | Nonessential | Nonessential |
| PA14_17400 | Nonessential | Nonessential |
| PA14_20670 | Nonessential | Nonessential |
| PA14_18500 | Nonessential | Nonessential |
| PA14_41470 | Nonessential | Nonessential |
| PA14_09440 | Nonessential | Nonessential |
| PA14_23270 | Nonessential | Nonessential |
| PA14_39330 | Nonessential | Nonessential |
| PA14_41830 | Nonessential | Nonessential |
| PA14_38630 | Nonessential | Nonessential |
| PA14_43180 | Nonessential | Nonessential |
| PA14_53010 | Nonessential | Nonessential |
| PA14_51330 | Nonessential | Nonessential |
| PA14_68340 | Nonessential | Nonessential |
| PA14_71630 | Nonessential | Nonessential |
| PA14_38610 | Nonessential | Nonessential |
| PA14_24950 | Nonessential | Nonessential |
| PA14_56250 | Nonessential | Nonessential |
| PA14_41840 | Nonessential | Nonessential |
| PA14_13620 | Nonessential | Nonessential |
| PA14_70160 | Nonessential | Nonessential |
| PA14_57770 | Nonessential | Nonessential |
| PA14_07890 | Nonessential | Nonessential |

| | | |
|---|---|---|
| PA14_25090 | Nonessential | Nonessential |
| PA14_30280 | Nonessential | Nonessential |
| PA14_33030 | Nonessential | Nonessential |
| PA14_70980 | Nonessential | Nonessential |
| PA14_40770 | Nonessential | Nonessential |
| PA14_36310 | Nonessential | Nonessential |
| PA14_12490 | Nonessential | Nonessential |
| PA14_67240 | Nonessential | Nonessential |
| PA14_39890 | Nonessential | Nonessential |
| PA14_31040 | Nonessential | Nonessential |
| PA14_23760 | Nonessential | Nonessential |
| PA14_53950 | Nonessential | Nonessential |
| PA14_66440 | Nonessential | Nonessential |
| PA14_26230 | Nonessential | Nonessential |
| PA14_23290 | Nonessential | Nonessential |
| PA14_33700 | Nonessential | Nonessential |
| PA14_31010 | Nonessential | Nonessential |
| PA14_62010 | Nonessential | Nonessential |
| PA14_67300 | Nonessential | Nonessential |
| PA14_64740 | Nonessential | Nonessential |
| PA14_12300 | Nonessential | Nonessential |
| PA14_04080 | Nonessential | Nonessential |
| PA14_67040 | Nonessential | Nonessential |
| PA14_43640 | Nonessential | Nonessential |
| PA14_60210 | Nonessential | Nonessential |
| PA14_01760 | Nonessential | Nonessential |
| PA14_29940 | Nonessential | Nonessential |
| PA14_22350 | Nonessential | Nonessential |
| PA14_17675 | Nonessential | Nonessential |
| PA14_64980 | Nonessential | Nonessential |
| PA14_51360 | Nonessential | Nonessential |
| PA14_47160 | Nonessential | Nonessential |
| PA14_58440 | Nonessential | Nonessential |
| PA14_57710 | Nonessential | Nonessential |
| PA14_04320 | Nonessential | Nonessential |
| PA14_56300 | Nonessential | Nonessential |
| PA14_03770 | Nonessential | Nonessential |
| PA14_36200 | Nonessential | Nonessential |
| PA14_33540 | Nonessential | Nonessential |
| PA14_10170 | Nonessential | Nonessential |
| PA14_71060 | Nonessential | Nonessential |
| PA14_26480 | Nonessential | Nonessential |
| PA14_34780 | Nonessential | Nonessential |
| PA14_35440 | Nonessential | Nonessential |
| PA14_68330 | Nonessential | Nonessential |
| PA14_02470 | Nonessential | Nonessential |
| PA14_66560 | Nonessential | Nonessential |
| PA14_33280 | Nonessential | Nonessential |
| PA14_18710 | Nonessential | Nonessential |
| PA14_43610 | Nonessential | Nonessential |
| PA14_58000 | Nonessential | Nonessential |
| PA14_63570 | Nonessential | Nonessential |
| PA14_28180 | Nonessential | Nonessential |
| PA14_64880 | Nonessential | Nonessential |
| PA14_41920 | Nonessential | Nonessential |
| PA14_27500 | Nonessential | Nonessential |
| PA14_48020 | Nonessential | Nonessential |
| PA14_70110 | Nonessential | Nonessential |
| PA14_18250 | Nonessential | Nonessential |
| PA14_23280 | Nonessential | Nonessential |
| PA14_23500 | Nonessential | Nonessential |
| PA14_45000 | Nonessential | Nonessential |
| PA14_17980 | Nonessential | Nonessential |
| PA14_04250 | Nonessential | Nonessential |
| PA14_09550 | Nonessential | Nonessential |
| PA14_03920 | Nonessential | Nonessential |
| PA14_56060 | Nonessential | Nonessential |
| PA14_71920 | Nonessential | Nonessential |
| PA14_45240 | Nonessential | Nonessential |
| PA14_11530 | Nonessential | Nonessential |
| PA14_66290 | Nonessential | Nonessential |
| PA14_47540 | Nonessential | Nonessential |
| PA14_69570 | Nonessential | Nonessential |
| PA14_14890 | Nonessential | Nonessential |

| | | |
|---|---|---|
| PA14_38140 | Nonessential | Nonessential |
| PA14_58700 | Nonessential | Nonessential |
| PA14_06670 | Nonessential | Nonessential |
| PA14_29900 | Nonessential | Nonessential |
| PA14_12940 | Nonessential | Nonessential |
| PA14_41563 | Nonessential | Nonessential |
| PA14_71280 | Nonessential | Nonessential |
| PA14_18550 | Nonessential | Nonessential |
| PA14_39910 | Nonessential | Nonessential |
| PA14_67930 | Nonessential | Nonessential |
| PA14_26510 | Nonessential | Nonessential |
| PA14_00450 | Nonessential | Nonessential |
| PA14_01620 | Nonessential | Nonessential |
| PA14_71440 | Nonessential | Nonessential |
| PA14_11430 | Nonessential | Nonessential |
| PA14_52180 | Nonessential | Nonessential |
| PA14_21340 | Nonessential | Nonessential |
| PA14_39925 | Nonessential | Nonessential |
| PA14_71020 | Nonessential | Nonessential |
| PA14_09150 | Nonessential | Nonessential |
| PA14_30010 | Nonessential | Nonessential |
| PA14_73250 | Nonessential | Nonessential |
| PA14_10790 | Nonessential | Nonessential |
| PA14_34970 | Nonessential | Nonessential |
| PA14_38580 | Nonessential | Nonessential |
| PA14_31530 | Nonessential | Nonessential |
| PA14_58030 | Nonessential | Nonessential |
| PA14_43620 | Nonessential | Nonessential |
| PA14_66260 | Nonessential | Nonessential |
| PA14_71620 | Nonessential | Nonessential |
| PA14_44070 | Nonessential | Nonessential |
| PA14_38440 | Nonessential | Nonessential |
| PA14_72960 | Nonessential | Nonessential |
| PA14_02630 | Nonessential | Nonessential |
| PA14_05790 | Nonessential | Nonessential |
| PA14_07190 | Nonessential | Nonessential |
| PA14_03680 | Nonessential | Nonessential |
| PA14_53050 | Nonessential | Nonessential |
| PA14_26210 | Nonessential | Nonessential |
| PA14_27960 | Nonessential | Nonessential |
| PA14_46950 | Nonessential | Nonessential |
| PA14_73280 | Nonessential | Nonessential |
| PA14_72170 | Nonessential | Nonessential |
| PA14_34770 | Nonessential | Nonessential |
| PA14_07170 | Nonessential | Nonessential |
| PA14_39050 | Nonessential | Nonessential |
| PA14_29850 | Nonessential | Nonessential |
| PA14_35530 | Nonessential | Nonessential |
| PA14_09280 | Nonessential | Nonessential |
| PA14_69795 | Nonessential | Nonessential |
| PA14_30190 | Nonessential | Nonessential |
| PA14_24170 | Nonessential | Nonessential |
| PA14_70040 | Nonessential | Nonessential |
| PA14_21920 | Nonessential | Nonessential |
| PA14_69990 | Nonessential | Nonessential |
| PA14_29880 | Nonessential | Nonessential |
| PA14_42690 | Nonessential | Nonessential |
| PA14_19740 | Nonessential | Nonessential |
| PA14_72000 | Nonessential | Nonessential |
| PA14_19630 | Nonessential | Nonessential |
| PA14_51350 | Nonessential | Nonessential |
| PA14_42090 | Nonessential | Nonessential |
| PA14_70100 | Nonessential | Nonessential |
| PA14_24640 | Nonessential | Nonessential |
| PA14_63800 | Nonessential | Nonessential |
| PA14_23090 | Nonessential | Nonessential |
| PA14_63080 | Nonessential | Nonessential |
| PA14_29860 | Nonessential | Nonessential |
| PA14_24190 | Nonessential | Nonessential |
| PA14_25920 | Nonessential | Nonessential |
| PA14_02810 | Nonessential | Nonessential |
| PA14_41140 | Nonessential | Nonessential |
| PA14_41540 | Nonessential | Nonessential |
| PA14_20270 | Nonessential | Nonessential |

| | | |
|---|---|---|
| PA14_25080 | Nonessential | Nonessential |
| PA14_09740 | Nonessential | Nonessential |
| PA14_37560 | Nonessential | Nonessential |
| PA14_46910 | Nonessential | Nonessential |
| PA14_68300 | Nonessential | Nonessential |
| PA14_61400 | Nonessential | Nonessential |
| PA14_19870 | Nonessential | Nonessential |
| PA14_38130 | Nonessential | Nonessential |
| PA14_44850 | Nonessential | Nonessential |
| PA14_09480 | Nonessential | Nonessential |
| PA14_71720 | Nonessential | Nonessential |
| PA14_04780 | Nonessential | Nonessential |
| PA14_61210 | Nonessential | Nonessential |
| PA14_19570 | Nonessential | Nonessential |
| PA14_14700 | Nonessential | Nonessential |
| PA14_05770 | Nonessential | Nonessential |
| PA14_08390 | Nonessential | Nonessential |
| PA14_44470 | Nonessential | Nonessential |
| PA14_71510 | Nonessential | Nonessential |
| PA14_12230 | Nonessential | Nonessential |
| PA14_65840 | Nonessential | Nonessential |
| PA14_38250 | Nonessential | Nonessential |
| PA14_50760 | Nonessential | Nonessential |
| PA14_16030 | Nonessential | Nonessential |
| PA14_10570 | Nonessential | Nonessential |
| PA14_22620 | Nonessential | Nonessential |
| PA14_19520 | Nonessential | Nonessential |
| PA14_53070 | Nonessential | Nonessential |
| PA14_72340 | Nonessential | Nonessential |
| PA14_33820 | Nonessential | Nonessential |
| PA14_00280 | Essential | Nonessential |
| PA14_67490 | Essential | Nonessential |
| PA14_49340 | Essential | Nonessential |
| PA14_34370 | Essential | Nonessential |
| PA14_31750 | Essential | Nonessential |
| PA14_67600 | Essential | Nonessential |
| PA14_44010 | Essential | Nonessential |
| PA14_60890 | Essential | Nonessential |
| PA14_58780 | Essential | Nonessential |
| PA14_44050 | Essential | Nonessential |
| PA14_49460 | Essential | Nonessential |
| PA14_70260 | Essential | Nonessential |
| PA14_44030 | Essential | Nonessential |
| PA14_66920 | Essential | Nonessential |
| PA14_44060 | Essential | Nonessential |
| PA14_49470 | Essential | Nonessential |
| PA14_04760 | Essential | Nonessential |
| PA14_11460 | Essential | Nonessential |
| PA14_11260 | Essential | Nonessential |
| PA14_50800 | Essential | Nonessential |
| PA14_73300 | Essential | Nonessential |
| PA14_11510 | Essential | Nonessential |
| PA14_44020 | Essential | Nonessential |
| PA14_17110 | Essential | Nonessential |
| PA14_11400 | Essential | Nonessential |
| PA14_11410 | Essential | Nonessential |
| PA14_70240 | Essential | Nonessential |
| PA14_56780 | Essential | Nonessential |
| PA14_08630 | Essential | Nonessential |
| PA14_07230 | Essential | Nonessential |
| PA14_25740 | Essential | Nonessential |
| PA14_29600 | Essential | Nonessential |
| PA14_73260 | Essential | Nonessential |
| PA14_44000 | Essential | Nonessential |
| PA14_11860 | Essential | Nonessential |
| PA14_39100 | Essential | Nonessential |
| PA14_07130 | Nonessential | Essential |
| PA14_00070 | Nonessential | Essential |
| PA14_70720 | Nonessential | Essential |
| PA14_70730 | Nonessential | Essential |
| PA14_12060 | Nonessential | Essential |
| PA14_66250 | Nonessential | Essential |
| PA14_15740 | Nonessential | Essential |
| PA14_17420 | Nonessential | Essential |

| | | |
|---|---|---|
| PA14_38350 | Nonessential | Essential |
| PA14_65230 | Nonessential | Essential |
| PA14_62120 | Nonessential | Essential |
| PA14_57890 | Nonessential | Essential |
| PA14_39190 | Nonessential | Essential |
| PA14_21410 | Nonessential | Essential |
| PA14_70440 | Nonessential | Essential |
| PA14_66600 | Nonessential | Essential |
| PA14_17030 | Nonessential | Essential |
| PA14_17180 | Nonessential | Essential |
| PA14_60470 | Nonessential | Essential |
| PA14_15340 | Nonessential | Essential |
| PA14_11560 | Nonessential | Essential |
| PA14_69240 | Nonessential | Essential |
| PA14_57260 | Nonessential | Essential |
| PA14_57900 | Nonessential | Essential |
| PA14_07580 | Nonessential | Essential |
| PA14_23880 | Nonessential | Essential |
| PA14_20890 | Nonessential | Essential |
| PA14_05620 | Nonessential | Essential |
| PA14_52580 | Nonessential | Essential |
| PA14_69690 | Nonessential | Essential |
| PA14_42760 | Nonessential | Essential |
| PA14_64200 | Nonessential | Essential |
| PA14_11470 | Nonessential | Essential |
| PA14_41820 | Nonessential | Essential |
| PA14_25650 | Nonessential | Essential |
| PA14_68980 | Nonessential | Essential |
| PA14_64220 | Nonessential | Essential |
| PA14_62840 | Nonessential | Essential |
| PA14_20950 | Nonessential | Essential |
| PA14_57810 | Nonessential | Essential |
| PA14_17340 | Nonessential | Essential |
| PA14_66220 | Nonessential | Essential |
| PA14_41170 | Nonessential | Essential |
| PA14_52040 | Nonessential | Essential |
| PA14_66240 | Nonessential | Essential |
| PA14_70270 | Nonessential | Essential |
| PA14_49380 | Nonessential | Essential |
| PA14_57370 | Nonessential | Essential |
| PA14_36570 | Nonessential | Essential |
| PA14_00290 | Nonessential | Essential |
| PA14_66210 | Nonessential | Essential |
| PA14_43690 | Nonessential | Essential |
| PA14_73220 | Nonessential | Essential |
| PA14_17310 | Nonessential | Essential |
| PA14_25710 | Nonessential | Essential |
| PA14_51240 | Nonessential | Essential |
| PA14_68210 | Nonessential | Essential |
| PA14_23920 | Nonessential | Essential |
| PA14_62830 | Nonessential | Essential |
| PA14_71600 | Nonessential | Essential |
| PA14_61770 | Nonessential | Essential |
| PA14_68190 | Nonessential | Essential |
| PA14_07090 | Nonessential | Essential |
| PA14_68170 | Nonessential | Essential |
| PA14_16700 | Nonessential | Essential |
| PA14_25640 | Nonessential | Essential |
| PA14_23310 | Nonessential | Essential |
