## Supplemental Data 6 for "An updated genome-scale metabolic network reconstruction of *Pseudomonas aeruginosa* PA14 to characterize mucin-driven shifts in bacterial metabolism"

| Median Exchange Fluxes for 4 Contextualized Models | | | | | | |
|---|---|---|---|---|---|---|
| | | | Median Flux from flux samples (n=500) | | | |
| | Rxn Name | Metabolite | ABTGC | MUC5AC | MUC5B | Glycans |
| Metabolites Consumed | EX_cpd00001_e | Water | -64.5 | -27.7 | -25.0 | -62.8 |
| | EX_cpd00007_e | O2 | -19.4 | -20.0 | -20.0 | -20.0 |
| | EX_cpd00021_e | Fe | 0.0 | 0.0 | 0.0 | 0.0 |
| | EX_cpd00023_e | L-Glutamate | -10.0 | -10.0 | -10.0 | -10.0 |
| | EX_cpd00027_e | D-Glucose | -8.2 | -8.0 | -10.0 | -8.3 |
| | EX_cpd00039_e | L-Lysine | -0.3 | -0.3 | -0.3 | -0.3 |
| | EX_cpd00041_e | L-Aspartate | -10.0 | -7.2 | -10.0 | -6.2 |
| | EX_cpd00051_e | L-Arginine | -10.0 | -10.0 | -10.0 | -10.0 |
| | EX_cpd00053_e | L-Glutamate | -8.8 | -10.0 | -9.4 | -9.7 |
| | EX_cpd00054_e | L-Serine | -10.0 | -3.7 | -10.0 | -10.0 |
| | EX_cpd00066_e | L-Phenylalanine | -0.4 | -0.4 | -0.4 | -0.4 |
| | EX_cpd00069_e | L-Tyrosine | -0.3 | -0.3 | -0.3 | -0.3 |
| | EX_cpd00084_e | L-Cysteine | -0.4 | -0.4 | -0.4 | -0.4 |
| | EX_cpd00107_e | L-Leucine | -1.5 | -1.4 | -1.4 | -1.5 |
| | EX_cpd00119_e | L-Histidine | -10.0 | -0.2 | -0.2 | -10.0 |
| | EX_cpd00129_e | L-Proline | -10.0 | -10.0 | -10.0 | -10.0 |
| | EX_cpd00132_e | L-Asparagine | -0.3 | -10.0 | -10.0 | -0.3 |
| | EX_cpd00156_e | L-Valine | -0.8 | -0.8 | -0.8 | -0.8 |
| | EX_cpd00161_e | L-Threonine | -7.8 | -9.9 | -7.0 | -10.0 |
| | EX_cpd00322_e | L-Isoleucine | -0.5 | -0.5 | -0.5 | -0.5 |
| Metabolites Produced | EX_cpd00011_e | CO2 | 51.1 | 52.4 | 56.9 | 52.3 |
| | EX_cpd00012_e | PPi | 0.7 | 0.6 | 0.6 | 0.7 |
| | EX_cpd00013_e | NH3 | 78.2 | 61.6 | 65.2 | 78.2 |
| | EX_cpd00029_e | Acetate | 51.8 | 38.1 | 33.8 | 50.4 |
| | EX_cpd00033_e | Glycine | 7.8 | 4.4 | 7.0 | 0.0 |
| | EX_cpd00035_e | L-Alanine | 0.0 | 0.0 | N/A | N/A |
| | EX_cpd00047_e | Formate | 9.7 | 0.0 | N/A | 9.7 |
| | EX_cpd00067_e | H+ | N/A | N/A | 79.0 | N/A |
| | EX_cpd00229_e | Glycoaldehyde | 0.0 | 0.0 | 0.0 | 0.0 |
| | EX_cpd00363_e | Ethanol | 7.3 | 9.4 | 6.5 | 9.4 |
| | EX_cpd00379_e | Glutarate | 8.4 | 8.5 | 8.6 | 8.3 |
