## Supplemental Data 7 for "An updated genome-scale metabolic network reconstruction of *Pseudomonas aeruginosa* PA14 to characterize mucin-driven shifts in bacterial metabolism"

| ABTGC | |
|---|---|
| **Rxn ID** | **Median Flux** |
| rxn01451 | -8.39 |
| rxn01387 | -35.06 |
| rxn00616 | -0.91 |
| rxn13726 | 73.58 |
| rxn00604 | 36.76 |
| rxn10902 | -36.76 |
| rxn05338 | -1.96 |
| rxn05322 | 1.96 |
| rxn05327 | 1.79 |
| rxn05324 | 1.62 |
| rxn13838 | 0.51 |
| rxn05328 | 0.04 |
| rxn05326 | 1.96 |
| rxn13850 | 1.45 |
| rxn08396 | 0.01 |
| rxn05325 | 1.96 |
| rxn05464 | 0.04 |
| rxn05339 | -1.96 |
| rxn05337 | -1.96 |
| rxn13844 | 0.01 |
| rxn05341 | -1.96 |
| rxn05336 | -1.45 |
| rxn05461 | 0.04 |
| rxn07993 | 0.69 |
| rxn05340 | -1.79 |
| rxn05342 | -1.45 |
| rxn00781 | 3.75 |
| rxn00199 | -35.06 |
| rxn08398 | 0.69 |
| rxn05289 | 0.30 |
| rxn01643 | 0.61 |
| rxn00154 | 45.44 |
| rxn00085 | -3.27 |
| rxn01740 | -0.18 |
| rxn00686 | -0.05 |
| rxn04954 | -0.27 |
| rxn01802 | -8.39 |

| MUC5AC | |
|---|---|
| **Rxn ID** | **Median Flux** |
| rxn01451 | -8.46 |
| rxn01387 | -26.18 |
| rxn00616 | -0.87 |
| rxn06493 | 0.00 |
| rxn13726 | 80.46 |
| rxn00604 | 27.81 |
| rxn10902 | -23.19 |
| rxn00248 | 9.99 |
| rxn05338 | -1.87 |
| rxn05322 | 1.87 |
| rxn05327 | 1.71 |
| rxn05324 | 1.55 |
| rxn13838 | 0.48 |
| rxn05328 | 0.04 |
| rxn05326 | 1.87 |
| rxn13850 | 1.39 |
| rxn08396 | 0.01 |
| rxn05325 | 1.87 |
| rxn05464 | 0.04 |
| rxn05339 | -1.87 |
| rxn05337 | -1.87 |
| rxn13844 | 0.01 |
| rxn05341 | -1.87 |
| rxn05336 | -1.39 |
| rxn05461 | 0.04 |
| rxn07993 | 0.65 |
| rxn05340 | -1.71 |
| rxn05342 | -1.39 |
| rxn00781 | 3.87 |
| rxn00199 | -26.18 |
| rxn08398 | 0.65 |
| rxn05289 | 0.29 |
| rxn01643 | 0.58 |
| rxn00154 | 40.29 |
| rxn01740 | -0.17 |
| rxn00686 | -0.05 |
| rxn04954 | -0.26 |

| MUC5B | |
|---|---|
| **Rxn ID** | **Median Flux** |
| rxn01451 | -8.56 |
| rxn01387 | -24.99 |
| rxn00616 | -0.87 |
| rxn13726 | 73.62 |
| rxn00604 | 27.08 |
| rxn10902 | -27.08 |
| rxn05338 | -1.87 |
| rxn05322 | 1.87 |
| rxn05327 | 1.71 |
| rxn05324 | 1.55 |
| rxn13838 | 0.48 |
| rxn05328 | 0.04 |
| rxn05326 | 1.87 |
| rxn13850 | 1.39 |
| rxn08396 | 0.01 |
| rxn05325 | 1.87 |
| rxn05464 | 0.04 |
| rxn05339 | -1.87 |
| rxn05337 | -1.87 |
| rxn13844 | 0.01 |
| rxn05341 | -1.87 |
| rxn05336 | -1.39 |
| rxn05461 | 0.04 |
| rxn07993 | 0.65 |
| rxn05340 | -1.71 |
| rxn05342 | -1.39 |
| rxn00781 | 5.38 |
| rxn00199 | -24.99 |
| rxn08398 | 0.65 |
| rxn05289 | 0.29 |
| rxn01643 | 0.58 |
| rxn00371 | 0.00 |
| rxn00154 | 43.96 |
| rxn00085 | -3.72 |
| rxn01740 | -0.17 |
| rxn00686 | -0.05 |
| rxn04954 | -0.26 |

| Glycans | |
|---|---|
| **Rxn ID** | **Median Flux** |
| rxn01451 | -8.33 |
| rxn01387 | -34.23 |
| rxn00616 | -0.91 |
| rxn06493 | 2.59 |
| rxn13726 | 74.45 |
| rxn00604 | 36.66 |
| rxn10902 | -36.66 |
| rxn05338 | -1.96 |
| rxn05322 | 1.96 |
| rxn05327 | 1.79 |
| rxn05324 | 1.62 |
| rxn13838 | 0.55 |
| rxn05326 | 1.96 |
| rxn13850 | 1.45 |
| rxn08396 | 0.05 |
| rxn05325 | 1.96 |
| rxn05339 | -1.96 |
| rxn05337 | -1.96 |
| rxn13844 | 0.05 |
| rxn05341 | -1.96 |
| rxn05336 | -1.45 |
| rxn07993 | 0.69 |
| rxn05340 | -1.79 |
| rxn05342 | -1.45 |
| rxn00781 | 3.85 |
| rxn00199 | -34.23 |
| rxn08398 | 0.69 |
| rxn05289 | 0.30 |
| rxn01643 | 0.61 |
| rxn00154 | 45.46 |
| rxn00085 | -3.54 |
| rxn01740 | -0.18 |
| rxn00686 | -0.05 |
| rxn04954 | -0.27 |
| rxn01802 | -8.33 |
| rxn02465 | 16.04 |
| rxn13895 | 9.33 |

| | |
|---|---|
| rxn02465 | 16.11 |
| rxn13895 | 9.36 |
| rxn00907 | 2.27 |
| rxn02285 | -0.34 |
| rxn03958 | 0.00 |
| rxn01302 | -0.27 |
| rxn00834 | 0.74 |
| rxn05293 | 0.00 |
| rxn08352 | 0.00 |
| rxn08756 | 0.00 |
| rxn06591 | 0.00 |
| rxn02929 | -0.34 |
| rxn02003 | -0.01 |
| rxn08087 | 0.10 |
| rxn08086 | 0.24 |
| rxn08089 | 0.33 |
| rxn08088 | 0.01 |
| rxn10158 | 5.58 |
| rxn00727 | 0.18 |
| rxn05155 | 8.81 |
| rxn08002 | 0.69 |
| rxn05349 | 1.96 |
| rxn05331 | 1.62 |
| rxn05333 | 1.79 |
| rxn08310 | 0.10 |
| rxn08311 | 0.01 |
| rxn08312 | 0.33 |
| rxn01629 | 0.00 |
| rxn00986 | -24.40 |
| rxn00675 | 24.40 |
| rPY00162 | 0.01 |
| rPY00167 | 0.03 |
| rxn01964 | 0.18 |
| rxn05237 | 0.39 |
| rxn08550 | 0.10 |
| rxn08552 | 0.33 |
| rxn02503 | 0.00 |
| rxn00256 | 35.06 |
| rJB00240 | -12.85 |

| | |
|---|---|
| rxn01802 | -8.46 |
| rxn02465 | 16.14 |
| rxn13895 | 9.38 |
| rxn00907 | 2.17 |
| rxn02285 | -0.32 |
| rxn03958 | 0.00 |
| rxn01302 | -0.26 |
| rxn00834 | 0.70 |
| rxn05293 | 0.00 |
| rxn08352 | 0.00 |
| rxn08756 | 0.00 |
| rxn06591 | 0.00 |
| rxn02929 | -0.32 |
| rxn02003 | -0.01 |
| rxn08087 | 0.10 |
| rxn08086 | 0.23 |
| rxn08089 | 0.31 |
| rxn08088 | 0.01 |
| rxn00727 | 0.17 |
| rxn05155 | 10.00 |
| rxn08002 | 0.65 |
| rxn05349 | 1.87 |
| rxn05331 | 1.55 |
| rxn05333 | 1.71 |
| rxn08310 | 0.10 |
| rxn08311 | 0.01 |
| rxn08312 | 0.31 |
| rxn01629 | 0.00 |
| rxn00986 | -26.72 |
| rxn00675 | 26.72 |
| rPY00162 | 0.01 |
| rPY00167 | 0.03 |
| rxn01964 | 0.17 |
| rxn05237 | 0.37 |
| rxn08550 | 0.10 |
| rxn08552 | 0.31 |
| rxn02503 | 0.00 |
| rxn00256 | 26.18 |
| rxn00799 | -9.99 |

| | |
|---|---|
| rxn01802 | -8.56 |
| rxn02465 | 15.21 |
| rxn13895 | 9.33 |
| rxn00907 | 2.17 |
| rxn02285 | -0.32 |
| rxn00935 | 1.63 |
| rxn03958 | 0.00 |
| rxn01302 | -0.26 |
| rxn00834 | 0.70 |
| rxn05293 | 0.00 |
| rxn08352 | 0.00 |
| rxn08756 | 0.00 |
| rxn06591 | 0.00 |
| rxn02929 | -0.32 |
| rxn02003 | -0.01 |
| rxn08087 | 0.10 |
| rxn08086 | 0.23 |
| rxn08089 | 0.31 |
| rxn08088 | 0.01 |
| rxn10158 | -44.70 |
| rxn00727 | 0.17 |
| rxn05155 | 9.39 |
| rxn08002 | 0.65 |
| rxn05349 | 1.87 |
| rxn05331 | 1.55 |
| rxn05333 | 1.71 |
| rxn08310 | 0.10 |
| rxn08311 | 0.01 |
| rxn08312 | 0.31 |
| rxn01629 | 0.00 |
| rxn00986 | -69.31 |
| rxn00675 | 69.31 |
| rPY00162 | 0.01 |
| rPY00167 | 0.03 |
| rxn01964 | 0.17 |
| rxn05237 | 0.37 |
| rxn08550 | 0.10 |
| rxn08552 | 0.31 |
| rxn02503 | 0.00 |

| | |
|---|---|
| rxn00907 | 2.27 |
| rxn02285 | -0.34 |
| rxn03958 | 0.00 |
| rxn01302 | -0.27 |
| rxn00834 | 0.74 |
| rxn05293 | 0.00 |
| rxn08352 | 0.00 |
| rxn08756 | 0.00 |
| rxn06591 | 0.00 |
| rxn02929 | -0.34 |
| rxn02003 | -0.01 |
| rxn08087 | 0.10 |
| rxn08086 | 0.24 |
| rxn08089 | 0.33 |
| rxn08088 | 0.01 |
| rxn10158 | 2.71 |
| rxn00727 | 0.18 |
| rxn05155 | 9.74 |
| rxn08002 | 0.69 |
| rxn05349 | 1.96 |
| rxn05331 | 1.62 |
| rxn05333 | 1.79 |
| rxn08310 | 0.10 |
| rxn08311 | 0.01 |
| rxn08312 | 0.33 |
| rxn01629 | 0.00 |
| rxn00986 | -23.96 |
| rxn00675 | 23.96 |
| rPY00162 | 0.01 |
| rPY00167 | 0.03 |
| rxn01964 | 0.18 |
| rxn05237 | 0.39 |
| rxn08550 | 0.10 |
| rxn08552 | 0.33 |
| rxn02503 | 0.00 |
| rxn00256 | 34.23 |
| rJB00240 | -12.78 |
| rxn03004 | 1.14 |
| rxn01644 | 0.34 |

| | |
|---|---|
| rxn03004 | 1.14 |
| rxn01644 | 0.34 |
| rxn05231 | 0.05 |
| rxn06076 | 0.10 |
| rxn05233 | 0.10 |
| rxn06075 | 0.05 |
| rxn05561 | 7.62 |
| rxn00394 | 7.09 |
| rxn00285 | -61.68 |
| rxn05334 | 1.96 |
| rxn05329 | 1.96 |
| rxn05330 | 1.96 |
| rxn13843 | 0.01 |
| rxn13840 | 1.41 |
| rxn13846 | 1.45 |
| rxn01332 | 0.18 |
| rxn00973 | -35.06 |
| rxn05307 | 10.00 |
| rxn05300 | 7.83 |
| rPY00184 | 0.00 |
| rxn08229 | 0.03 |
| rxn08230 | 0.01 |
| rPY00183 | 0.01 |
| rxn08231 | 0.00 |
| rxn08232 | 0.04 |
| rPY00172 | 0.01 |
| rPY00175 | 0.02 |
| rxn00175 | -51.29 |
| rxn03884 | 5.47 |
| rxn00214 | 0.00 |
| rxn01639 | 9.73 |
| rxn05147 | 2.56 |
| rxn00299 | 0.00 |
| rxn05215 | 0.00 |
| rxn00615 | 0.09 |
| rxn00533 | 13.26 |
| rxn05332 | 0.04 |
| rxn07980 | 0.69 |
| rxn05462 | 0.04 |

| | |
|---|---|
| rJB00238 | 3.29 |
| rxn03004 | 1.08 |
| rxn01644 | 0.32 |
| rxn05231 | 0.05 |
| rxn06076 | 0.10 |
| rxn05233 | 0.10 |
| rxn06075 | 0.05 |
| rxn10154 | -3.29 |
| rxn00394 | 9.03 |
| rxn00285 | -46.19 |
| rxn05334 | 1.87 |
| rxn05329 | 1.87 |
| rxn05330 | 1.87 |
| rxn13843 | 0.01 |
| rxn13840 | 1.35 |
| rxn13846 | 1.39 |
| rxn01332 | 0.17 |
| rxn00973 | -26.18 |
| rxn05307 | 3.71 |
| rxn05300 | 9.92 |
| rPY00184 | 0.00 |
| rxn08229 | 0.02 |
| rxn08230 | 0.01 |
| rPY00183 | 0.01 |
| rxn08231 | 0.00 |
| rxn08232 | 0.04 |
| rxn00342 | 9.65 |
| rxn06600 | 0.00 |
| rxn06377 | 0.00 |
| rPY00172 | 0.01 |
| rPY00175 | 0.02 |
| rxn00175 | -37.68 |
| rxn03884 | 5.50 |
| rxn00214 | 0.00 |
| rxn05147 | 8.02 |
| rxn00299 | 0.00 |
| rxn05215 | 0.00 |
| rxn00615 | 0.08 |
| rxn00533 | 12.65 |

| | |
|---|---|
| rxn00256 | 24.99 |
| rxn00799 | -1.63 |
| rxn03004 | 1.08 |
| rxn01644 | 0.32 |
| rxn05231 | 0.05 |
| rxn06076 | 0.10 |
| rxn05233 | 0.10 |
| rxn06075 | 0.05 |
| rxn05561 | 44.70 |
| rxn00394 | 7.24 |
| rxn00285 | 0.32 |
| rxn05334 | 1.87 |
| rxn05329 | 1.87 |
| rxn05330 | 1.87 |
| rxn13843 | 0.01 |
| rxn13840 | 1.35 |
| rxn13846 | 1.39 |
| rxn01332 | 0.17 |
| rxn00973 | -24.99 |
| rxn05307 | 10.00 |
| rxn05300 | 7.04 |
| rPY00184 | 0.00 |
| rxn08229 | 0.02 |
| rxn08230 | 0.01 |
| rPY00183 | 0.01 |
| rxn08231 | 0.00 |
| rxn08232 | 0.04 |
| rxn00342 | 9.67 |
| rPY00172 | 0.01 |
| rPY00175 | 0.02 |
| rxn00175 | -33.37 |
| rxn03884 | 7.20 |
| rxn00214 | 0.00 |
| rxn05147 | 2.44 |
| rxn00299 | 0.00 |
| rxn00615 | 0.08 |
| rxn00533 | 12.65 |
| rxn05332 | 0.04 |
| rxn07980 | 0.65 |

| | |
|---|---|
| rxn05231 | 0.05 |
| rxn06076 | 0.10 |
| rxn05233 | 0.10 |
| rxn06075 | 0.05 |
| rxn05561 | 8.36 |
| rxn00394 | 7.04 |
| rxn00285 | -58.03 |
| rxn05334 | 1.96 |
| rxn05329 | 1.96 |
| rxn05330 | 1.96 |
| rxn13843 | 0.05 |
| rxn13840 | 1.45 |
| rxn13846 | 1.45 |
| rxn01332 | 0.18 |
| rxn00973 | -34.23 |
| rxn05307 | 10.00 |
| rxn05300 | 10.00 |
| rPY00184 | 0.00 |
| rxn08229 | 0.03 |
| rxn08230 | 0.01 |
| rPY00183 | 0.01 |
| rxn08231 | 0.00 |
| rxn08232 | 0.04 |
| rxn06600 | 2.59 |
| rxn06377 | 2.59 |
| rPY00172 | 0.01 |
| rPY00175 | 0.02 |
| rxn00175 | -49.96 |
| rxn03884 | 5.59 |
| rxn00214 | 0.00 |
| rxn01639 | 9.71 |
| rxn05147 | 2.56 |
| rxn00299 | 0.00 |
| rxn00615 | 0.09 |
| rxn00533 | 13.26 |
| rxn07980 | 0.69 |
| rxn08309 | 0.24 |
| rPY00163 | 0.01 |
| rPY00168 | 0.03 |

| | |
|---|---|
| rxn08309 | 0.24 |
| rPY00163 | 0.01 |
| rPY00168 | 0.03 |
| rxn05582 | -7.79 |
| rxn13836 | 5.47 |
| rxn00148 | 38.06 |
| rxn13689 | 77.79 |
| rxn00405 | 1.81 |
| rxn02213 | 0.18 |
| rxn00101 | 7.09 |
| rxn13870 | 0.00 |
| rxn00867 | 9.73 |
| rxn05347 | 1.96 |
| rxn13107 | 0.00 |
| rxn05466 | -78.19 |
| rxn00283 | 0.68 |
| rxn08103 | -38.89 |
| rxn00692 | -2.59 |
| rxn05229 | -13.26 |
| rxn13804 | -7.29 |
| rxn10042 | 151.63 |
| rxn03919 | 0.01 |
| rxn02303 | 0.00 |
| rxn01029 | 1.81 |
| rxn00853 | 1.81 |
| rxn05508 | 0.31 |
| rxn00777 | -1.67 |
| rxn01520 | 0.05 |
| rxn01303 | 0.27 |
| rxn00953 | 5.78 |
| rxn00950 | 6.07 |
| rxn02302 | 0.27 |
| rxn01018 | 1.11 |
| rxn00141 | 0.00 |
| rxn11268 | 0.66 |
| rxn00126 | 0.03 |
| rxn01200 | 0.75 |
| rxn00785 | 0.93 |
| rxn01100 | -3.75 |

| | |
|---|---|
| rxn05332 | 0.04 |
| rxn07980 | 0.65 |
| rxn05462 | 0.04 |
| rxn08309 | 0.23 |
| rPY00163 | 0.01 |
| rPY00168 | 0.03 |
| rxn05209 | -19.55 |
| rxn05582 | -4.35 |
| rxn00148 | 36.11 |
| rxn13689 | 80.00 |
| rxn00470 | 1.73 |
| rxn02213 | 0.17 |
| rxn00101 | 9.03 |
| rxn13870 | 0.00 |
| rxn05347 | 1.87 |
| rxn13107 | 0.00 |
| rxn05466 | -61.65 |
| rxn00283 | 0.65 |
| rxn08103 | -40.00 |
| rxn00692 | -2.47 |
| rxn05229 | -12.65 |
| rxn13804 | -9.36 |
| rxn10042 | 158.93 |
| rxn03919 | 0.01 |
| rxn02303 | 0.00 |
| rxn05508 | 10.00 |
| rxn00777 | -1.60 |
| rxn01520 | 0.05 |
| rxn01303 | 0.26 |
| rxn00950 | 0.26 |
| rxn02302 | 0.26 |
| rxn01018 | 1.06 |
| rxn00141 | 0.00 |
| rxn11268 | 0.63 |
| rxn00126 | 0.03 |
| rxn01200 | 0.71 |
| rxn00785 | 0.89 |
| rxn01100 | -3.87 |
| rxn02504 | 0.00 |

| | |
|---|---|
| rxn05462 | 0.04 |
| rxn08309 | 0.23 |
| rPY00163 | 0.01 |
| rPY00168 | 0.03 |
| rxn05209 | -12.66 |
| rxn05582 | -6.99 |
| rxn13836 | 7.20 |
| rxn00148 | 37.63 |
| rxn13689 | 79.99 |
| rxn00405 | 1.73 |
| rxn02213 | 0.17 |
| rxn00101 | 7.24 |
| rxn13870 | 0.00 |
| rxn05347 | 1.87 |
| rxn13107 | 0.00 |
| rxn05466 | -65.25 |
| rxn00283 | 0.65 |
| rxn08103 | -40.00 |
| rxn00692 | -2.47 |
| rxn05229 | -12.65 |
| rxn13804 | -6.46 |
| rxn10042 | 152.09 |
| rxn03919 | 0.01 |
| rxn02303 | 0.00 |
| rxn01029 | 1.73 |
| rxn00853 | 1.73 |
| rxn05508 | 10.00 |
| rxn00777 | -1.60 |
| rxn01520 | 0.05 |
| rxn01303 | 0.26 |
| rxn00953 | 5.77 |
| rxn00950 | 6.07 |
| rxn02302 | 0.26 |
| rxn01018 | 1.06 |
| rxn00141 | 0.00 |
| rxn11268 | 0.63 |
| rxn00126 | 0.03 |
| rxn01200 | 0.71 |
| rxn00785 | 0.89 |

| | |
|---|---|
| rxn05582 | 0.00 |
| rxn13836 | 5.59 |
| rxn00148 | 37.38 |
| rxn13689 | 79.99 |
| rxn00405 | 1.81 |
| rxn02213 | 0.18 |
| rxn00101 | 7.04 |
| rxn13870 | 0.00 |
| rxn00867 | 9.71 |
| rxn05347 | 1.96 |
| rxn13107 | 0.00 |
| rxn05466 | -78.24 |
| rxn00283 | 0.68 |
| rxn08103 | -40.00 |
| rxn00692 | 0.00 |
| rxn05229 | -13.26 |
| rxn13804 | -9.38 |
| rxn10042 | 152.89 |
| rxn03919 | 0.01 |
| rxn02303 | 0.00 |
| rxn01029 | 1.81 |
| rxn00853 | 1.81 |
| rxn05508 | 0.31 |
| rxn00777 | -1.67 |
| rxn01520 | 0.05 |
| rxn01303 | 0.27 |
| rxn00953 | 5.76 |
| rxn00950 | 6.05 |
| rxn02302 | 0.27 |
| rxn01018 | 1.11 |
| rxn00141 | 0.00 |
| rxn11268 | 0.66 |
| rxn00126 | 0.03 |
| rxn01200 | 0.75 |
| rxn00785 | 0.93 |
| rxn01100 | -3.85 |
| rxn02504 | 0.00 |
| rxn01116 | -1.68 |
| rxn00791 | 0.18 |

| | |
|---|---|
| rxn02504 | 0.00 |
| rxn01116 | -1.68 |
| rxn00791 | 0.18 |
| rxn02507 | 0.18 |
| rxn12880 | 0.00 |
| rxn05221 | 10.00 |
| rxn00670 | 24.40 |
| rxn00985 | -24.40 |
| rxn01637 | 16.11 |
| rxn00467 | -9.36 |
| rxn00541 | 7.29 |
| rxn00337 | 0.61 |
| rxn02937 | 1.14 |
| rxn03147 | 1.14 |
| rxn09566 | 38.89 |
| rxn01973 | 0.34 |
| rxn05241 | 0.50 |
| rxn05240 | 1.47 |
| rxn05242 | 0.81 |
| rxn05460 | 0.04 |
| rxn05346 | 1.96 |
| rxn05348 | 1.79 |
| rxn05345 | 1.45 |
| rxn13851 | 0.01 |
| rxn05350 | 1.96 |
| rxn05343 | 1.96 |
| rxn05344 | 1.45 |
| rxn01255 | 0.18 |
| rxn02167 | 8.39 |
| rxn01257 | 0.00 |
| rxn13884 | 0.08 |
| rxn01211 | 2.27 |
| rxn05687 | 10.00 |
| rxn10182 | -10.00 |
| rxn00693 | 0.27 |
| rxn00134 | 0.00 |
| rxn03901 | 0.34 |
| rxn00988 | 62.02 |
| rxn00290 | -62.02 |

| | |
|---|---|
| rxn01116 | -1.60 |
| rxn00791 | 0.17 |
| rxn02507 | 0.17 |
| rxn12880 | 0.00 |
| rxn05221 | 10.00 |
| rxn00670 | 26.72 |
| rxn00985 | -26.72 |
| rxn01637 | 16.14 |
| rxn00467 | -9.38 |
| rxn00541 | 9.36 |
| rxn00337 | 0.58 |
| rxn02937 | 1.08 |
| rxn03147 | 1.08 |
| rxn09566 | 40.00 |
| rxn01973 | 0.32 |
| rxn05241 | 0.47 |
| rxn05240 | 1.41 |
| rxn05242 | 0.78 |
| rxn05460 | 0.04 |
| rxn05346 | 1.87 |
| rxn05348 | 1.71 |
| rxn05345 | 1.39 |
| rxn13851 | 0.01 |
| rxn05350 | 1.87 |
| rxn05343 | 1.87 |
| rxn05344 | 1.39 |
| rxn01255 | 0.17 |
| rxn02167 | 8.46 |
| rxn01257 | 0.00 |
| rxn13884 | 0.08 |
| rxn01211 | 2.17 |
| rxn05687 | 10.00 |
| rxn10182 | -10.00 |
| rxn00693 | 0.26 |
| rxn00134 | 0.00 |
| rxn03901 | 0.32 |
| rxn00988 | 46.51 |
| rxn00290 | -46.51 |
| rxn00213 | 0.02 |

| | |
|---|---|
| rxn01100 | -5.38 |
| rxn02504 | 0.00 |
| rxn01116 | -1.60 |
| rxn00791 | 0.17 |
| rxn02507 | 0.17 |
| rxn12880 | 0.00 |
| rxn05221 | 10.00 |
| rxn00670 | 69.31 |
| rxn00985 | -69.31 |
| rxn01637 | 15.21 |
| rxn00467 | -9.33 |
| rxn00541 | 6.46 |
| rxn00337 | 0.58 |
| rxn02937 | 1.08 |
| rxn03147 | 1.08 |
| rxn09566 | 40.00 |
| rxn01973 | 0.32 |
| rxn05241 | 0.47 |
| rxn05240 | 1.41 |
| rxn05242 | 0.78 |
| rxn05460 | 0.04 |
| rxn05346 | 1.87 |
| rxn05348 | 1.71 |
| rxn05345 | 1.39 |
| rxn13851 | 0.01 |
| rxn05350 | 1.87 |
| rxn05343 | 1.87 |
| rxn05344 | 1.39 |
| rxn01255 | 0.17 |
| rxn02167 | 8.56 |
| rxn01257 | 0.00 |
| rxn13884 | 0.08 |
| rxn01211 | 2.17 |
| rxn05687 | 10.00 |
| rxn10182 | -10.00 |
| rxn00693 | 0.26 |
| rxn00134 | 0.00 |
| rxn03901 | 0.32 |
| rxn00213 | 0.02 |

| | |
|---|---|
| rxn02507 | 0.18 |
| rxn12880 | 0.00 |
| rxn05221 | 10.00 |
| rxn00670 | 23.96 |
| rxn00985 | -23.96 |
| rxn01637 | 16.04 |
| rxn00467 | -9.33 |
| rxn00541 | 9.38 |
| rxn00337 | 0.61 |
| rxn02937 | 1.14 |
| rxn03147 | 1.14 |
| rxn09566 | 40.00 |
| rxn01973 | 0.34 |
| rxn05241 | 0.50 |
| rxn05240 | 1.47 |
| rxn05242 | 0.81 |
| rxn05346 | 1.96 |
| rxn05348 | 1.79 |
| rxn05345 | 1.45 |
| rxn13851 | 0.05 |
| rxn05350 | 1.96 |
| rxn05343 | 1.96 |
| rxn05344 | 1.45 |
| rxn01255 | 0.18 |
| rxn02167 | 8.33 |
| rxn01257 | 0.00 |
| rxn13884 | 0.08 |
| rxn01211 | 2.27 |
| rxn05687 | 10.00 |
| rxn10182 | -10.00 |
| rxn00693 | 0.27 |
| rxn00134 | 0.00 |
| rxn03901 | 0.34 |
| rxn00988 | 58.37 |
| rxn00290 | -58.37 |
| rxn00213 | 0.02 |
| rxn13643 | 0.00 |
| rJB00278 | 5.59 |
| rxn01275 | 5.59 |

| | |
|---|---|
| rxn00213 | 0.02 |
| rxn13643 | 0.00 |
| rJB00278 | 5.47 |
| rxn01275 | 5.47 |
| rxn05571 | 5.47 |
| rxn09111 | 0.09 |
| rxn09112 | 0.04 |
| rxn09113 | 0.00 |
| rxn09114 | 0.11 |
| rxn03136 | 1.14 |
| rxn00800 | 0.40 |
| rxn05559 | -9.73 |
| rxn01333 | -0.75 |
| rxn00191 | 37.21 |
| rxn00710 | 1.11 |
| rxn03841 | 0.00 |
| rxn05465 | 13.26 |
| rxn05459 | 0.04 |
| rxn02405 | 0.01 |
| rxn13880 | 0.08 |
| rxn05306 | 0.42 |
| rxn08335 | 1.11 |
| rxn00790 | 1.14 |
| rxn01603 | 0.00 |
| rxn02508 | 0.18 |
| rxn06937 | 0.00 |
| rxn00364 | 0.10 |
| rxn01219 | -0.10 |
| rxn02476 | 0.18 |
| rxn12892 | 0.00 |
| rxn05003 | 0.00 |
| rxn09502 | 2.53 |
| rxn00216 | 0.02 |
| rxn01477 | 5.47 |
| rxn13873 | 0.08 |
| rxn03167 | 0.00 |
| rxn03511 | 0.01 |
| rxn00374 | 9.73 |
| rxn01465 | -1.11 |

| | |
|---|---|
| rxn13643 | 0.00 |
| rxn09111 | 0.09 |
| rxn09112 | 0.04 |
| rxn09113 | 0.00 |
| rxn09114 | 0.11 |
| rxn03136 | 1.08 |
| rxn00800 | 0.38 |
| rxn05559 | 0.00 |
| rxn01333 | -0.72 |
| rxn00191 | 37.49 |
| rxn00710 | 1.06 |
| rxn03841 | 0.00 |
| rxn05465 | 12.65 |
| rxn05459 | 0.04 |
| rxn02405 | 0.01 |
| rxn13880 | 0.08 |
| rxn05306 | 0.40 |
| rxn08335 | 1.06 |
| rxn00790 | 1.08 |
| rxn01603 | 0.00 |
| rxn02508 | 0.17 |
| rxn06937 | 0.00 |
| rxn00364 | 0.10 |
| rxn01219 | -0.10 |
| rxn02476 | 0.17 |
| rxn12892 | 0.00 |
| rxn05003 | 0.00 |
| rxn01476 | 5.50 |
| rxn09502 | 2.42 |
| rxn00216 | 5.52 |
| rxn01477 | 5.50 |
| rxn13873 | 0.08 |
| rxn03167 | 0.00 |
| rxn03511 | 0.01 |
| rxn01465 | -1.06 |
| rxn03910 | 0.00 |
| rxn03907 | 0.00 |
| rxn00459 | 3.87 |
| rxn02331 | 0.01 |

| | |
|---|---|
| rxn13643 | 0.00 |
| rJB00278 | 7.20 |
| rxn01275 | 7.20 |
| rxn05571 | 7.20 |
| rxn09111 | 0.09 |
| rxn09112 | 0.04 |
| rxn09113 | 0.00 |
| rxn09114 | 0.11 |
| rxn03136 | 1.08 |
| rxn00800 | 0.38 |
| rxn01333 | -0.72 |
| rxn00191 | 37.28 |
| rxn00710 | 1.06 |
| rxn03841 | 0.00 |
| rxn05465 | 12.65 |
| rxn05459 | 0.04 |
| rxn02405 | 0.01 |
| rxn13880 | 0.08 |
| rxn05306 | 0.40 |
| rxn08335 | 1.06 |
| rxn00790 | 1.08 |
| rxn01603 | 0.00 |
| rxn02508 | 0.17 |
| rxn06937 | 0.00 |
| rxn00364 | 0.10 |
| rxn01219 | -0.10 |
| rxn02476 | 0.17 |
| rxn12892 | 0.00 |
| rxn05003 | 0.00 |
| rxn09502 | 2.42 |
| rxn00216 | 0.02 |
| rxn01477 | 7.20 |
| rxn13873 | 0.08 |
| rxn03167 | 0.00 |
| rxn03511 | 0.01 |
| rxn01465 | -1.06 |
| rxn03910 | 0.00 |
| rxn03907 | 0.00 |
| rxn00459 | 5.38 |

| | |
|---|---|
| rxn05571 | 5.59 |
| rxn09111 | 0.09 |
| rxn09112 | 0.04 |
| rxn09113 | 0.00 |
| rxn09114 | 0.11 |
| rxn03136 | 1.14 |
| rxn00800 | 0.40 |
| rxn05559 | -9.71 |
| rxn01333 | -0.75 |
| rxn00191 | 37.07 |
| rxn00710 | 1.11 |
| rxn03841 | 0.00 |
| rxn05465 | 13.26 |
| rxn02405 | 0.01 |
| rxn13880 | 0.08 |
| rxn05306 | 0.42 |
| rxn08335 | 1.11 |
| rxn00790 | 1.14 |
| rxn01603 | 0.00 |
| rxn02508 | 0.18 |
| rxn06937 | 0.00 |
| rxn00364 | 0.10 |
| rxn01219 | -0.10 |
| rxn02476 | 0.18 |
| rxn12892 | 0.00 |
| rxn05003 | 0.00 |
| rxn09502 | 2.53 |
| rxn00216 | 0.02 |
| rxn01477 | 5.59 |
| rxn13873 | 0.08 |
| rxn03167 | 0.00 |
| rxn03511 | 0.01 |
| rxn00374 | 9.71 |
| rxn01465 | -1.11 |
| rxn03910 | 0.00 |
| rxn03907 | 0.00 |
| rxn00459 | 3.85 |
| rxn02331 | 0.01 |
| rxn00412 | 0.73 |

| | |
|---|---|
| rxn03910 | 0.00 |
| rxn03907 | 0.00 |
| rxn00459 | 3.75 |
| rxn02331 | 0.01 |
| rxn00412 | 0.73 |
| rxn13874 | 0.08 |
| rxn13882 | 0.17 |
| rxn13881 | 0.17 |
| rxn00119 | 1.21 |
| rxn01517 | -0.05 |
| rxn03087 | -0.34 |
| rxn03031 | 0.34 |
| rxn08549 | 0.26 |
| rxn10204 | 0.04 |
| rxn01127 | -0.05 |
| rxn00251 | 36.13 |
| rxn03084 | 1.14 |
| rxn05301 | 0.30 |
| rxn00917 | 0.74 |
| rxn00409 | 0.00 |
| rxn01512 | 0.01 |
| rxn00237 | 0.40 |
| rxn00117 | 1.60 |
| rxn10199 | 0.17 |
| rxn13044 | 0.00 |
| rxn00001 | 41.37 |
| rxn13826 | 0.00 |
| rxn01466 | 0.00 |
| rxn03909 | 0.00 |
| rPY00173 | 0.01 |
| rxn09104 | 0.09 |
| rxn09105 | 0.04 |
| rPY00176 | 0.02 |
| rxn09106 | 0.00 |
| rxn09107 | 0.11 |
| rxn10963 | 0.00 |
| rxn01636 | -16.11 |
| rxn13883 | 0.17 |
| rxn00851 | 0.34 |

| | |
|---|---|
| rxn00412 | 4.83 |
| rxn00410 | -3.99 |
| rxn13874 | 0.08 |
| rxn13882 | 0.16 |
| rxn13881 | 0.16 |
| rxn00119 | 1.15 |
| rxn01517 | -0.05 |
| rxn03087 | -0.32 |
| rxn03031 | 0.32 |
| rxn08549 | 0.24 |
| rxn10204 | 0.04 |
| rxn01127 | -0.05 |
| rxn00251 | 33.67 |
| rxn03084 | 1.08 |
| rxn05301 | 0.29 |
| rxn00917 | 0.70 |
| rxn00409 | 0.00 |
| rxn01512 | 0.01 |
| rxn00237 | 0.38 |
| rxn00117 | 1.53 |
| rxn10199 | 0.16 |
| rxn13044 | 0.00 |
| rxn00001 | 39.18 |
| rxn13826 | 0.00 |
| rxn01466 | 0.00 |
| rxn03909 | 0.00 |
| rPY00173 | 0.01 |
| rxn09104 | 0.09 |
| rxn09105 | 0.04 |
| rPY00176 | 0.02 |
| rxn09106 | 0.00 |
| rxn09107 | 0.11 |
| rxn10963 | 0.00 |
| rxn01636 | -16.14 |
| rxn13883 | 0.16 |
| rxn00851 | 0.32 |
| rxn02286 | 0.32 |
| rxn03408 | 0.32 |
| rxn02008 | 0.32 |

| | |
|---|---|
| rxn02331 | 0.01 |
| rxn00412 | 0.70 |
| rxn13874 | 0.08 |
| rxn13882 | 0.16 |
| rxn13881 | 0.16 |
| rxn00119 | 1.15 |
| rxn01517 | -0.05 |
| rxn03087 | -0.32 |
| rxn03031 | 0.32 |
| rxn08549 | 0.24 |
| rxn10204 | 0.04 |
| rxn01127 | -0.05 |
| rxn00251 | 34.37 |
| rxn03084 | 1.08 |
| rxn05301 | 0.29 |
| rxn00917 | 0.70 |
| rxn00409 | 0.00 |
| rxn01512 | 0.01 |
| rxn00237 | 0.38 |
| rxn00117 | 1.53 |
| rxn10199 | 0.16 |
| rxn13044 | 0.00 |
| rxn00001 | 40.71 |
| rxn13826 | 0.00 |
| rxn01466 | 0.00 |
| rxn03909 | 0.00 |
| rPY00173 | 0.01 |
| rxn09104 | 0.09 |
| rxn09105 | 0.04 |
| rPY00176 | 0.02 |
| rxn09106 | 0.00 |
| rxn09107 | 0.11 |
| rxn10963 | 0.00 |
| rxn01636 | -15.21 |
| rxn13883 | 0.16 |
| rxn00851 | 0.32 |
| rxn02286 | 0.32 |
| rxn03408 | 0.32 |
| rxn02008 | 0.32 |

| | |
|---|---|
| rxn13874 | 0.08 |
| rxn13882 | 0.17 |
| rxn13881 | 0.17 |
| rxn00119 | 1.21 |
| rxn01517 | -0.05 |
| rxn03087 | -0.34 |
| rxn03031 | 0.34 |
| rxn08549 | 0.26 |
| rxn08551 | 0.04 |
| rxn01127 | -0.05 |
| rxn00251 | 34.15 |
| rxn03084 | 1.14 |
| rxn05301 | 0.30 |
| rxn00917 | 0.74 |
| rxn00409 | 0.00 |
| rxn01512 | 0.01 |
| rxn00237 | 0.40 |
| rxn00117 | 1.60 |
| rxn10199 | 0.17 |
| rxn13044 | 0.00 |
| rxn00001 | 40.68 |
| rxn13826 | 0.00 |
| rxn01466 | 0.00 |
| rxn03909 | 0.00 |
| rPY00173 | 0.01 |
| rxn09104 | 0.09 |
| rxn09105 | 0.04 |
| rPY00176 | 0.02 |
| rxn09106 | 0.00 |
| rxn09107 | 0.11 |
| rxn10963 | 0.00 |
| rxn01636 | -16.04 |
| rxn13883 | 0.17 |
| rxn00851 | 0.34 |
| rxn02286 | 0.34 |
| rxn03408 | 0.34 |
| rxn02008 | 0.34 |
| rxn03904 | 0.34 |
| rxn03164 | 0.34 |

| | |
|---|---|
| rxn02286 | 0.34 |
| rxn03408 | 0.34 |
| rxn02008 | 0.34 |
| rxn03904 | 0.34 |
| rxn03164 | 0.34 |
| rxn02011 | 0.34 |
| rxn03917 | 0.01 |
| rxn00461 | 0.34 |
| rxn01117 | -0.01 |
| rxn02404 | 0.01 |
| rxn09037 | 0.00 |
| rxn05028 | 0.00 |
| rxn00014 | 77.79 |
| rxn09674 | 0.34 |
| rxn12218 | 0.00 |
| rxn00193 | 0.34 |
| rxn03908 | 0.00 |
| rxn00770 | 2.42 |
| rPY00164 | 0.00 |
| rxn09208 | 0.15 |
| rxn09209 | 0.06 |
| rPY00169 | 0.01 |
| rxn09210 | 0.01 |
| rxn09211 | 0.22 |
| rxn02380 | 2.53 |
| rxn00747 | 0.91 |
| rxn01485 | -0.85 |
| rxn02201 | 0.00 |
| rxn00832 | -1.14 |
| rxn03137 | 1.14 |
| rxn02895 | 1.14 |
| rxn00838 | 0.40 |
| rPY00165 | 0.00 |
| rxn09200 | 0.15 |
| rxn09201 | 0.06 |
| rPY00170 | 0.01 |
| rxn09202 | 0.01 |
| rxn09203 | 0.22 |
| rxn00260 | 7.09 |

| | |
|---|---|
| rxn03904 | 0.32 |
| rxn03164 | 0.32 |
| rxn02011 | 0.32 |
| rxn03917 | 0.01 |
| rxn00461 | 0.32 |
| rxn01117 | -0.01 |
| rxn02404 | 0.01 |
| rxn09037 | 0.00 |
| rxn05028 | 0.00 |
| rxn00014 | 80.00 |
| rxn09674 | 0.33 |
| rxn12218 | 0.00 |
| rxn00193 | 0.32 |
| rxn03908 | 0.00 |
| rxn00770 | 2.31 |
| rPY00164 | 0.00 |
| rxn09208 | 0.15 |
| rxn09209 | 0.06 |
| rPY00169 | 0.01 |
| rxn09210 | 0.01 |
| rxn09211 | 0.21 |
| rxn02380 | 2.42 |
| rxn00747 | 0.87 |
| rxn01485 | -0.81 |
| rxn02201 | 0.00 |
| rxn00832 | -1.08 |
| rxn03137 | 1.08 |
| rxn02895 | 1.08 |
| rxn00838 | 0.38 |
| rPY00165 | 0.00 |
| rxn09200 | 0.15 |
| rxn09201 | 0.06 |
| rPY00170 | 0.01 |
| rxn09202 | 0.01 |
| rxn09203 | 0.21 |
| rxn13878 | 0.00 |
| rxn13879 | 0.00 |
| rxn03916 | 0.01 |
| rxn03918 | 0.01 |

| | |
|---|---|
| rxn03904 | 0.32 |
| rxn03164 | 0.32 |
| rxn02011 | 0.32 |
| rxn03917 | 0.01 |
| rxn00461 | 0.32 |
| rxn01117 | -0.01 |
| rxn02404 | 0.01 |
| rxn09037 | 0.00 |
| rxn05028 | 0.00 |
| rxn00014 | 79.99 |
| rxn09674 | 0.33 |
| rxn12218 | 0.00 |
| rxn00193 | 0.32 |
| rxn03908 | 0.00 |
| rxn00770 | 2.31 |
| rPY00164 | 0.00 |
| rxn09208 | 0.15 |
| rxn09209 | 0.06 |
| rPY00169 | 0.01 |
| rxn09210 | 0.01 |
| rxn09211 | 0.21 |
| rxn02380 | 2.42 |
| rxn00747 | 0.87 |
| rxn01485 | -0.81 |
| rxn02201 | 0.00 |
| rxn00832 | -1.08 |
| rxn03137 | 1.08 |
| rxn02895 | 1.08 |
| rxn00838 | 0.38 |
| rPY00165 | 0.00 |
| rxn09200 | 0.15 |
| rxn09201 | 0.06 |
| rPY00170 | 0.01 |
| rxn09202 | 0.01 |
| rxn09203 | 0.21 |
| rxn00260 | 6.35 |
| rxn13878 | 0.00 |
| rxn13879 | 0.00 |
| rxn03916 | 0.01 |

| | |
|---|---|
| rxn02011 | 0.34 |
| rxn03917 | 0.01 |
| rxn00461 | 0.34 |
| rxn01117 | -0.01 |
| rxn02404 | 0.01 |
| rxn09037 | 0.00 |
| rxn05028 | 0.00 |
| rxn00014 | 79.99 |
| rxn09674 | 0.34 |
| rxn12218 | 0.00 |
| rxn00193 | 0.34 |
| rxn03908 | 0.00 |
| rxn00770 | 2.42 |
| rPY00164 | 0.00 |
| rxn09208 | 0.15 |
| rxn09209 | 0.06 |
| rPY00169 | 0.01 |
| rxn09210 | 0.01 |
| rxn09211 | 0.22 |
| rxn02380 | 2.53 |
| rxn00747 | 0.91 |
| rxn01485 | -0.85 |
| rxn02201 | 0.00 |
| rxn00832 | -1.14 |
| rxn03137 | 1.14 |
| rxn02895 | 1.14 |
| rxn00838 | 0.40 |
| rPY00165 | 0.00 |
| rxn09200 | 0.15 |
| rxn09201 | 0.06 |
| rPY00170 | 0.01 |
| rxn09202 | 0.01 |
| rxn09203 | 0.22 |
| rxn00260 | 3.69 |
| rxn13878 | 0.00 |
| rxn13879 | 0.00 |
| rxn03916 | 0.01 |
| rxn03918 | 0.01 |
| rxn13867 | 0.00 |

| | |
|---|---|
| rxn13878 | 0.00 |
| rxn13879 | 0.00 |
| rxn03916 | 0.01 |
| rxn03918 | 0.01 |
| rxn13867 | 0.00 |
| rxn13866 | 0.00 |
| rxn13869 | 0.00 |
| rPY00222 | 0.00 |
| rPY00221 | 0.00 |
| rxn02288 | 0.00 |
| rxn02212 | 0.18 |
| rxn01739 | 0.18 |
| rxn12879 | 0.00 |
| rxn01642 | 9.73 |
| rxn05299 | 10.00 |
| rxn02085 | -9.73 |
| rxn01106 | -3.75 |
| rxn01997 | 0.01 |
| rxn01675 | 0.01 |
| rxn02000 | 0.01 |
| rxn10131 | 10.00 |
| rxn00114 | 1.11 |
| rxn13108 | 0.00 |
| rxn00029 | 0.00 |
| rxn02264 | 0.00 |
| rxn00060 | 0.00 |
| rxn01974 | 0.34 |
| rxn00704 | -0.02 |
| rxn01917 | -16.11 |
| rxn01362 | -1.11 |
| rxn01509 | -0.10 |
| rxn00239 | 0.10 |
| rxn00966 | 0.00 |
| rxn05034 | 0.00 |
| rxn10053 | 14.40 |
| rxn00347 | -1.54 |
| rxn05217 | 10.00 |
| rxn05297 | 10.00 |
| rxn00124 | -38.89 |

| | |
|---|---|
| rxn13867 | 0.00 |
| rxn13866 | 0.00 |
| rxn13869 | 0.00 |
| rPY00222 | 0.00 |
| rPY00221 | 0.00 |
| rxn02288 | 0.00 |
| rxn02212 | 0.17 |
| rxn01739 | 0.17 |
| rxn12879 | 0.00 |
| rxn05299 | 0.25 |
| rxn01106 | -3.87 |
| rxn01997 | 0.01 |
| rxn01675 | 0.01 |
| rxn02000 | 0.01 |
| rxn10131 | 10.00 |
| rxn00114 | 1.06 |
| rxn13108 | 0.00 |
| rxn00029 | 0.00 |
| rxn02264 | 0.00 |
| rxn00060 | 0.00 |
| rxn01974 | 0.32 |
| rxn00704 | -0.02 |
| rxn01917 | -16.14 |
| rxn01362 | -1.06 |
| rxn01509 | -0.10 |
| rxn00239 | 0.10 |
| rxn00966 | 0.00 |
| rxn05034 | 0.00 |
| rxn10053 | 13.73 |
| rxn00347 | 8.43 |
| rxn10151 | 7.18 |
| rxn05297 | 10.00 |
| rxn00124 | -40.00 |
| rxn01807 | 40.00 |
| rxn00555 | 0.81 |
| rxn03638 | 0.81 |
| rxn00293 | 0.81 |
| rxn05467 | -52.42 |
| rxn05319 | 27.75 |

| | |
|---|---|
| rxn03918 | 0.01 |
| rxn13867 | 0.00 |
| rxn13866 | 0.00 |
| rxn13869 | 0.00 |
| rPY00222 | 0.00 |
| rPY00221 | 0.00 |
| rxn02288 | 0.00 |
| rxn02212 | 0.17 |
| rxn01739 | 0.17 |
| rxn12879 | 0.00 |
| rxn05299 | 0.25 |
| rxn01106 | -5.38 |
| rxn01997 | 0.01 |
| rxn01675 | 0.01 |
| rxn02000 | 0.01 |
| rxn10131 | 10.00 |
| rxn00114 | 1.06 |
| rxn13108 | 0.00 |
| rxn00029 | 0.00 |
| rxn02264 | 0.00 |
| rxn00060 | 0.00 |
| rxn01974 | 0.32 |
| rxn00704 | -0.02 |
| rxn01917 | -15.21 |
| rxn01362 | -1.06 |
| rxn01509 | -0.10 |
| rxn00239 | 0.10 |
| rxn00966 | 0.00 |
| rxn05034 | 0.00 |
| rxn10053 | 13.73 |
| rxn00347 | 0.00 |
| rxn05217 | 10.00 |
| rxn05297 | 10.00 |
| rxn00124 | -40.00 |
| rxn01807 | 40.00 |
| rxn00555 | 0.81 |
| rxn03638 | 0.81 |
| rxn00293 | 0.81 |
| rxn05467 | -56.86 |

| | |
|---|---|
| rxn13866 | 0.00 |
| rxn13869 | 0.00 |
| rPY00222 | 0.00 |
| rPY00221 | 0.00 |
| rxn02288 | 0.00 |
| rxn02212 | 0.18 |
| rxn01739 | 0.18 |
| rxn12879 | 0.00 |
| rxn01642 | 9.71 |
| rxn05299 | 10.00 |
| rxn02085 | -9.71 |
| rxn01106 | -3.85 |
| rxn01997 | 0.01 |
| rxn01675 | 0.01 |
| rxn02000 | 0.01 |
| rxn10131 | 10.00 |
| rxn00114 | 1.11 |
| rxn13108 | 0.00 |
| rxn00029 | 0.00 |
| rxn02264 | 0.00 |
| rxn00060 | 0.00 |
| rxn01974 | 0.34 |
| rxn00704 | -0.02 |
| rxn01917 | -16.04 |
| rxn01362 | -1.11 |
| rxn01509 | -0.10 |
| rxn00239 | 0.10 |
| rxn00966 | 0.00 |
| rxn05034 | 0.00 |
| rxn10053 | 14.40 |
| rxn00347 | -1.54 |
| rxn05217 | 6.21 |
| rxn05297 | 10.00 |
| rxn00124 | -40.00 |
| rxn01807 | 40.00 |
| rxn00555 | 0.85 |
| rxn03638 | 0.85 |
| rxn00293 | 0.85 |
| rxn05467 | -52.31 |

| | |
|---|---|
| rxn01807 | 38.89 |
| rxn00555 | 0.85 |
| rxn03638 | 0.85 |
| rxn00293 | 0.85 |
| rxn05467 | -51.11 |
| rxn05319 | 64.53 |
| rxn05468 | 19.45 |
| rPY00182 | 0.00 |
| rxn13868 | 0.00 |
| rxn13871 | 0.00 |
| rPY00180 | 0.00 |
| rPY00166 | 0.00 |
| rPY00171 | 0.01 |
| rPY00174 | 0.00 |
| rPY00177 | 0.01 |
| rxn05488 | -51.76 |
| rxn09683 | -7.29 |
| EX_cpd00029_e | 51.76 |
| EX_cpd00011_e | 51.11 |
| EX_cpd00027_e | -8.19 |
| EX_cpd00021_e | 0.00 |
| EX_cpd00033_e | 7.79 |
| EX_cpd00001_e | -64.53 |
| EX_cpd00035_e | 0.00 |
| EX_cpd00051_e | -10.00 |
| EX_cpd00132_e | -0.31 |
| EX_cpd00041_e | -10.00 |
| EX_cpd00084_e | -0.39 |
| EX_cpd00023_e | -10.00 |
| EX_cpd00053_e | -8.81 |
| EX_cpd00119_e | -10.00 |
| EX_cpd00322_e | -0.50 |
| EX_cpd00107_e | -1.47 |
| EX_cpd00039_e | -0.34 |
| EX_cpd00066_e | -0.42 |
| EX_cpd00129_e | -10.00 |
| EX_cpd00054_e | -10.00 |
| EX_cpd00161_e | -7.83 |
| EX_cpd00069_e | -0.30 |

| | |
|---|---|
| rxn05468 | 20.00 |
| rPY00182 | 0.00 |
| rxn13868 | 0.00 |
| rxn13871 | 0.00 |
| rPY00180 | 0.00 |
| rPY00166 | 0.00 |
| rPY00171 | 0.01 |
| rPY00174 | 0.00 |
| rPY00177 | 0.00 |
| rxn05488 | -38.14 |
| rxn09683 | -9.36 |
| EX_cpd00029_e | 38.14 |
| EX_cpd00011_e | 52.42 |
| EX_cpd00027_e | -8.02 |
| EX_cpd00021_e | 0.00 |
| EX_cpd00033_e | 4.35 |
| EX_cpd00001_e | -27.75 |
| EX_cpd00035_e | 0.00 |
| EX_cpd00051_e | -10.00 |
| EX_cpd00132_e | -10.00 |
| EX_cpd00041_e | -7.18 |
| EX_cpd00084_e | -0.37 |
| EX_cpd00023_e | -10.00 |
| EX_cpd00053_e | -10.00 |
| EX_cpd00119_e | -0.25 |
| EX_cpd00322_e | -0.47 |
| EX_cpd00107_e | -1.41 |
| EX_cpd00039_e | -0.33 |
| EX_cpd00066_e | -0.40 |
| EX_cpd00129_e | -10.00 |
| EX_cpd00054_e | -3.71 |
| EX_cpd00161_e | -9.92 |
| EX_cpd00069_e | -0.29 |
| EX_cpd00156_e | -0.78 |
| EX_cpd00013_e | 61.65 |
| EX_cpd00007_e | -20.00 |
| EX_cpd00012_e | 0.63 |
| rxn00695 | 0.00 |
| rxn13837 | -8.46 |

| | |
|---|---|
| rxn05319 | 25.04 |
| rxn05468 | 20.00 |
| rPY00182 | 0.00 |
| rxn13868 | 0.00 |
| rxn13871 | 0.00 |
| rPY00180 | 0.00 |
| rPY00166 | 0.00 |
| rPY00171 | 0.01 |
| rPY00174 | 0.00 |
| rPY00177 | 0.00 |
| rxn05488 | -33.82 |
| rxn09683 | -6.46 |
| EX_cpd00029_e | 33.82 |
| EX_cpd00011_e | 56.86 |
| EX_cpd00027_e | -10.00 |
| EX_cpd00021_e | 0.00 |
| EX_cpd00033_e | 6.99 |
| EX_cpd00067_e | 79.05 |
| EX_cpd00001_e | -25.04 |
| EX_cpd00051_e | -10.00 |
| EX_cpd00132_e | -10.00 |
| EX_cpd00041_e | -10.00 |
| EX_cpd00084_e | -0.37 |
| EX_cpd00023_e | -10.00 |
| EX_cpd00053_e | -9.39 |
| EX_cpd00119_e | -0.25 |
| EX_cpd00322_e | -0.47 |
| EX_cpd00107_e | -1.41 |
| EX_cpd00039_e | -0.33 |
| EX_cpd00066_e | -0.40 |
| EX_cpd00129_e | -10.00 |
| EX_cpd00054_e | -10.00 |
| EX_cpd00161_e | -7.04 |
| EX_cpd00069_e | -0.29 |
| EX_cpd00156_e | -0.78 |
| EX_cpd00013_e | 65.25 |
| EX_cpd00007_e | -20.00 |
| EX_cpd00012_e | 0.63 |
| rxn00695 | 0.00 |

| | |
|---|---|
| rxn05319 | 62.78 |
| rxn05468 | 20.00 |
| rPY00182 | 0.00 |
| rxn13868 | 0.00 |
| rxn13871 | 0.00 |
| rPY00180 | 0.00 |
| rPY00166 | 0.00 |
| rPY00171 | 0.01 |
| rPY00174 | 0.00 |
| rPY00177 | 0.01 |
| rxn05488 | -50.42 |
| rxn09683 | -9.38 |
| EX_cpd00029_e | 50.42 |
| EX_cpd00011_e | 52.31 |
| EX_cpd00027_e | -8.35 |
| EX_cpd00021_e | 0.00 |
| EX_cpd00033_e | 0.00 |
| EX_cpd00001_e | -62.78 |
| EX_cpd00051_e | -10.00 |
| EX_cpd00132_e | -0.31 |
| EX_cpd00041_e | -6.21 |
| EX_cpd00084_e | -0.39 |
| EX_cpd00023_e | -10.00 |
| EX_cpd00053_e | -9.74 |
| EX_cpd00119_e | -10.00 |
| EX_cpd00322_e | -0.50 |
| EX_cpd00107_e | -1.47 |
| EX_cpd00039_e | -0.34 |
| EX_cpd00066_e | -0.42 |
| EX_cpd00129_e | -10.00 |
| EX_cpd00054_e | -10.00 |
| EX_cpd00161_e | -10.00 |
| EX_cpd00069_e | -0.30 |
| EX_cpd00156_e | -0.81 |
| EX_cpd00013_e | 78.24 |
| EX_cpd00007_e | -20.00 |
| EX_cpd00012_e | 0.66 |
| rxn00695 | 0.00 |
| rxn13837 | -8.33 |

| | |
|---|---|
| EX_cpd00156_e | -0.81 |
| EX_cpd00013_e | 78.19 |
| EX_cpd00007_e | -19.45 |
| EX_cpd00012_e | 0.66 |
| rxn00695 | 0.00 |
| rxn13837 | -8.39 |
| rxn01730 | -8.39 |
| rxn09680 | 0.00 |
| rxn13893 | 0.00 |
| ATPM | 8.39 |
| PA14_Biomass | 3.18 |
| rxn00182 | 77.11 |
| EX_cpd00047_e | 9.73 |
| EX_cpd00229_e | 0.00 |
| EX_cpd00363_e | 7.29 |
| EX_cpd00379_e | 8.39 |
| rxn00178 | 8.39 |
| rxn12751 | 0.17 |
| protein_rxn | 3.18 |
| rna_rxn | 3.18 |
| dna_rxn | 3.18 |
| lipid_rxn | 3.18 |
| SK_cpd11416_c | 3.18 |

| | |
|---|---|
| rxn01730 | -8.46 |
| rxn09680 | 0.00 |
| rxn13893 | 0.00 |
| ATPM | 8.39 |
| PA14_Biomass | 3.04 |
| rxn00182 | 66.50 |
| EX_cpd00047_e | 0.00 |
| EX_cpd00229_e | 0.00 |
| EX_cpd00363_e | 9.36 |
| EX_cpd00379_e | 8.46 |
| rxn00178 | 8.46 |
| rxn12751 | 0.16 |
| protein_rxn | 3.04 |
| rna_rxn | 3.04 |
| dna_rxn | 3.04 |
| lipid_rxn | 3.04 |
| SK_cpd11416_c | 3.04 |

| | |
|---|---|
| rxn13837 | -8.56 |
| rxn01730 | -8.56 |
| rxn09680 | 0.00 |
| rxn13893 | 0.00 |
| ATPM | 8.39 |
| PA14_Biomass | 3.04 |
| rxn00182 | 71.37 |
| EX_cpd00229_e | 0.00 |
| EX_cpd00363_e | 6.46 |
| EX_cpd00379_e | 8.56 |
| rxn00178 | 8.56 |
| rxn12751 | 0.16 |
| protein_rxn | 3.04 |
| rna_rxn | 3.04 |
| dna_rxn | 3.04 |
| lipid_rxn | 3.04 |
| SK_cpd11416_c | 3.04 |

| | |
|---|---|
| rxn01730 | -8.33 |
| rxn09680 | 0.00 |
| rxn13893 | 0.00 |
| ATPM | 8.39 |
| PA14_Biomass | 3.18 |
| rxn00182 | 77.17 |
| EX_cpd00047_e | 9.71 |
| EX_cpd00229_e | 0.00 |
| EX_cpd00363_e | 9.38 |
| EX_cpd00379_e | 8.33 |
| rxn00178 | 8.33 |
| rxn12751 | 0.17 |
| protein_rxn | 3.18 |
| rna_rxn | 3.18 |
| dna_rxn | 3.18 |
| lipid_rxn | 3.18 |
| SK_cpd11416_c | 3.18 |
